# Mapping planktonic communities: a network approach to assess the role of scale and centrality on their diversity and composition

**DOI:** 10.1101/2024.01.11.574175

**Authors:** David Cunillera-Montcusí, Mia Bengtsson, Blake Matthews, Christian Preiler, Zsófia Horváth, Csaba F. Vad, Robert Ptacnik

## Abstract

The distribution of habitats across a landscape and their centrality gradient are key elements defining the effective pathways of dispersal, and thus of metacommunity assembly. Understanding how centrality shapes diversity patterns is essential for predicting the impact of future landscape changes on diversity. While alpine lakes have been extensively studied, often considering the fluvial network as a potential landscape, small planktonic communities have frequently been overlooked as potential dispersers due to their assumed ubiquity. In this study, we investigate the diversity patterns of alpine lake planktonic communities along lake networks constructed at different scales, ranging from 6.5 to 650 km and the fluvial network. We sampled 55 lakes in the northern Alps (16S, 18S, phytoplankton and zooplankton) and calculated several diversity metrics (alpha, beta diversity and LCBD) and multivariate analysis. We then constructed several networks responding to different scales, determined their centrality gradients, and finally explored their relationship with the diversity of each planktonic group. We expected that a groups’ diversity would relate differently across scales based on body size, but the outcomes were varied. Bacterioplankton and zooplankton diversity were both affected across scales higher than 100 km, whereas phytoplankton appeared completely unrelated to centrality. Nonetheless, we could observe that when significant, the relationships between diversity and centrality were shared among organisms. These findings not only underscore that planktonic organisms are influenced by landscape configurations larger than the fluvial system but also emphasise the critical role of dispersal for these groups and the scales at which it impacts metacommunity assembly.

**Significance statement:** While dispersal is widely recognized as a key driver of assembly, some groups and systems remain insufficiently explored to fully grasp the impact of landscape and dispersal on their assembly. Planktonic communities have traditionally been considered ubiquitous and detached from regional-level structure, primarily due to their small size, leading to the notion that “everything is everywhere”. Additionally, alpine lake communities have traditionally been perceived as solely connected through fluvial systems. In this study, we challenge these notions by demonstrating how planktonic communities are indeed influenced by the relative positioning of lakes in the landscape, with significant impacts occurring at larger scales, spanning hundreds of kilometres. However, not all planktonic groups responded uniformly to the analysed factors, emphasizing the marked differences among groups and the diverging drivers shaping planktonic metacommunities.

## Introduction

Metacommunity theory has contributed to a better comprehension of how the spatial arrangement of communities influences community assembly (Leibold et al. 2004, Leibold and Chase 2018, Thompson et al. 2020). The increasing recognition and quantification of spatial features has greatly advanced the field (Dale 2017, Gounand et al. 2018, Chubaty et al. 2020) since the first seminal works on metacommunity ecology (MacArthur and Wilson 1963, Hanski 1999). We are now able to quantify the scales at which the spatial arrangement plays a stronger role (Dray et al. 2006, Borcard et al. 2011), how these scales might be affected by directionality (Larsen et al. 2012, Altermatt 2013, Horváth et al. 2016), how the spatial arrangement impacts community patterns (Carrara et al. 2014, Epele et al. 2021, Borthagaray et al. 2023b) and how the relative position within a network defines diversity patterns (Economo and Keitt 2010, Altermatt et al. 2011, Henriques-Silva et al. 2019). All these approximations have been mostly focused on disentangling how space, in a broad sense, was shaping community assembly and its relative importance with respect to other variables (Peres-Neto et al. 2006, Gilbert and Bennett 2010, Bo et al. 2020, Almeida-Gomes et al. 2020). However, more detailed landscape analyses have unravelled the inherent complexity of spatial features, especially when considering the capacity of organisms to interact with, or to disperse across, a landscape (Newman 2003, Borthagaray et al. 2015a, Erős and Lowe 2019). Such organism-landscape interactions might greatly determine the scale at which spatial processes influence diversity and how species interact with one another (Ishiyama et al. 2014, Borthagaray et al. 2015b, Cunillera-Montcusí et al. 2021). Ultimately, dispersal, together with landscape structure, are the elements defining the canvas through which metacommunities assemble (Leibold and Chase 2018, Thompson et al. 2020, Borthagaray et al. 2023b).

Most previous works on aquatic metacommunity assembly have been focused on how organisms are able to actively disperse between habitats (i.e. invertebrates; (Ishiyama et al. 2014, Perez Rocha et al. 2018, Burgazzi et al. 2020), if they move shorter or longer distances within aquatic habitats (i.e. fish, molluscs; (Kappes and Haase 2012, Radinger and Wolter 2014, Herrera-Pérez et al. 2019), or disperse passively relying on different vectors, with adaptations that enables them to cross the “dry ocean” (Dodson 1992, Incagnone et al. 2015). However, the role of spatial processes in smaller microscopic organisms has received less attention until recently (Reche et al. 2005, Declerck et al. 2013, Winter et al. 2013, Jamoneau et al. 2018, Daffonchio et al. 2019, Choudoir and DeAngelis 2022). For these groups we are beginning to unfold how “everything might not be actually everywhere” (Whitfield 2005, Fontaneto et al. 2008, O’Malley 2008, Custer et al. 2022, Barbour et al. 2023).

When focusing on landscape structure, lotic systems (riverscapes) and small lentic system (pondscapes) have been mostly the grounds where studies focusing on spatial structure have been carried out in limnology (Almeida-Gomes et al. 2020, Heino et al. 2021, Hill et al. 2021). These systems often have specific dynamics that constrain dispersal of most of their inhabitants, because the strong directional component pushes organisms downstream (Seymour et al. 2015, Tonkin et al. 2018b), or its environmental features force organisms to move between habitats in order to complete their life cycles (Cunillera-Montcusí et al. 2020, Hill et al. 2021, Onandia et al. 2021), or in case of temporary lentic or lotic systems, drying mandates organisms to leave to more perennial neighbouring systems or resist drought in the soil (Martins et al. 2019, Cañedo-Argüelles et al. 2020, Florencio et al. 2020). In this sense, bigger and deeper lentic systems such as alpine lakes have remained understudied (Almeida-Gomes et al. 2020). In general, for these systems, landscape has been primarily considered as a determinant of local lake conditions linked to catchment geo-physical characteristics (e.g. bedrock, water residence time, land use; (Füreder et al. 2006, Schindler 2009, Klaus et al. 2021). However, the spatial position of lakes in the landscape can also define the connectivity between its communities and its neighbouring aquatic habitats. Being connected through waterways and/or closer to each other should foster more similar communities due to higher probability of exchange of individuals between lakes (Leibold and Norberg 2004, Heino et al. 2015). However, while the relevance of catchment geo-physical characteristics is clearly acknowledged as a diversity driver, the role of connectivity is often debated, especially for planktonic and microscopic aquatic organisms (Leibold and Norberg 2004, Winter et al. 2013, Custer et al. 2022).

The influence that landscape position might have on organisms may largely vary depending on the scale at which the landscape is considered (Declerck et al. 2011, Chase and Knight 2013, Borthagaray et al. 2015a, Morán-Ordóñez et al. 2015). Ultimately, a landscape is defined a priori based on the objectives, metrics or sites under study (Newman 2003, Turner 2005, Turner and Gardner 2015). However, how we simplify the landscape in our analyses can affect our inference about the assembly of focal communities (Baguette and Van Dyck 2007, Newman 2019). For example, if we set the boundaries of our landscape too large or too small, we might fail to capture the spatial pattern that affects community assembly. Thus, the challenge for our analyses lies in how to define an appropriate scale to consider the landscape (Baguette et al. 2013, Borthagaray et al. 2015a, Cunillera-Montcusí et al. 2021). In the case of organisms that can be transported along long distances due to dominant winds (Cáceres and Soluk 2002, Csabai and Boda 2005, Pinceel et al. 2016) or inside/attached to other organisms such as aquatic microorganisms or small crustaceans (Vanschoenwinkel et al. 2008, Coughlan et al. 2017, Lovas-Kiss et al. 2020), the scales at which landscape features might be relevant could extend several hundreds of kilometres (Green et al. 2002, Lovas-Kiss et al. 2019) or be only linked to the fluvial network connecting lakes if only dispersal by drift exist (Doretto et al. 2020). Hence, if we want to assess the role of landscape centrality for lake planktonic communities it is key to consider different scales granting the possibility to quantify how landscape arrangement might impact diversity patterns for these different taxonomic groups (Borthagaray et al. 2014, Cunillera-Montcusí et al. 2021).

Network theory can contribute to a better representation of landscape structures and understanding about the potential effect of different landscapes on biodiversity patterns (Urban and Keitt 2001, Dale 2017). By creating networks based on previously defined landscapes, we are able to calculate metrics that quantify the relative position of habitats in these networks, and directly test how relevant these are for biodiversity (e.g. centrality gradient; (Estrada and Bodin 2008, Economo and Keitt 2010, Horváth et al. 2019, Borthagaray et al. 2020). Furthermore, congruent patterns between different organisms and the scales at which they relate with centrality can inform us about the extent at which landscape is impacting their assembly patterns (Hill et al. 2017, Sarremejane et al. 2017, Schmera et al. 2018). Such congruence might be related with body size (Ortiz et al. 2023), that will markedly differ across planktonic groups, from bacterioplankton to zooplankton (Figure 1 H1). Smaller organisms (e.g. bacterioplankton and protists) might be related with larger scales not necessarily linked to fluvial connections whereas zooplankton might be related to smaller scales or to the fluvial network (Michels et al. 2001, Braghin et al. 2015, Chaudhary et al. 2022, Custer et al. 2022). However, even though the relationship of organisms with centrality might differ across scales, we could expect that the direction of these relationships would be preserved across organism groups (Figure 1 H2). Thus, on the one hand, we could hypothesise that communities located at more central positions would present greater species richness diversity and lower beta diversity as central communities could be easily reached by most of the regional pool of species (Mouquet and Loreau 2003, Ai et al. 2013, Grainger and Gilbert 2016, Custer et al. 2022). On the other hand, more isolated communities would contain lower alpha diversity and higher beta diversity (Chase and Shulman 2009, Tonkin et al. 2016b, Bender et al. 2017, Custer et al. 2022). This would reflect a concomitant change in composition, with central communities being more similar among them while greater differentiation would be observed in more isolated ones, potentially affecting communities contribution to beta diversity (Legendre and De Cáceres 2013). These general patterns may foster a stronger mismatch with environment when it relates with species richness but not against composition as environmental characteristics may fail to explain the total number of species in central sites (i.e. higher due to mass effects) but may perform better in explaining community dissimilarity (i.e. higher homogeneity). At which scales centrality will drive this mismatch will shed light on the impact of landscape on these processes and the equilibria between main assembly drivers (Thompson et al. 2020, Suzuki and Economo 2021).

**Figure 1:**
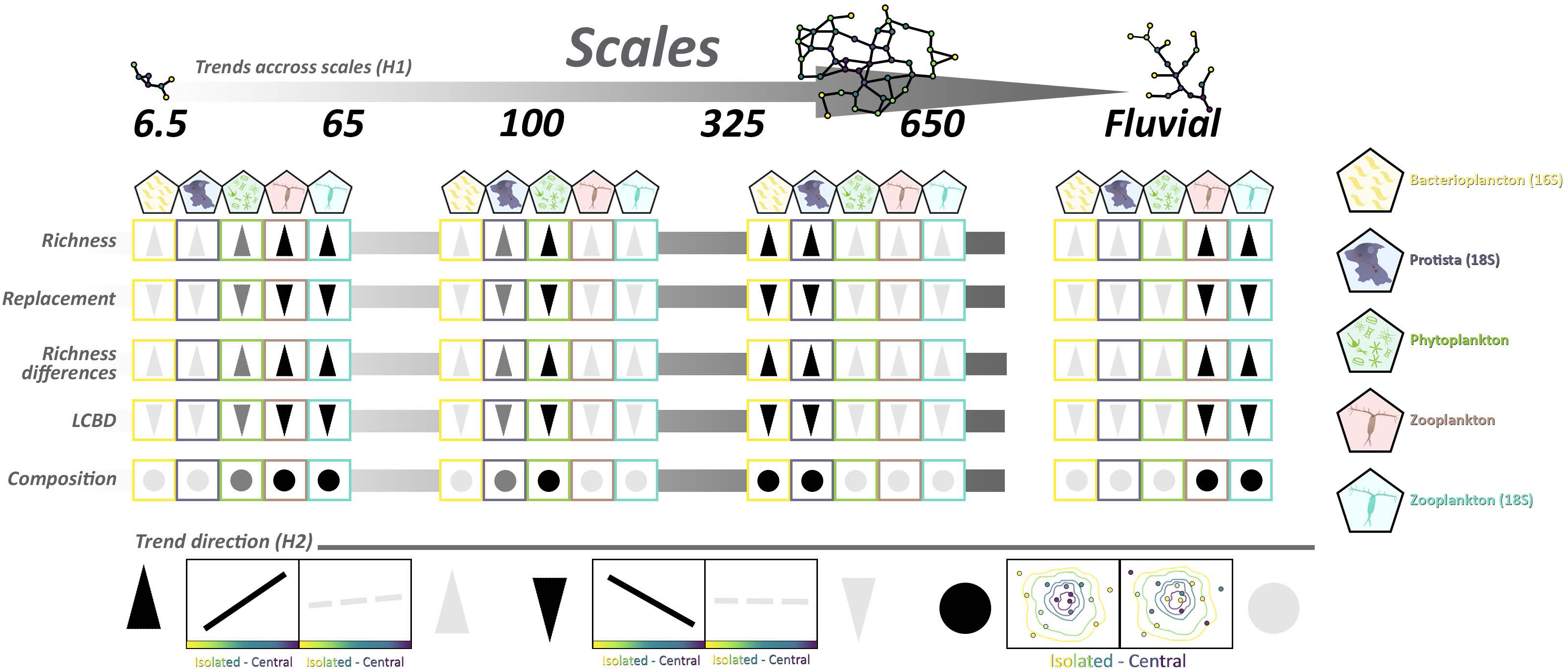
Hypothesised trends for each taxonomic group along the six scales defined in this study for the four metrics of community (richness, replacement, richness difference and LCBD) and multivariate analysis (dbRDA). Each scale corresponds to a network built out from all lakes found at 6.5, 65,100, 325, 650 km and along the fluvial network of the sampled lakes from which centrality gradients were calculated (see methods). Centrality gradient is represented in the x axis of the plots ranging from isolated (yellow) until central (purple) lakes. Dark solid lines (or symbols) represent significant trends and grey dashed lines (or symbols) non-significant trends.

In this study, we explore the connection between network centrality and how diversity patterns interact with the environment in planktonic communities. Our goal is to enhance our understanding of the role that landscape structure plays in influencing the assembly of these taxonomic groups. To explore this, we obtained data from four groups of organisms ranging in size and ecological role (i.e. bacterioplankton, protists, phytoplankton and zooplankton) from 55 alpine lakes located across Austria, Germany, and Switzerland. Using the sampled lakes as reference, we constructed six networks that cover various landscape scales, considering the neighbouring lakes (ranging from 6.5 km to 650 km) as well as the fluvial network. Based on these data, we were able to relate how diversity and composition of planktonic communities were affected by the centrality gradient calculated at each scale. Across all these elements, we hypothesise that 1) large-bodied organisms (e.g. zooplankton) will relate with centrality at small scales (6.5 or 65 km) or the fluvial network while we expect the opposite for small-bodied organisms (Figure 1-H1). Nevertheless, we also hypothesise that 2) the sign of the relationships will be shared among groups being more central lakes, the ones consisting of richer and more similar communities contrasting with isolated ones (Figure 1-H2), thus raising community mismatch with lakes environmental characteristics.

## Methods

### Sampling procedure in the field

The general sampling protocol is described in (Horváth et al. 2017). Briefly, we sampled 55 lakes in the perialpine area of Austria, Southern Germany and Switzerland in 2011 & 2012 (Switzerland: n = 20; EAWAG sampling, summer and autumn 2011; Austria and Germany, WasserCluster sampling, summer and autumn 2012; n = 27 and 8, respectively; see Figure 2). Coordinates for all 55 lakes can be found in Supplementary S1. Five environmental variables were recorded for each lake: water temperature (°C), chlorophyll *a* concentration (μg/l), electrical conductivity (µS/cm), lake area (hectares), and altitude (m). These variables were used as proxies of the lakes’ main environmental characteristics and the level of anthropogenic influence. Sampling was carried out by a boat, with smaller lakes being sampled above their deepest points, and larger lakes at depths >15m, except for Hubertussee, where water was collected from the outlet of the lake. Water samples were collected by a tube sampler from the upper mixed layer (i.e., epilimnion), following inspection of temperature profiles which were measured prior to collecting the water samples. Phytoplankton samples were obtained following the same protocol as with zooplankton (see Horvath et al. 2017) and fixed with Lugol’s iodine solution. Sampling equipment was always rinsed with lake water before collecting water samples and cleaned by dilute hypochloric acid solution in between lakes. Analytical protocols for recording and measuring standard limnological variables are outlined in (Horváth et al. 2017). For molecular analysis 300-800 ml water samples were filtered through 0.2 µm nitrocellulose membrane filters until they were clogged. The equipment for filtration was thoroughly rinsed with water and cleaned with ethanol in between lakes to avoid contamination. Molecular filters were frozen immediately after filtration and kept stored at -20°C until further processing. Unfortunately, we could not obtain the same number of samples for each one of the targeted taxonomic groups and therefore we eventually used 52 lakes for bacterioplankton based on 16S, 47 for protists based on 18S, 50 for phytoplankton, 52 for zooplankton obtained by microscopic analyses, and 49 for zooplankton based on 18S of a total of 55 lakes (Figure 2).

**Figure 2:**
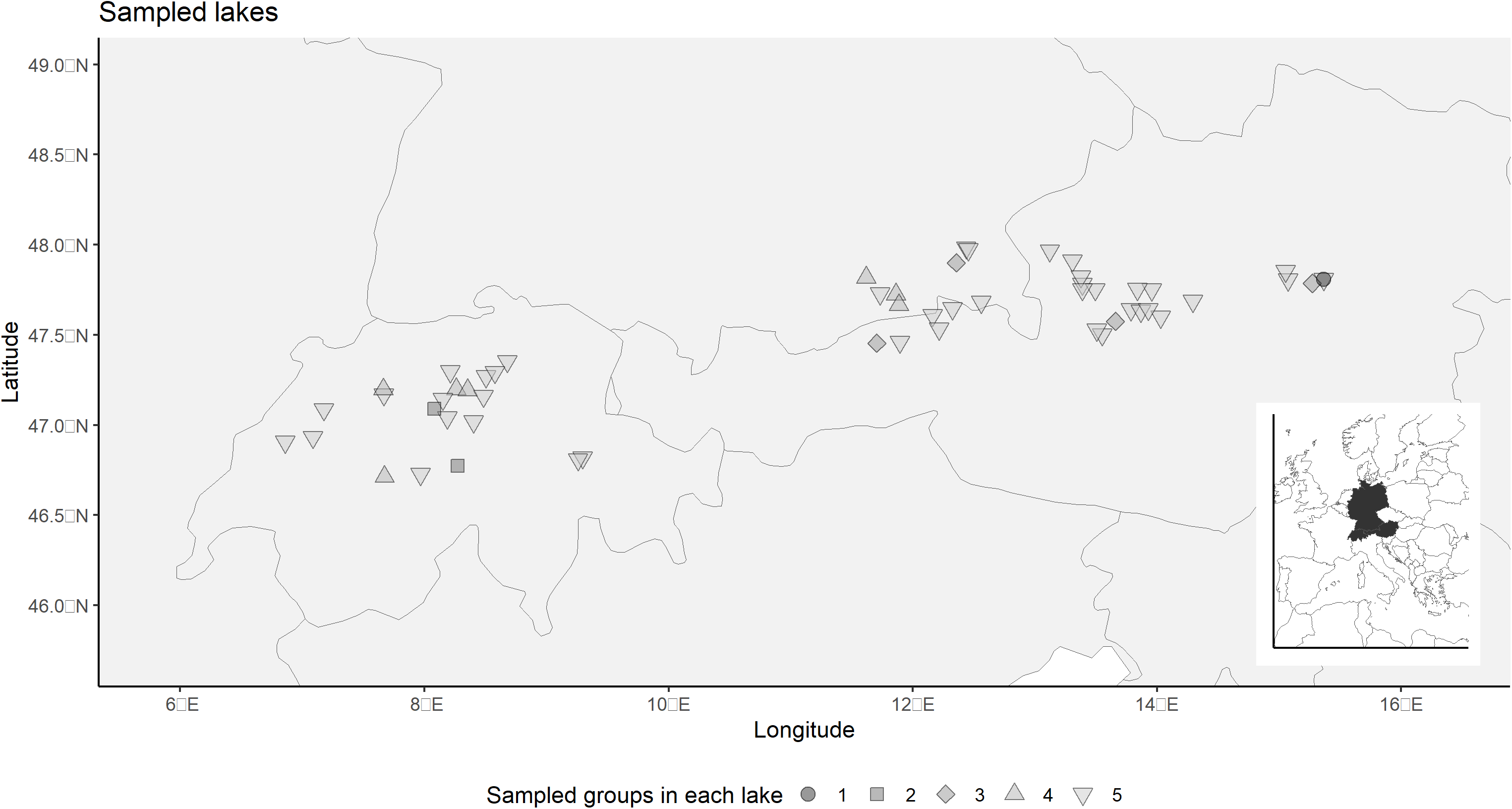
Sampled lakes locations and the number of groups that we sampled in each one of the lakes, not all groups were samples in all lakes. The smaller map highlights in black the three countries containing sampled lakes: Austria, Germany and Switzerland.

### Molecular analyses for 16S and 18S

DNA from the molecular filters was extracted using the MoBio PowerSoil DNA isolation kit and amplified with general primers targeting the 16S rRNA gene of bacteria (Caporaso et al. 2011) and 18S rRNA gene of eukaryotes (Hugerth et al. 2014). The two-step PCR amplification was carried out according to Caporaso et al. (2011) and included sample-specific barcode combinations. Illumina MiSeq sequencing libraries were prepared at the Centre of Genomic Research in Liverpool using Nextera kits (Illumina). For 16S, overlapping forward and reverse reads were combined. For 18S only the forward direction reads, ∼180 bp were used. Raw sequencing reads were quality filtered and OTU clustered (16S: 97% identity, 18S: 95% identity) using the VSEARCH pipeline (Rognes et al. 2016). OTUs were taxonomically classified using a manually curated version of the SILVA database with the CREST LCA Classifier (Lanzén et al. 2012). Through this molecular processing and classification, we obtained an OTUs and species list for bacterioplankton, protists, and zooplankton (18S zooplankton). The raw sequence data have been submitted to the European Nucleotide Archive under project PRJEB70292 and the processed datasets are available in the online supplementary material.

### Microscopic analysis of phyto- and zooplankton

A detailed outline on the microscopic analysis of zooplankton is given in Horváth et al. 2017. Overall, all crustacean zooplankton were identified to the minimum taxonomic level possible (species or species complex). Phytoplankton were quantified by inverted microscopy, following standard protocols (Utermöhl 1958, Başoğlu et al. 2017). Depending on the biomass, 20-100 ml were allowed to settle for 24-48 hours. A minimum of 500 cells were identified and counted, employing different magnifications. The raw datasets are available in the online supplementary material.

### Community structure and composition

In total, we obtained five community datasets of different organism groups: bacterioplankton, protists, phytoplankton and zooplankton. For the latter, we independently considered datasets originating from morphological identification (“zooplankton”) and molecular identification (“18S zooplankton”). In all these five datasets, we used binary data on species or OTUs (presence/absence) after the exclusion of rare species (<10 sites for bacterioplankton and 18S zooplankton, <3 for protists, phytoplankton, and zooplankton). With these datasets, we analysed planktonic diversity by calculating specific community metrics (species richness, average replacement, richness difference and LCBD) and multivariate responses (dbRDA). Species richness (Rich) corresponded to the number of taxa found in each lake. Replacement and richness differences correspond to the beta diversity partitions that we averaged for each lake to obtain a unique sample value. Replacement (Repl) refers to what extent the composition of each pair of communities corresponds to species substitutions, whereas richness difference (Rich-Diff) informs to what extent this difference can be attributed to differences in species numbers between each pair of lakes (Legendre 2014). Local contribution to beta diversity, hereafter named LCBD, was used to quantify the relevance of each lake at a regional level, providing an estimate on how unique its community is with respect to all the other lakes (Legendre and De Cáceres 2013). For average replacement, average richness difference and LCBD we used Jaccard’s dissimilarity index for presence absence data from the Podani’s families of indices (Borcard et al. 2011, Podani and Schmera 2011, Legendre and De Cáceres 2013). Finally, for the multivariate analysis we carried out a distance-based redundancy analysis (dbRDA) also using the Jaccard index. All these analyses were done with the *vegan* package (Oksanen et al. 2010). For the calculation of all dissimilarity metrics and multivariate analysis, we considered only the lakes that were belonging to the same network for the cases where more than one subgraph was built (i.e. 65 km, 6.5 km and fluvial scales; see next paragraph).

### Landscape scales and network construction

In order to create a set of different networks representing different spatial scales, we defined a total of five different landscapes by using the geographic position of all lakes surrounding the 55 sampled lakes based on a global database (Lehner and Do 2004). These five scales represent five different landscapes capturing lake distribution surrounding the sampled lakes. We used these five landscapes to construct five networks from which to calculate centrality values. First, we calculated the maximum distance between all the sampled lakes (from Neuenburgersee to Hubertussee of around 650 km). Second, we identified all the lakes within a buffer of that distance (resulting in a total of 2406 lakes in our dataset). This landscape constituted the largest scale (hereafter referred to as 650 km). We then repeated this procedure but dividing the scale by approximately half at each step to decrease the considered scale and the number of lakes included in each buffer. Thus, modifying the obtained potential network structure. This resulted in four landscapes with the following scales: 325 km including 771 lakes, 100 km including 185 lakes, 65 km including 127 lakes and 6.5 km including 58 lakes (Supplementary S2).

With the five landscapes, which included the 55 sampled lakes, we constructed five undirected networks (Figure 3). In these networks, each lake was considered a node and each connection an effective dispersal route. We connected these networks using the percolation distance, which is defined as the minimum linkage distance needed to connect all network nodes to at least one neighbour (Keitt 1997, Urban and Keitt 2001, Solé 2012). This distance was of 142 km for the 650 km scale, 71 km for the 325 km scale, 68 km for the 100 km scale, and of 6 and 6.9 km for the two subgraphs of the 65 and 6.5 km scales. We used the percolation distance because it allowed us to connect all nodes within one unique network while conserving the regional structure caused by the distribution of nodes in the landscape. Also, it allows consideration of all nodes when calculating centrality metrics. Thus, percolation networks represent the intermediate transition between a disconnected and a global network, where all nodes are potentially reachable but still preserve their regional structure (Borthagaray et al. 2015a). For each network we calculated closeness centrality, which represents the centrality gradient (Figure 3). Closeness centrality is the reciprocal of the average distance from the focal node to all the other network nodes, where higher values represent more central habitats in the network (Borthagaray et al. 2015a, Dale 2017). The two smaller scales (65 km and 6.5 km) consisted of two separated subgraphs and their closeness centrality values were calculated separately for each subgraph and later normalised dividing the values by each graph maximum. By including these scales in our analysis, we assumed that dispersal at these two scales was not occurring between subgraphs and maintained each subgraph centrality gradient avoiding the effect of having different numbers of nodes. Overall, with the five landscapes, we generated five networks and hence five gradients of centrality using the percolation distances (Figure 3).

**Figure 3:**
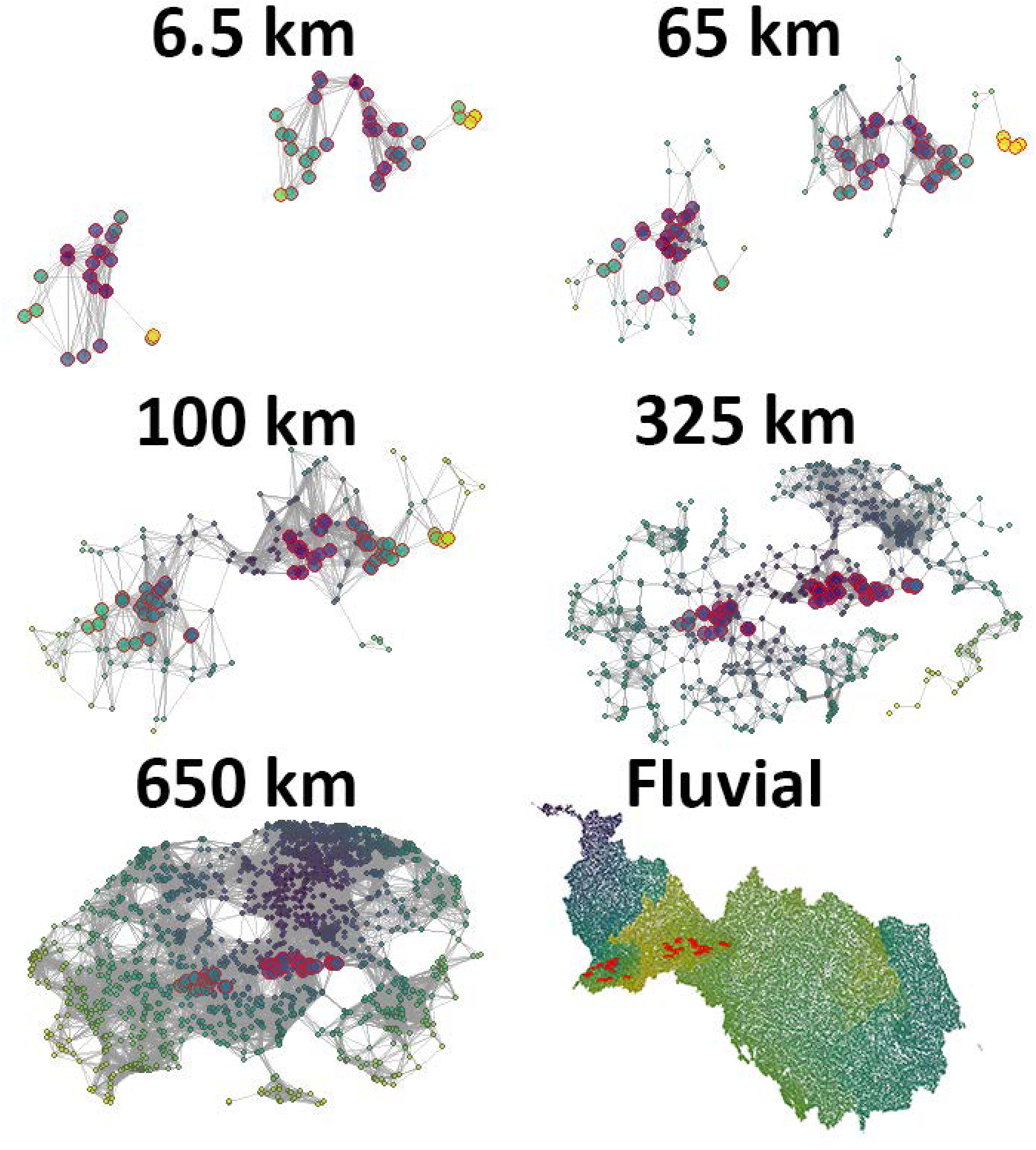
The five networks generated using global lakes database. From top-left to bottom-right there is 6.5 km, 65 km, 100 km, 352 km, 650 km, and fluvial respectively. Red contoured circles indicate the 55 sampled lakes. Circle fill colours indicate closeness centrality being the more purple/blue circles the ones more centrally located and green/yellow circles the ones in a more isolated position.

The fluvial directed network was created using the two river networks where the sampled lakes belong (the Danube and the Rhine river catchments: Figure 3F). We obtained the shapefiles from these two fluvial networks from HydroRIVERS and HydroBASINS databases (Lehner et al. 2008, Lehner and Grill 2013). Then, we transformed the shapefile into a graph where each segment intersection was considered as a node using the package *shp2graph* (Lu et al. 2018). Once the network was defined, we located the position of the sampled lakes in it. Finally, we calculated the centrality isolation gradient considering a directed network from up- to downstream (i.e., out-closeness centrality). We considered a directed network to capture the natural dendritic structure of the rivers to also account for drift dispersal through the fluvial network. The two fluvial catchments consisted of two different subgraphs as the 65 and 6.5 km scales, thus, we followed the same approach and calculated centrality values (i.e. out-closeness) separately for each subgraph and later normalised dividing the values by each subgraph maximum. As out-closenness presented a marked skewness due to the dendritic network of rivers we additionally log-transformed their values to correct this. Overall, out-closeness presented the same gradient as the values of undirected networks with lower values of out-closeness representing more isolated parts of the stream basins (i.e. headstream nodes; Figure 3) and *vice versa*. For the five undirected networks and the fluvial network calculations and representation, we used the packages *igraph* (Csárdi and Nepusz 2006) and *sna* (Butts 2023).

### Centrality-isolation against community facets

We followed two different procedures for community metrics and the multivariate analysis to relate them against centrality but in both cases we excluded environmentally explained variation in order to focus on the pure effect that the centrality gradient had on diversity and composition. For each one of the community metrics (Rich, LCBD, Repl, and Rich-Diff) we first selected the most relevant environmental variables using a random forest (randomForestSRC package, (Ishwaran and Kogalur 2023). Second, we conducted a linear model with each community metric as response variable and the selected environmental variables as explanatory variables. Third, we extracted the residuals of that linear relation retaining the unexplained part of this relationship. Finally, we used the residuals as a response variable against the centrality values of each one of the six networks using a generalized additive model (GAM; mgcv package, Wood 2017). We considered the residuals of each diversity metric as the response variable and the centrality-isolation gradient as the smooth term. Our focus on using residuals aimed to capture the discrepancies between environmental variables and the diversity patterns observed in planktonic communities, exploring their relationship with centrality measures (i.e. closeness and inverse out-closeness). In essence, any significant trends identified would indicate the pure impact of the centrality gradient on community assembly, independent of underlying gradients in environmental drivers and eliminating any potential autocorrelation between the environment and centrality.

For community composition (dbRDA; (Oksanen et al. 2010), we first subtracted the unexplained coordinates of the first two axes of the dbRDA. Second, we fitted a GAM using the *ordisurf* function of the vegan package (Oksanen et al. 2010) into the unexplained multivariate space. Here, centrality (closeness and out-closeness) was fitted as a smooth surface using penalised splines in GAM (Wood 2003) to later predict its fitted values in a regular grid. In summary, this function plots the fitted contours with a convex hull of data points over the existing unexplained multivariate space and calculates its significance (Oksanen et al. 2010). The two analyses considering community metrics and multivariate analysis were carried for each one of the five considered taxonomic groups (Bacterioplankton, Protists, Phytoplankton, Zooplankton and S18 Zooplankton) and considering the closeness values of each network (650 km, 325 km, 100 km, 65 km, 6.5 km and fluvial). So, overall, we conducted a total of 120 models for community structure and 30 dbRDA for community composition. From these we selected only the significant smooth terms (p<0.05), which indicated that there was some significant trend between community structure or composition and the centrality.

## Results

### Community structure and composition

While there were major differences in the five taxonomic groups in their richness values, inherent to the two main identification pipelines (microscopic versus molecular), average LCBD values were quite similar between the groups (Table 1). Within LCBD, protists, zooplankton and S18 zooplankton presented a greater standard deviation, indicating a greater variation in uniqueness of their communities in comparison to Phytoplankton and Bacterioplankton. Average replacement and richness also showed differences between groups that overall indicated the strong variation at the regional level of each group (Table 1). Average replacement was markedly high for bacterioplankton and phytoplankton but bacterioplankton presented the lowest values of richness difference. Contrastingly, other groups had much more similar beta diversity partition values such as protists and S18 zooplankton. Tables with the specific values for each sampled lake as well as the plots representing these metrics can be found in Supplementary S3 and S4.

**Table 1.**
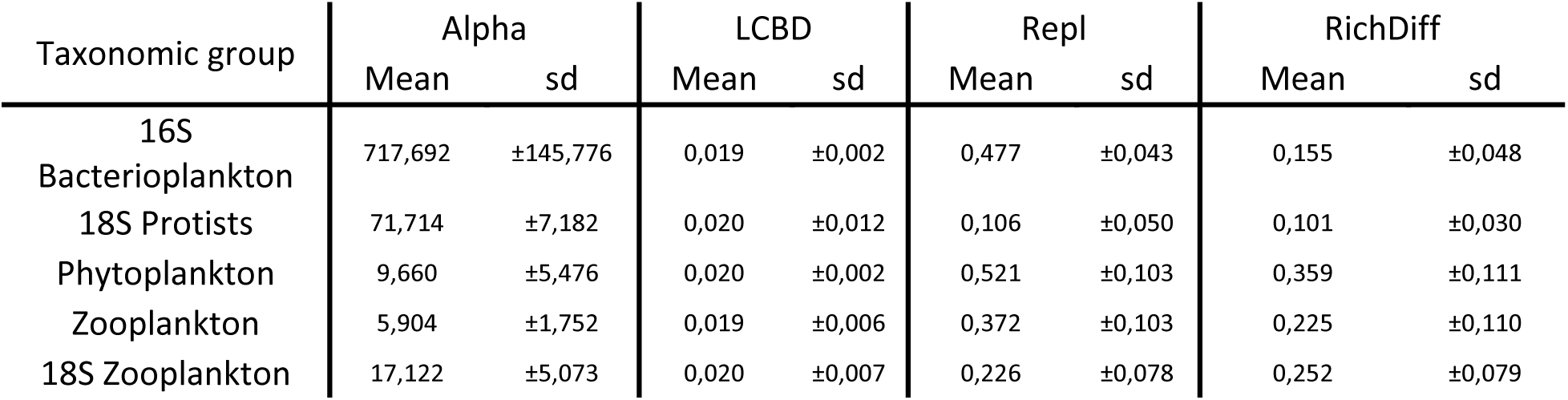

### Landscape scales and network building

The six networks that we built presented some differences between the structure that they were capturing (Figure 3). For example, 650 km and 325 km networks represented markedly different scenarios, for 650 km the network was much denser and many links were present while for 325 km the structure was much less connected and related to the distribution of nodes through the landscape (Figure 3). On the other hand, 65 km and 6.5 km networks were more similar in terms of connections between lakes even though the first had two times the number of lakes of the other (Figure 3). Finally, the fluvial network was unique among the other networks as it was directed (i.e. each node was connected to its downstream neighbour following a head-to downstream sense). The parameters describing the six built networks can be found in Supplementary S5. For all the networks, the sampled lakes were located along the entire centrality-isolation gradient indicating that sampled lakes were representing most of the variability in closeness centrality of each network (Supplementary S6).

### Centrality-isolation against community facets

#### Community metrics

The GAMs conducted for each one of the five organism groups (bacterioplankton, protists, phytoplankton, zooplankton, and 18S zooplankton) using each one of the residual values from the four community metrics (Rich, Repl, RichDiff, and LCBD) as response variables and the centrality gradient from the six networks (6.5, 65, 100, 325, 650 km and fluvial) as smooth term showed differences between taxonomic groups and across scales (Figure 4). Mostly bacterioplankton, zooplankton and 18S zooplankton showed significant trend with centrality isolation gradient at some scales (650km, 320km, 100km, 65km and Fluvial) while protists only appeared related with one scale (Fluvial) and phytoplankton did not show any significant trend. From these, 18S zooplankton was the group presenting more significant trends of all the studied ones being the only group where Richness difference and LCBD values showed some significant trends (Figure 4). See all model results in Supplementary material S7.

**Figure 4:**
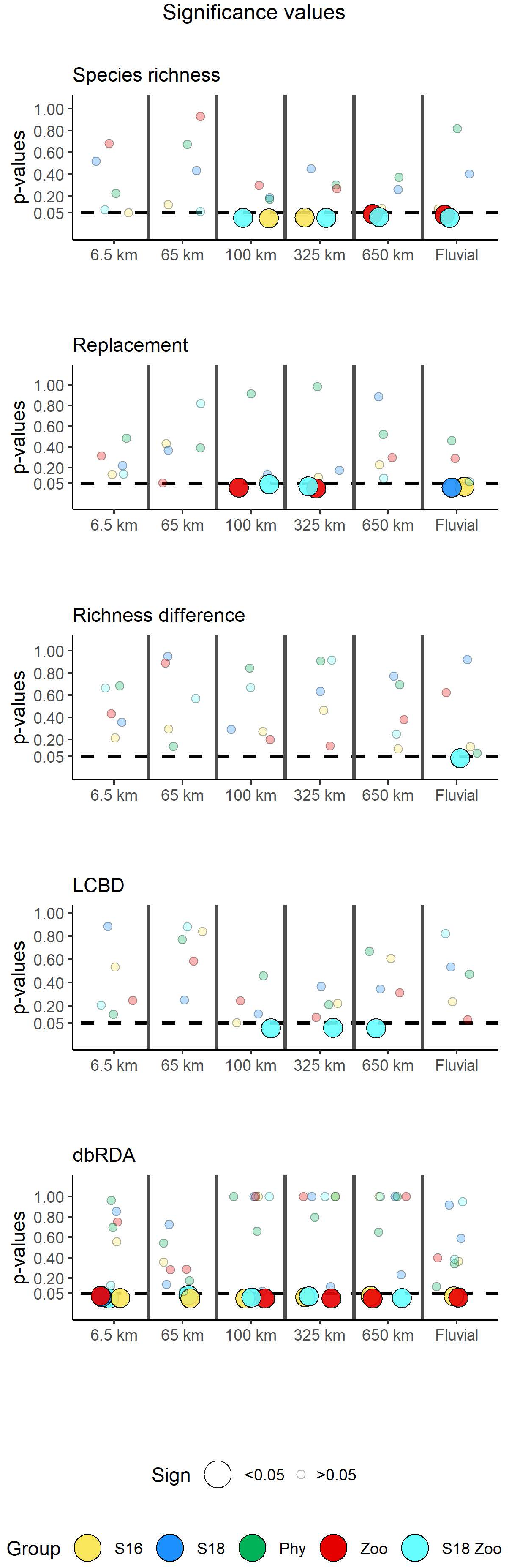
P-values for all generalised additive models considering closeness centrality as smooth term for community metrics (Rich, LCBD, Repl, RichDiff) and for the multivariate analysis ordination surface (dbRDA). Results are represented for all the networks (6.5, 65, 100, 325, 650 km and Fluvial) and considering the five organism groups (Bacterioplankon in yellow, Protists in blue, Phytoplankton in green, Zooplankton in red, and 18S zooplankton in cyan). Larger coloured circles indicate the models with a significant smooth term (i.e. below 0.05). Note that point location along the X axis is randomly assigned within each network.

From the significant trends that we observed, species richness was significant for bacterioplankton, zooplankton and 18S zooplankton (Figure 5). Richness was significant for the networks built at 100, 325, 650 km, the Fluvial and for most of the groups it presented higher residual values at more central sites (Figure 5A:H), thus pointing to a greater mismatch between environment and species richness in these locations. Only for bacterioplankton it presented a more U-shaped trend for alpha diversity with lower residuals in sites with intermediate centrality values. In contrast, for the Fluvial network these patterns were reversed with lower residuals in more downstream sites (Figure 5D & H). Replacement presented a negative trend being more central communities, the ones having lower residual values for both zooplankton and 18S zooplankton at 100 and 325 km networks (Figure 5I,J,L & M) and thus showing a better match with environmental constraints for replacement. On the other hand, bacterioplankton and protists replacement was negatively related with centrality but from the fluvial network (Figure 5K & N), being downstream sites better explained by environment. Richness difference was only related with 18S zooplankton for the fluvial network (Figure 5O), having a positive trend with higher residuals in more downstream lakes. Finally, LCBD had significant trends only for the 18S zooplankton at 100, 325, 650 km, being more central sites, the ones having lower residuals and thus having a stronger link with environment (Figure 5P:R). Overall, all the significant models explained 10-20% of deviance and had consistent patterns across scales and even across groups for the significant community structure metrics (Figure 5). In general, there was a positive association between species richness residuals and centrality, indicating a disconnection from environmental factors in central positions. Conversely, the residuals for average replacement and LCBD exhibited a negative relationship with centrality, suggesting that in central locations, replacement and LCBD were more effectively explained by environment. Most observed trends followed a monotonous pattern, apart from bacterioplankton, which displayed distinctive U-shaped trends (Figure 5).

**Figure 5:**
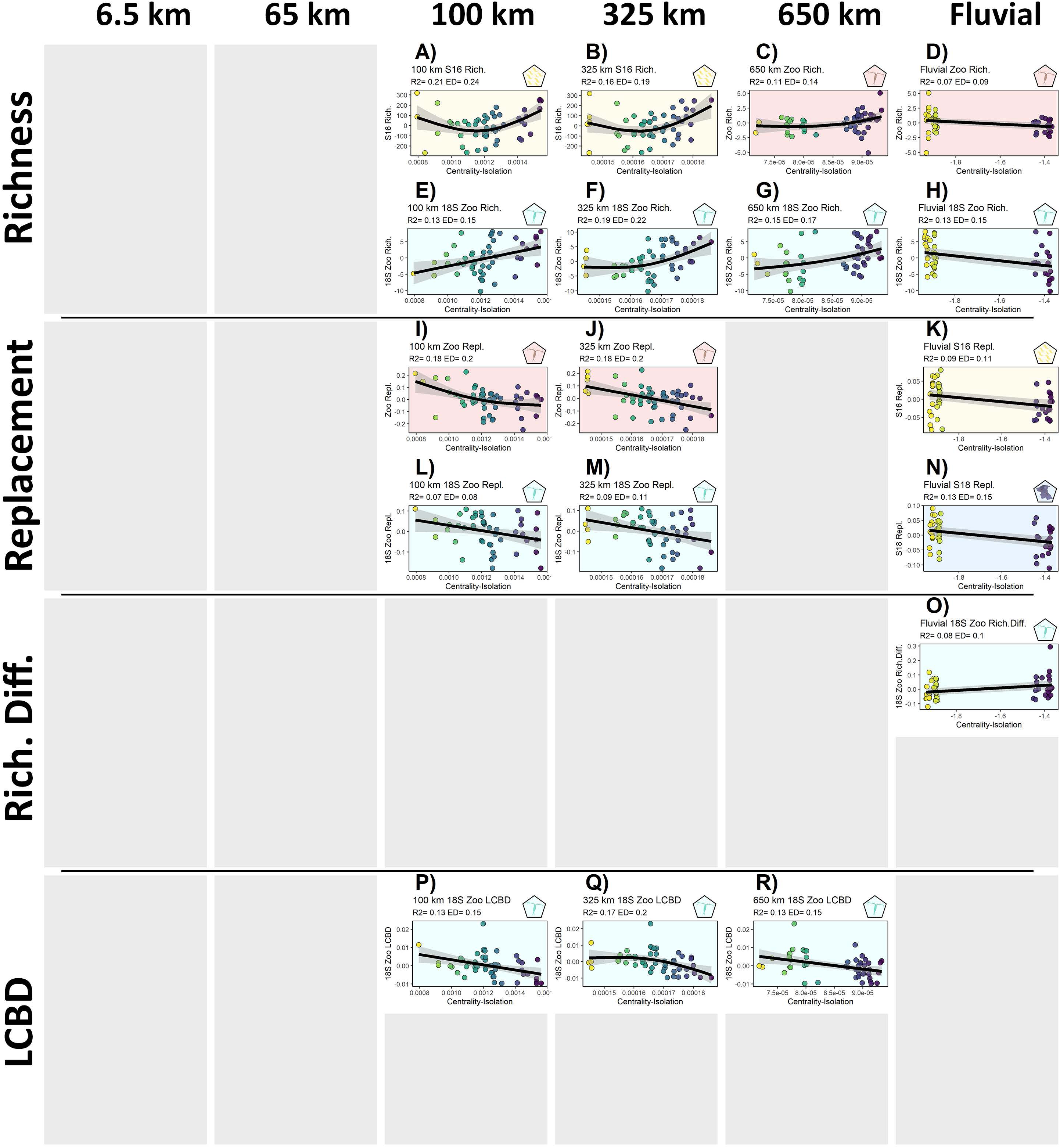
Significant trends for each community metric against centrality based on generalised additive models (GAM). Plot background and top-right symbol indicate the corresponding organismal group: bacterioplankton (yellow), zooplankton (red), and 18S zooplankton (cyan). Circle colours represent centrality gradient (same as X axis) to facilitate visualisation, purple colours correspond to more central lakes and yellow colours more isolated lakes. Out of the 120 models, only the 17 significant models (p<0.05) are presented here. See all model results in Supplementary material S7.

#### Multivariate analysis

For the multivariate analysis we observed similar results than with community metrics, the groups presenting a significant smooth surface against centrality were basically bacterioplankton, zooplankton and 18S zooplankton (Figure 6) with protists being only significant in one case (Figure 6G). Bacterioplankton had a significant relationship across all networks (Figure 6A:F). For networks encompassing larger scales (e.g. 650km), more central sites tended to be more separated (Figure 6E & F). On the other hand, as scales decreased (e.g. 65 and 6.5km) this pattern was the inverse and more isolated lakes were the ones being in more external positions of the multivariate space (Figure 6A & B). For these smaller scales only one subgraph showed a significant relationship for composition (the lakes within Austria and Germany) for bacterioplankton, protists and zooplankton at 6.5 km scales (Figure 6A, G & H) and bacterioplankton at 65 km scales (Figure 6B). Contrastingly, 18S zooplankton appeared significantly related with 6.5 and 65km scales (Figure H & M) but only from the other group of lakes (lakes within Switzerland). For these scales, protists, zooplankton and 18S zooplankton showed more similar communities among isolated lakes. Finally, zooplankton and 18S zooplankton presented the same significance trends for 100, 325, 650 km (Figure 6I:K & O:Q). In these cases, the relationship with the centrality gradient was similar being more central sites, the ones tending to have similar composition through all the significant scales. See all dbRDA model results in Supplementary material S8.

**Figure 6:**
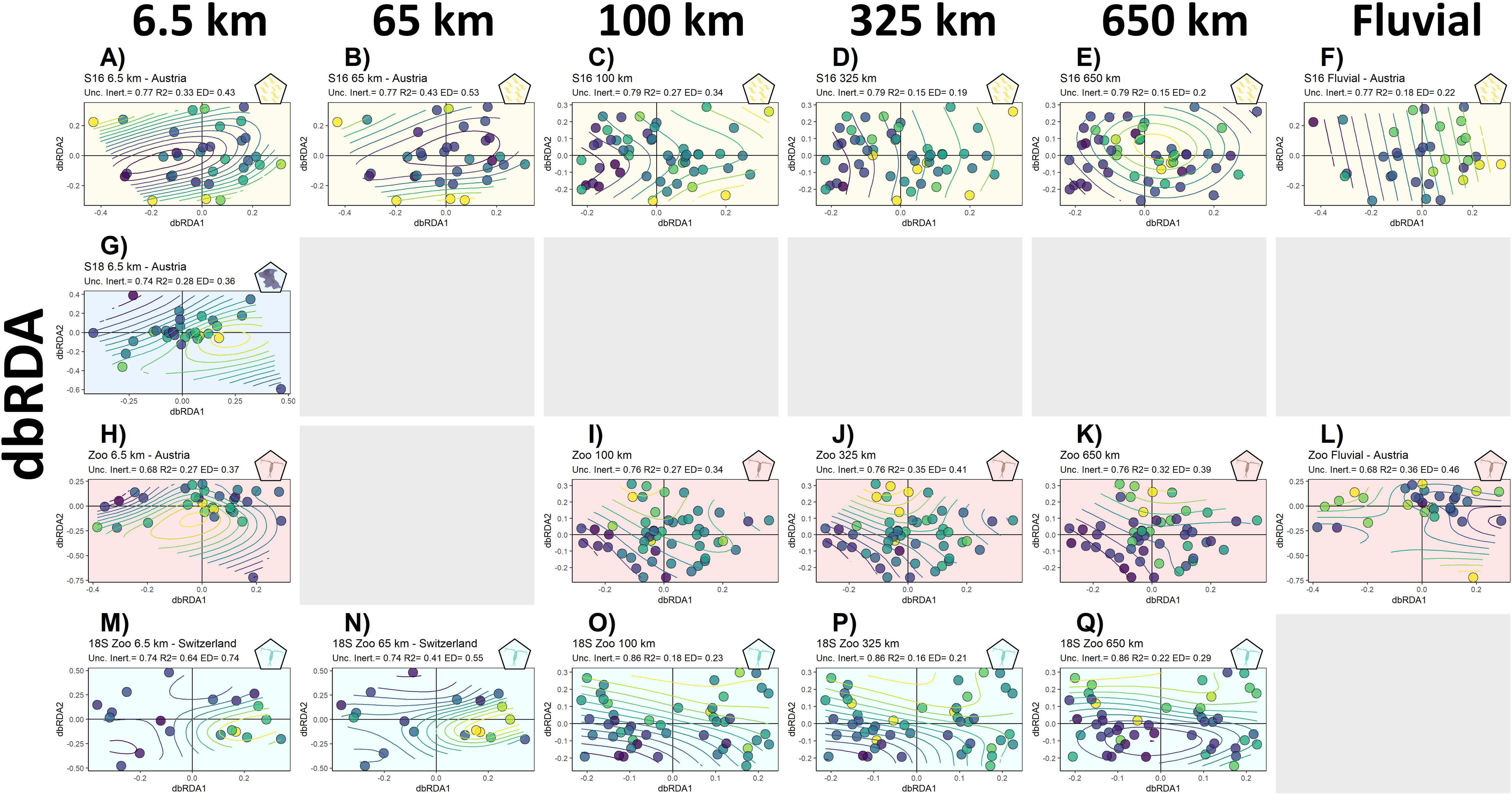
Significant multivariate analysis based on the generalised additive models (dbRDA with smooth surfaces). Plot background and top-right symbol indicate the corresponding taxonomic group: bacterioplankton (yellow), zooplankton (red), and 18S zooplankton (cyan). Contour lines and circle colours represent centrality gradient, purple colours correspond to more central lakes and yellow colours more isolated lakes. See all model results in Supplementary S8.

## Discussion

The impact of spatial structure on diversity patterns has been broadly discussed in the last years of ecological research (Turner and Gardner 2015, Leibold and Chase 2018, Erős and Lowe 2019, O’Connor et al. 2020). It connects directly with main assembly patterns as spatial structure defines the canvas in which metacommunities assemble and therefore, it drives diversity of species that are able to move throughout it (Vellend 2016, Thompson et al. 2020). In the current work, we highlighted the strong role that landscape structure plays as a driver of alpine lakes planktonic communities as well as the scales at which it takes place (i.e. 650, 325, 100 km and Fluvial). In these systems, landscape structure has generally been attributed to the fluvial connection between lakes (Almeida-Gomes et al. 2020). However, here, we saw that not only fluvial structure affects diversity patterns and that bacterioplankton, protists and zooplankton diversity appear also shaped at undirected network structures (i.e. 650, 325, 100, 65 and 6.5 km of distance). Furthermore, Phytoplankton did not show any relationship with centrality of these landscape structures, neither fluvial nor undirected. Such findings emphasise the complexity in quantifying spatial structures that appear relevant for diversity patterns and the strong variability that can exist between taxonomic groups (Boix et al. 2017, Tonkin et al. 2018a, Newman 2019), and they contribute to gather new insights in how, and more specifically at which scales, planktonic and microscopic organisms are being affected by centrality and thus the scales at which metacommunity processes might be taking place for them (Leibold and Chase 2018, Custer et al. 2022).

On the one hand, our hypotheses were partially refuted because we observed that organisms did not respond to different scales in relation to size (hypothesis 1, i.e. Bacterioplankton and Zooplankton appeared related through similar scales), and some groups community structure and composition was not correlated with centrality (i.e. Phytoplankton). On the other hand, when a pattern was detected across several groups the sign of it was consistent for all of them, which implies that the captured process was having the same effect across groups. For example, alpha diversity responded in the same way against centrality for both bacterioplankton and zooplankton or replacement for bacterioplankton, protists & zooplankton (hypothesis 2). An increase in centrality was concomitantly increasing the mismatch between environment and species richness. This loss of connections might be related to the higher species richness of more central sites (Economo and Keitt 2010, Altermatt 2013, Borthagaray et al. 2020). Interestingly, composition responded the other way around (for 325 and 100 km landscapes), where more central lakes and downstream lakes replacement had a better fit with environment. Although not shared among groups, zooplankton beta diversity was related with several scales and for both, Zooplankton and 18S Zooplankton. In all these cases, more central communities were better explained in central positions both for composition and uniqueness in comparison with more isolated lakes. Such patterns respond to a stronger impact of dispersal in more central communities (e.g. mass effects archetype; (Leibold et al. 2004), which would favour the foster community homogenization and the overall match of its replacement and LCBD values with respect to the regional pool (Altermatt et al. 2011, Ai et al. 2013, Carrara et al. 2014, Tonkin et al. 2016a, Jamoneau et al. 2018). These patterns were found at similar landscape scales for both Bacterioplankton and Zooplankton, which meant that the network structure obtained between 650 and 100 km was actually capturing a landscape structure that was affecting these two groups beta diversity patterns (Economo and Keitt 2008, Chisholm et al. 2011, Dale 2017). Thus, these scales could represent the most probable level at which dispersal processes would be impacting these groups defining the effective landscape at which the exchange of individuals drives diversity patterns and metacommunity assembly (Vellend 2016, Leibold and Chase 2018, Thompson et al. 2020).

The observed responses highlight not only that smaller-bodied organism groups are affected by the centrality gradient (i.e. Bacterioplankton), but also that the assembly of these groups is affected across different regional scales (Custer et al. 2022, Barbour et al. 2023, Tytgat et al. 2023). Therefore, even for smaller microscopic taxa, broad-regional patterns might explain their regional diversity distribution (Tytgat et al. 2023), something that highlights the relevance for accounting for landscape and centrality metrics even for these groups (Whitfield 2005, Fontaneto et al. 2008, Vannette and Fukami 2017). Along the same lines, Zooplankton appeared related with the same scales as Bacterioplankton, which could point towards both groups being dispersed by the same vectors or by vectors that act at the same scales of hundreds of kilometres (Incagnone et al. 2015, Pinceel et al. 2016, Coughlan et al. 2017). It is worth noting that bacterioplankton was sampled via filtration in a size-inclusive manner, meaning that bacteria associated with larger particles such as Zooplankton bodies are also included in our analysis. Contrastingly, phytoplankton showed the total opposite pattern, not being affected by any of the tested centrality isolation gradients. Protists behaved similarly only showing relationships at the fluvial scale and at the smallest network scales. Based on our results, we should expect that these two groups would have been related to same scales than bacterioplankton and zooplankton, as they could also be transported by the same vectors (Naselli-Flores and Padisák 2016, Padisák et al. 2016). As a consequence, we assume that these organisms were potentially transported through the same corridors but their establishment failed either due to different environmental conditions (Abonyi et al. 2018, Chaparro et al. 2018), or due to stronger priority effects (Fukami 2015, Leibold and Chase 2018). Furthermore, these two groups, specially Phytoplankton, are primary producers which strongly rely on environmental conditions (e.g. available nutrients, stratification; (Stomp et al. 2011, Li et al. 2019, Borics et al. 2021), which, together with the fact that we focused on the pure spatial effect (i.e. centrality gradient across scales) might have hidden the spatial signal for these groups if both environment and centrality gradients where equally relevant or correlated (Chaparro et al. 2018, Loewen et al. 2020, Rocha et al. 2020). Nonetheless, the current findings point towards the relevance of spatial distribution for the diversity of two key planktonic groups such as bacterioplankton and zooplankton not uniquely related to the fluvial network. Such results stress the strength of spatial processes for these groups but also the scales at which this occurs (i.e. hundreds of kilometres) that should be considered if regional diversity patterns are to be preserved (Edge et al. 2017, Horváth et al. 2019, Brauer and Beheregaray 2020).

One limitation of our work is that our environmental variables are a limited simplification of the actual local environment. Our simplification of the environment and our focus on how well landscape structure (i.e. centrality) was explaining the mismatch between it and diversity patterns was marked by the few environmental variables available. Although we accounted for several environmental variables to tease apart their contribution to diversity, the number of metrics was short (i.e. water temperature, chlorophyll-a concentration, electrical conductivity, lake area, and altitude) and therefore some key environmental variables may not have been considered (e.g. nutrients, fish community). However, the five considered metrics also represent main proxies of lake state and human impact for alpine lakes (Jacquemin et al. 2019, Li et al. 2021). In a similar line, landscapes were built based on existing databases with a determined level of accuracy (Lehner and Do 2004). Although satellite identification of water bodies is improving (Pekel et al. 2016), there is still a lack of accuracy to detect small water bodies, which in turn can play an important role in determining network scale fluxes (Downing 2010, Borthagaray et al. 2023a). Furthermore, we connected our landscapes based on the percolation distance which represents a compromise between local and regional network structure. However, networks could have been built based on different distances representing different perceptions of the landscape (Borthagaray et al. 2015a, Dale 2017, Cunillera-Montcusí et al. 2021). These networks would have corresponded to different ways to perceive each one of the different landscapes and thus, represent different centrality gradients (Borthagaray et al. 2014, Newman 2019, Savary et al. 2023). In the current work, we focused on varying the scale at which the landscape is constructed to focus on the relevance of habitat distribution at different scales in determining diversity patterns (Gilarranz 2020, Faggioni et al. 2021, Borthagaray et al. 2023b). Nevertheless, to additionally explore how different threshold distances might also impact diversity patterns constitutes an avenue to follow in the coming years to further disentangle how planktonic communities are perceiving and interacting with the landscape (Almeida-Gomes et al. 2020, Custer et al. 2022). We employed both morphological and molecular techniques to describe community structure and composition of the different organism groups, aligned with current and emerging best practice approaches to target aquatic diversity (Hillebrand et al. 1999, Deiner et al. 2015). The detection of similar patterns for the same organism group using independent methodology (e.g. zooplankton and 18S zooplankton), strengthens our observations.

In the current work we highlighted the complexity of defining a landscape that will drive diversity patterns across ecosystems at different scales. This complexity will become more evident with the greater availability of satellite information and computationally powerful tools that will allow us to process further layers of the landscape (Newman 2019, O’Connor et al. 2020, Savary et al. 2023). Therefore, the current work is an important step toward accounting for landscape structural features that shape planktonic groups in bigger lentic systems (Almeida-Gomes et al. 2020). These features are, together with abiotic and biotic interactions, key when assessing the diversity of these communities in face of global change as they have an impact on the species that are available to assemble communities (Leibold and Chase 2018, Thompson et al. 2020). The landscape structure of Alpine lakes is also particularly threatened as alterations at the regional level on the presence of these lakes (Zhang et al. 2017), their connectivity (Monaghan et al. 2005, Robinson and Kawecka 2005) and environmental characteristics (Moser et al. 2019) is subject to global change. Thus, potentially impacting future community diversity patterns (Fuller et al. 2015, Thompson et al. 2017, Pardini et al. 2018). As a consequence, to improve the current knowledge on the relevance of landscape structure on these groups appears key to further understanding the consequences of global change on them as they constitute the basal elements of lake trophic structure. Finally, the consideration of network scale patterns for alpine lakes, moving further from lotic networks, constitutes an advance to better disentangle how these systems are impacted by landscape structure (Almeida-Gomes et al. 2020) and thus, how their metacommunities are assembled and how the layers driving diversity are acting in these threatened ecosystems (Leibold and Chase 2018, Moser et al. 2019).

## Acknowledgments

D.C.M. was supported by the H2020 EU-funded project AQUACOSM-plus (no. 871081), the H2020 EU-funded project AQUACOSM (no. 731065), by the European Union-NextGenerationEU, Ministry of Universities and Recovery, Transformation and Resilience Plan, through a call from Universitat de Girona and by the European Union’s Horizon 2020 research and innovation programme under the Marie Skłodowska-Curie grant agreement No 101062388. MMB was supported through the Austrian Science Fund (FWF) project P 26119-B25 and the German Research Foundation (DFG) project LakeMix (BE 6194/ 1-1).

---

**Supplementary S1:**
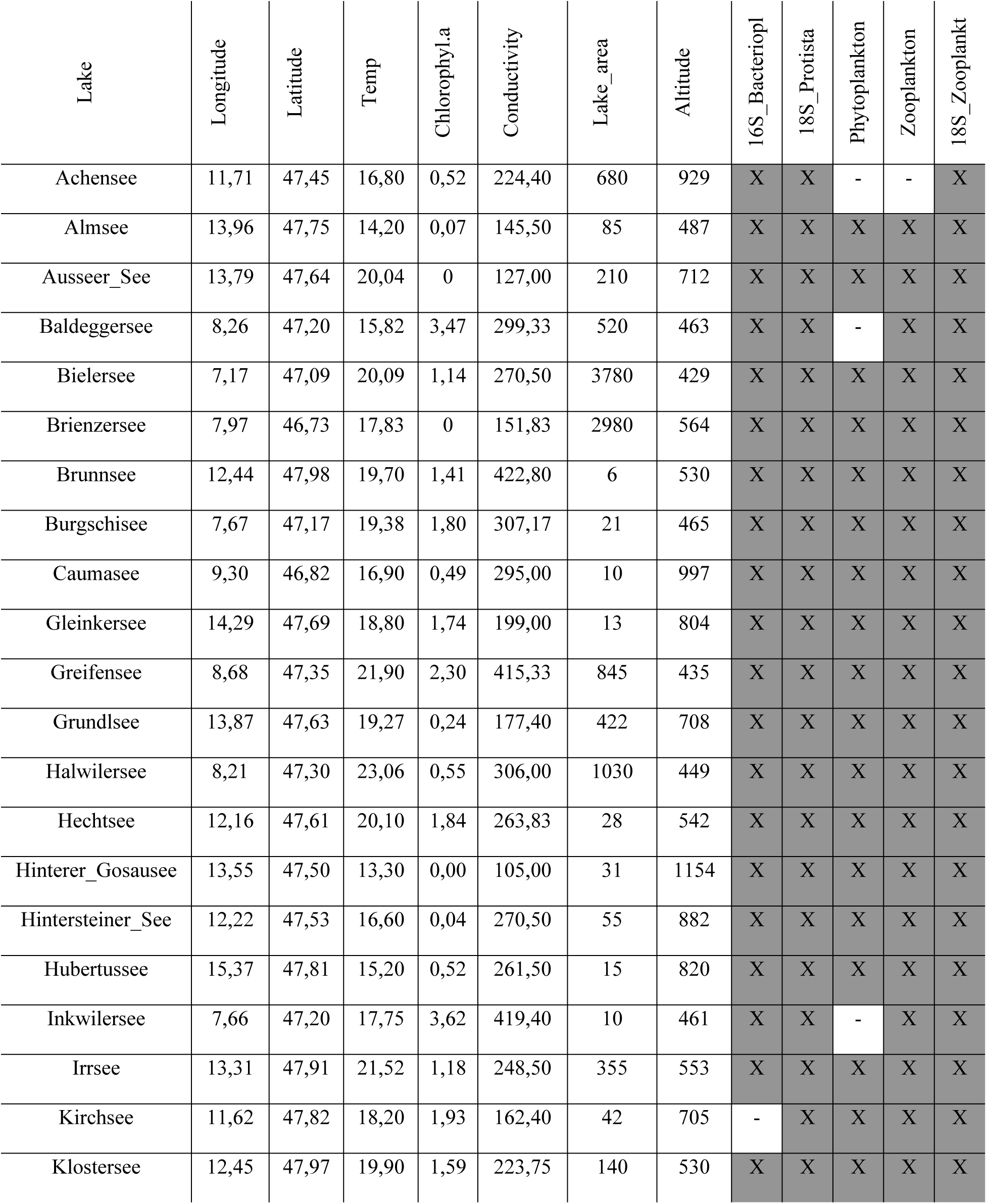
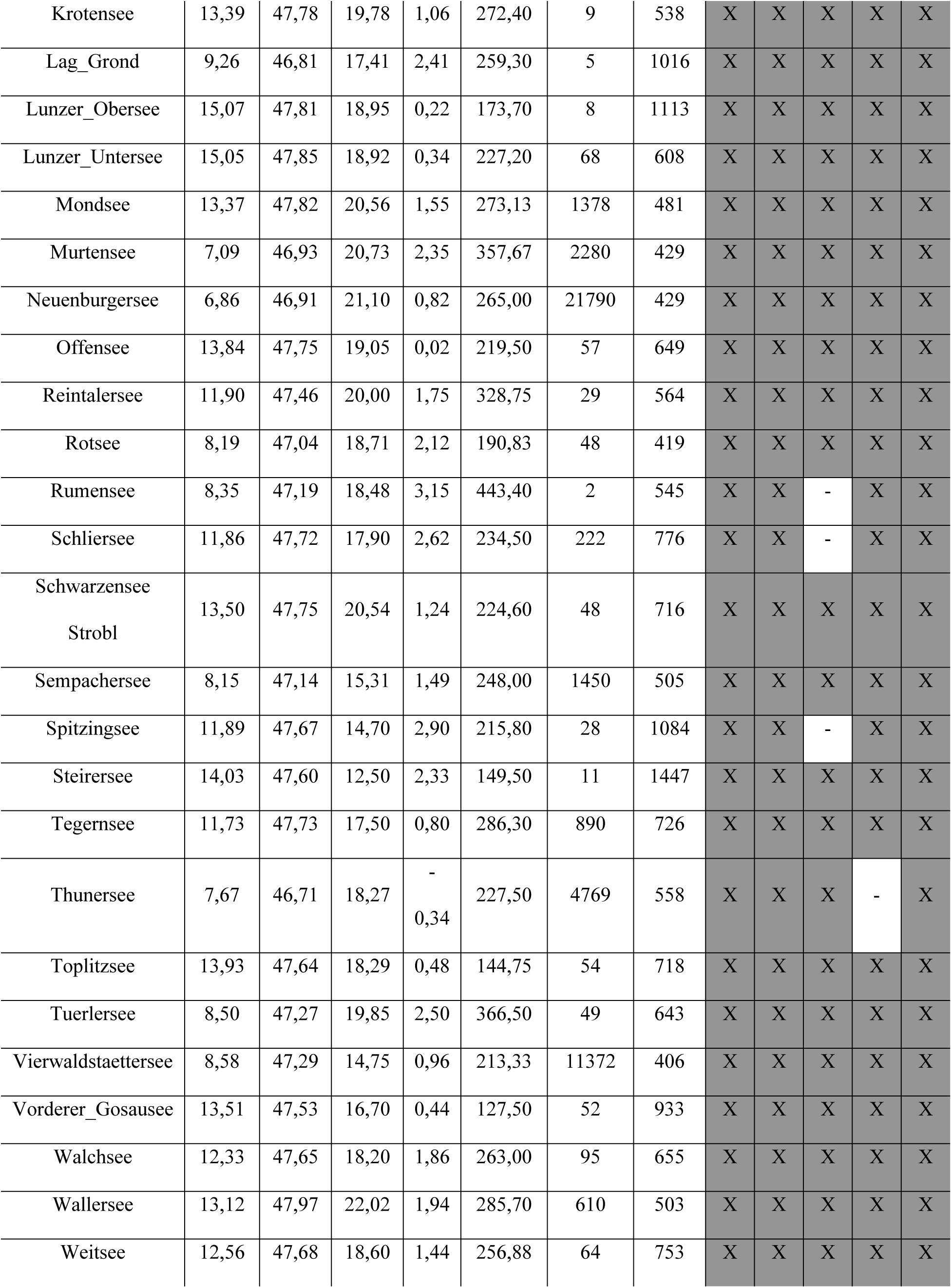
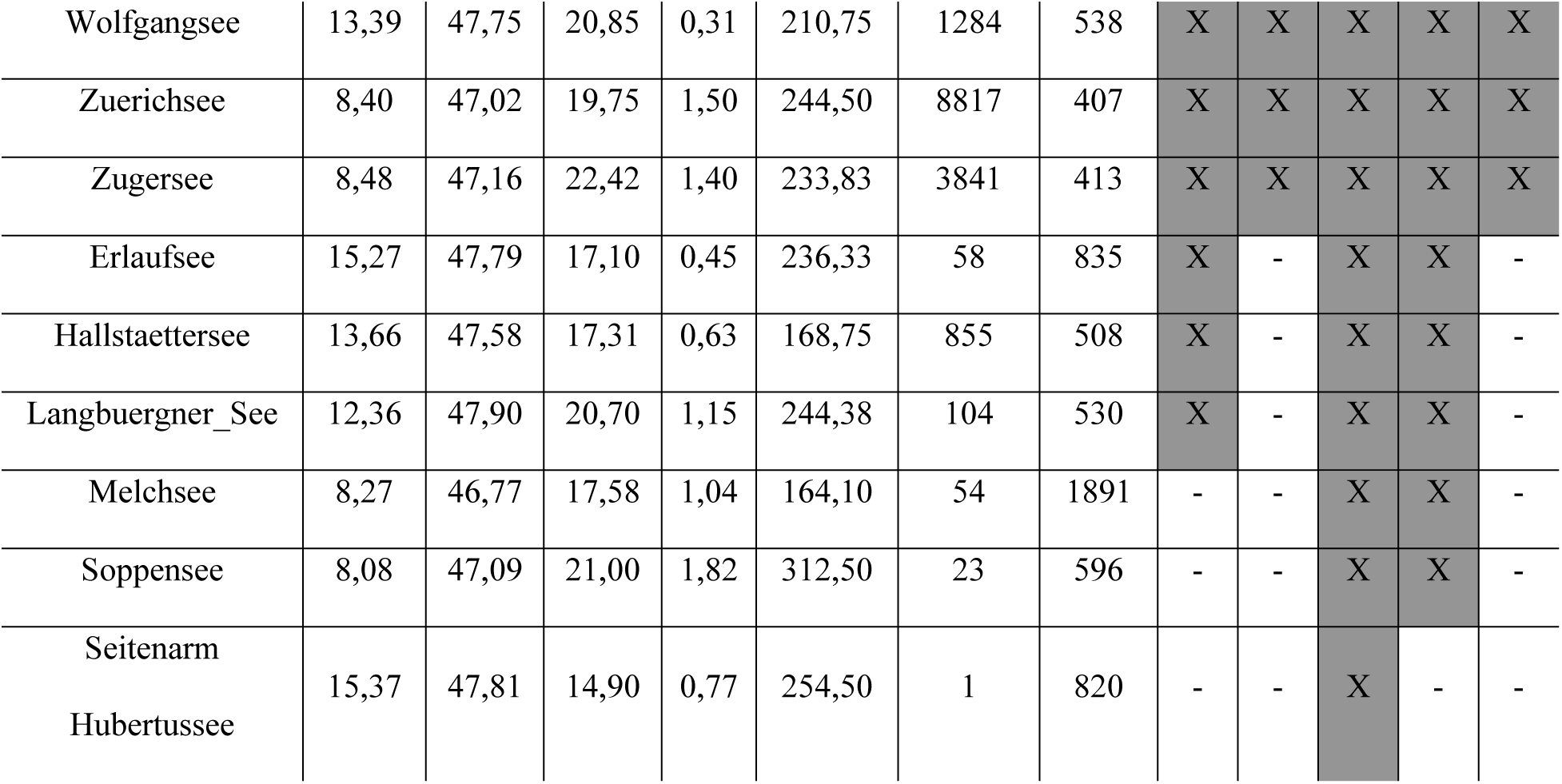
Lakes coordinates and recorded environmental variables. Not all sampled lakes were sampled for the same organism groups. Groups sampled in a respective lake are marked with an “X” and a map can be seen in Figure 2 of the main manuscript.

**Supplementary S1.2:**
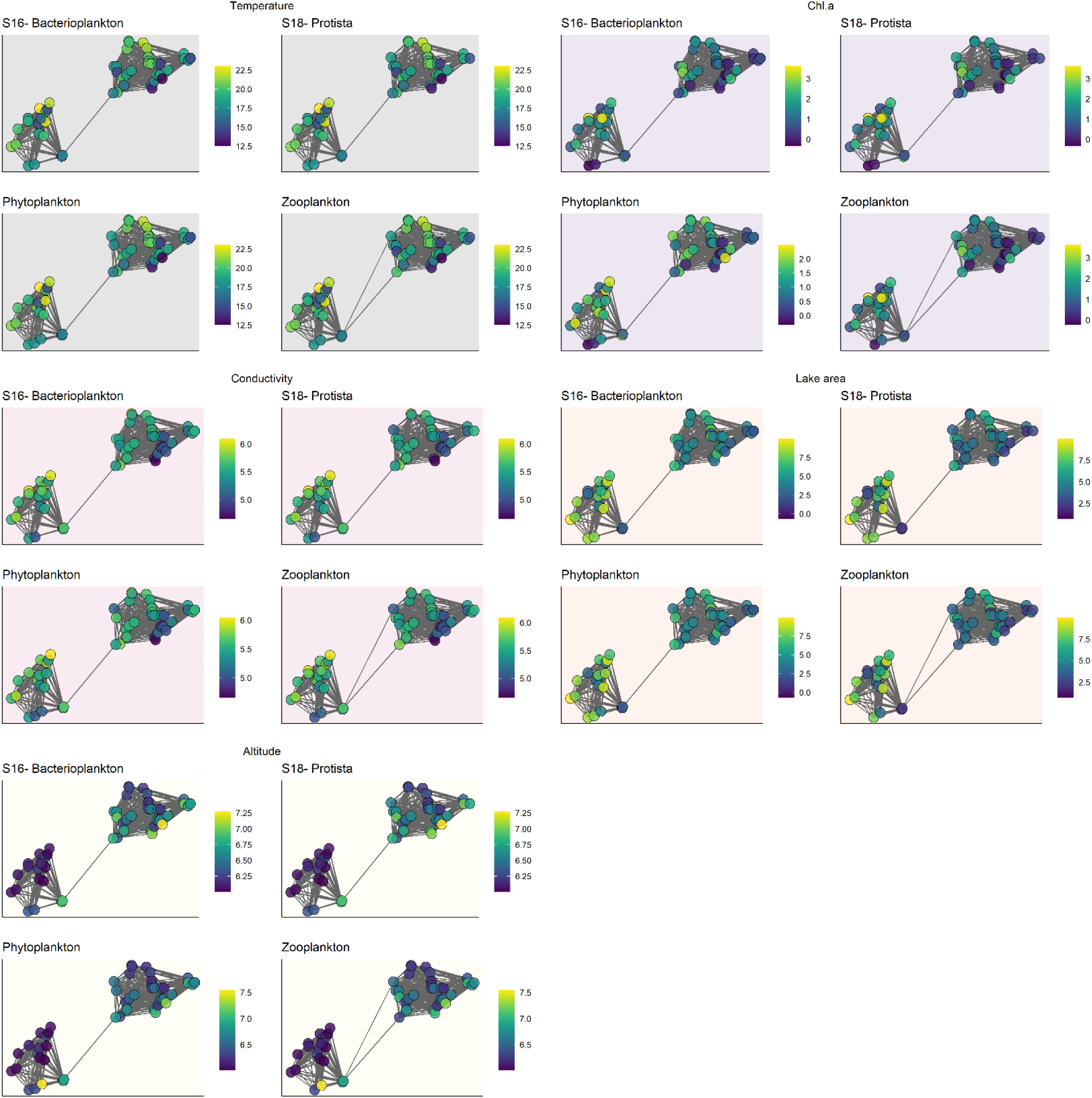
Maps representing environmental variables across lakes and for each one of the groups of organisms.

**Supplementary S2:**
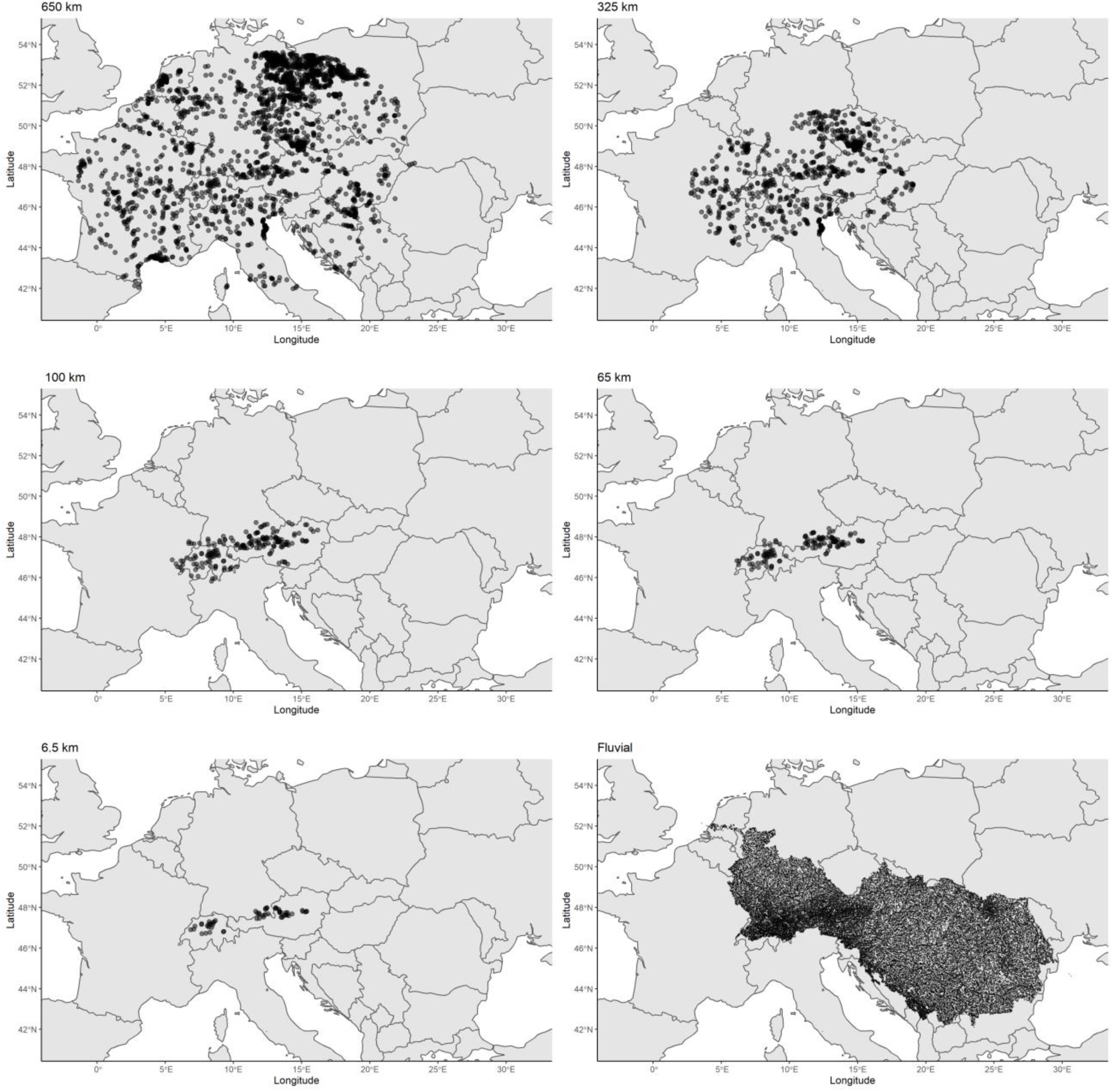
List of detected lakes for each scale considered. Lakes were subtracted from Lehner, B., and P. Do. 2004. Development and validation of a global database of lakes, reservoirs and wetlands. Journal of Hydrology 296: 1–22. doi:10.1016/j.jhydrol.2004.03.028. Fluvial network was subtracted from on Lehner, B., K. Verdin, and A. Jarvis. 2008. New global hydrography derived from spaceborne elevation data. Eos, Transactions, American Geophysical Union 89: 93–94. For the fluvial network, each point corresponds to a river reach where lakes were snapped.

**Supplementary S3:**
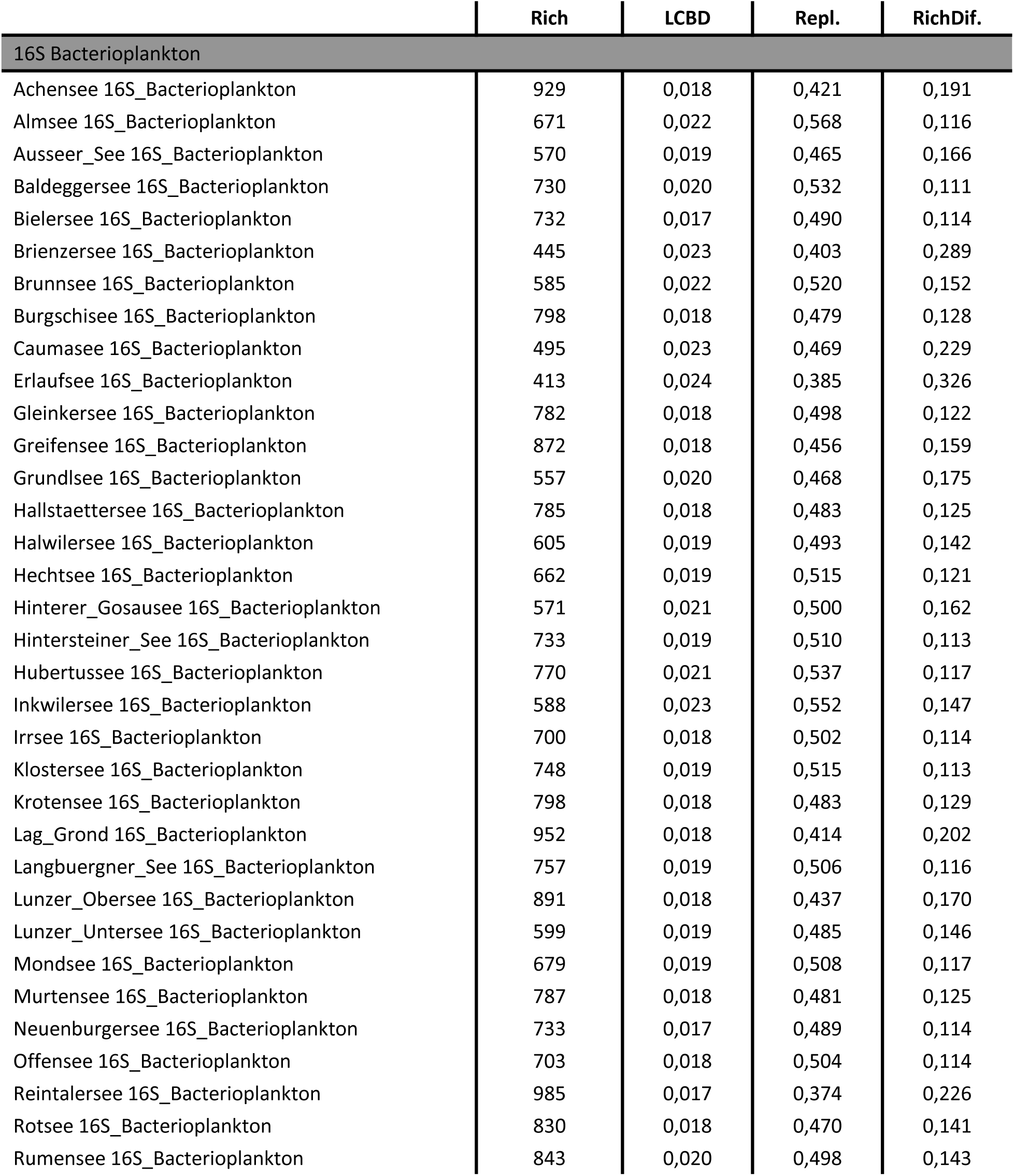
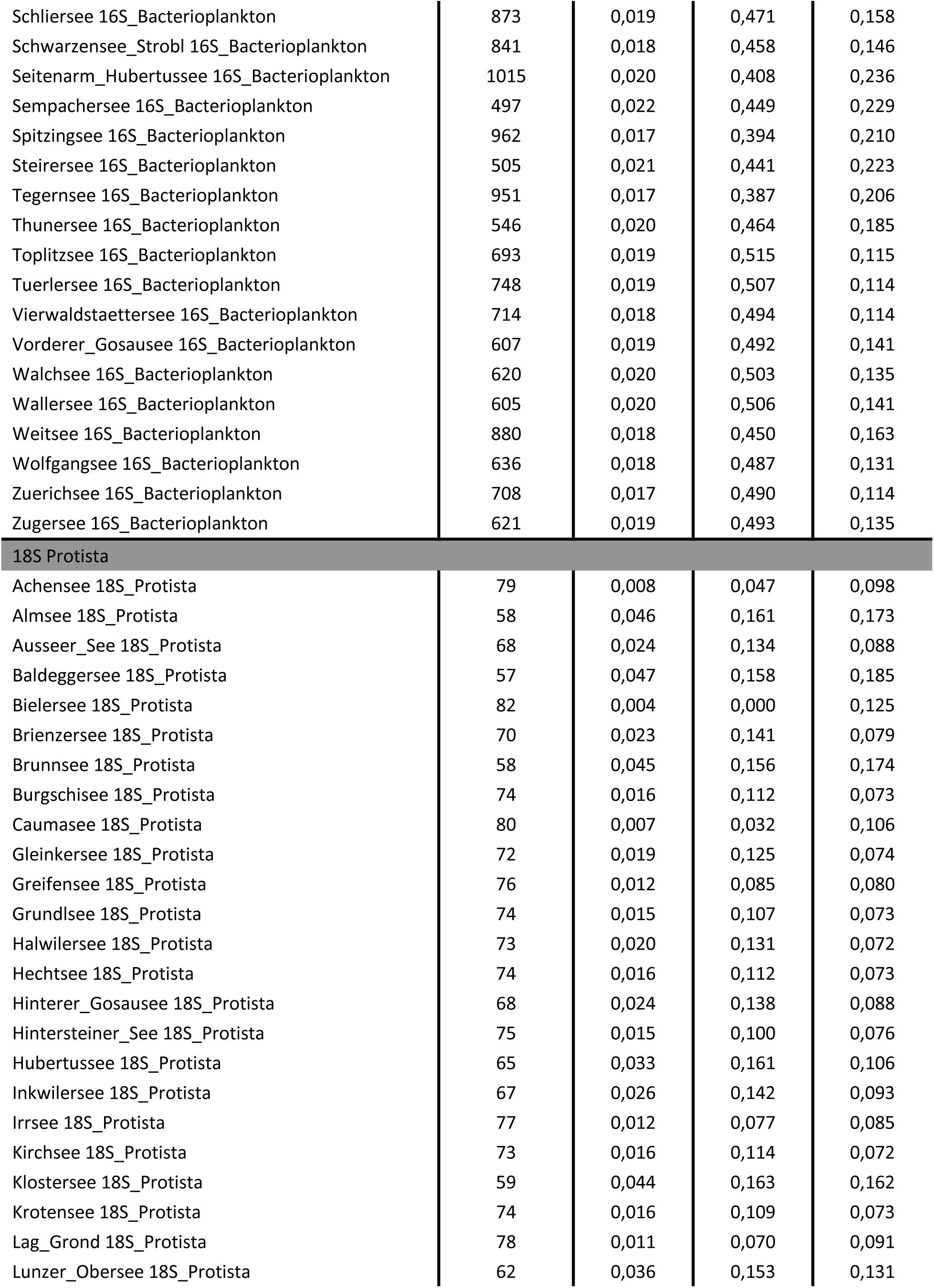
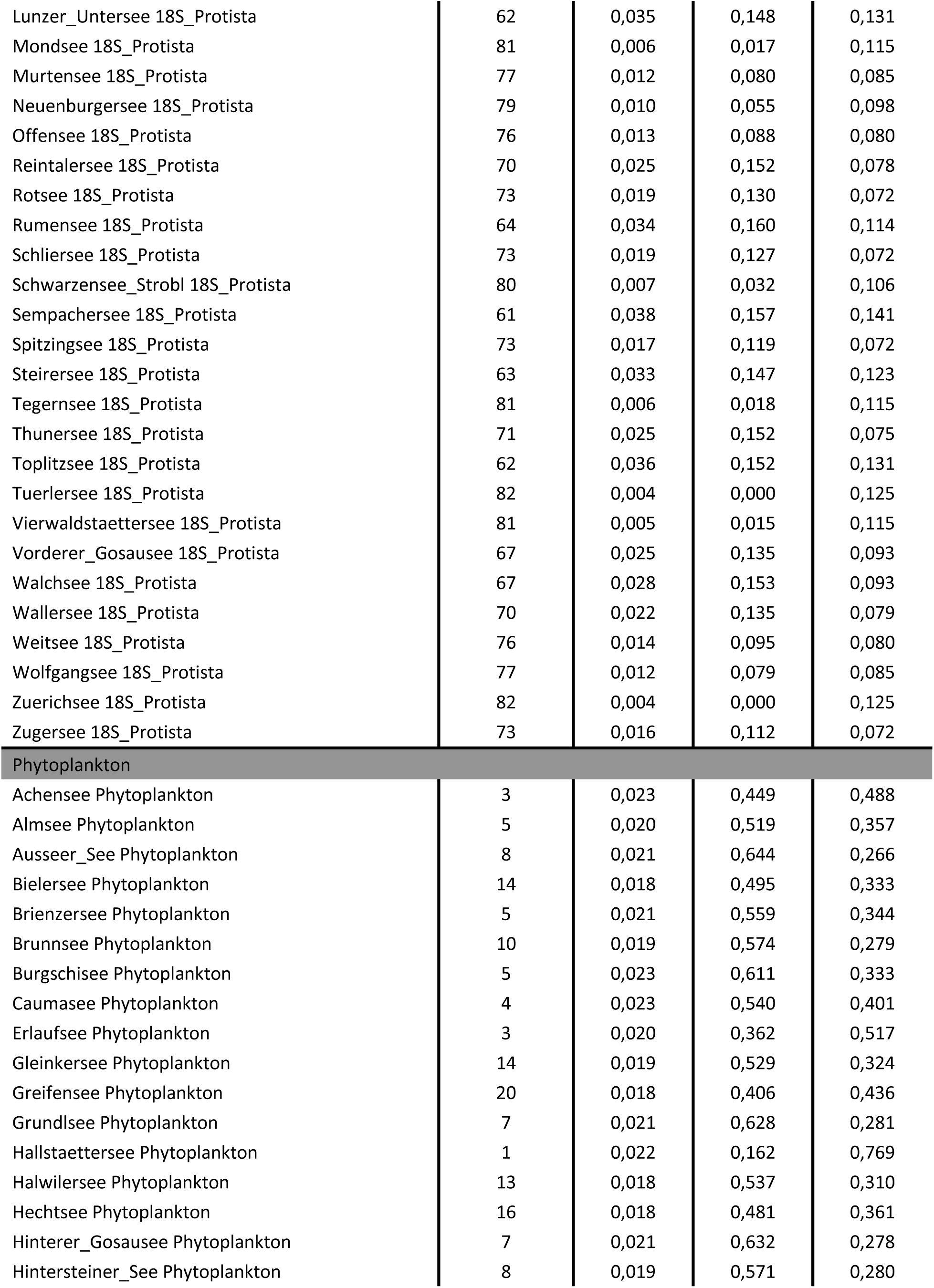
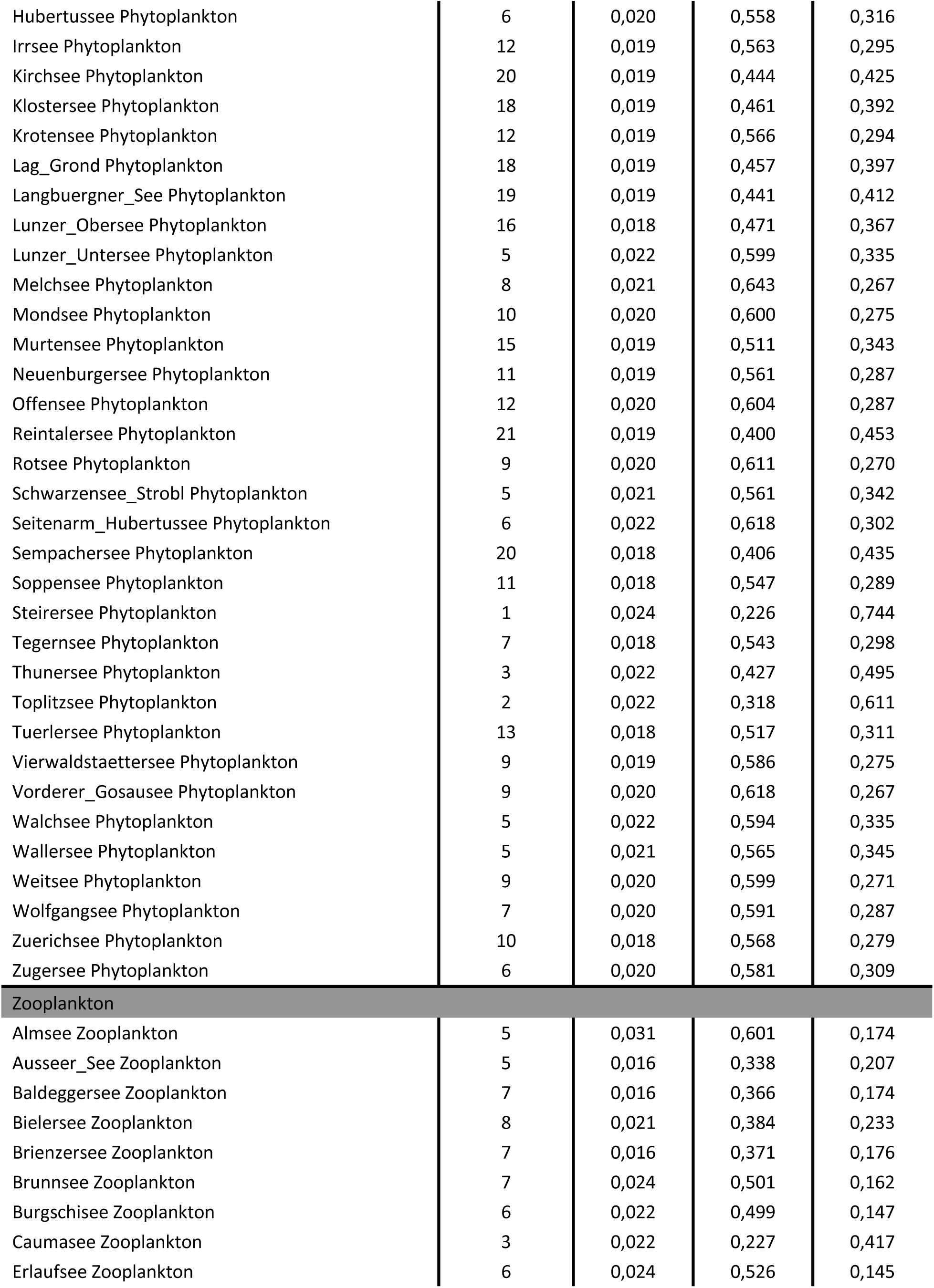
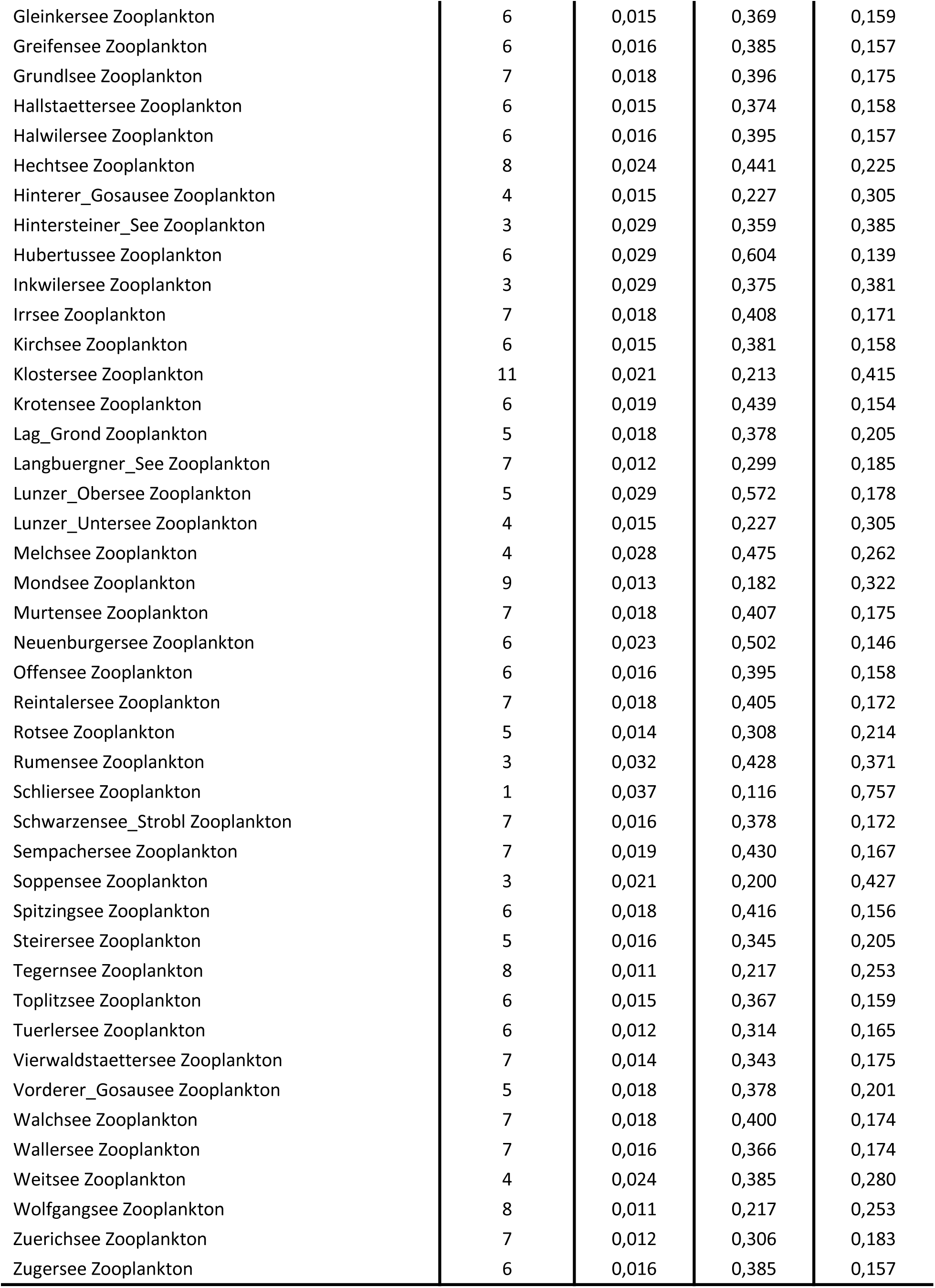
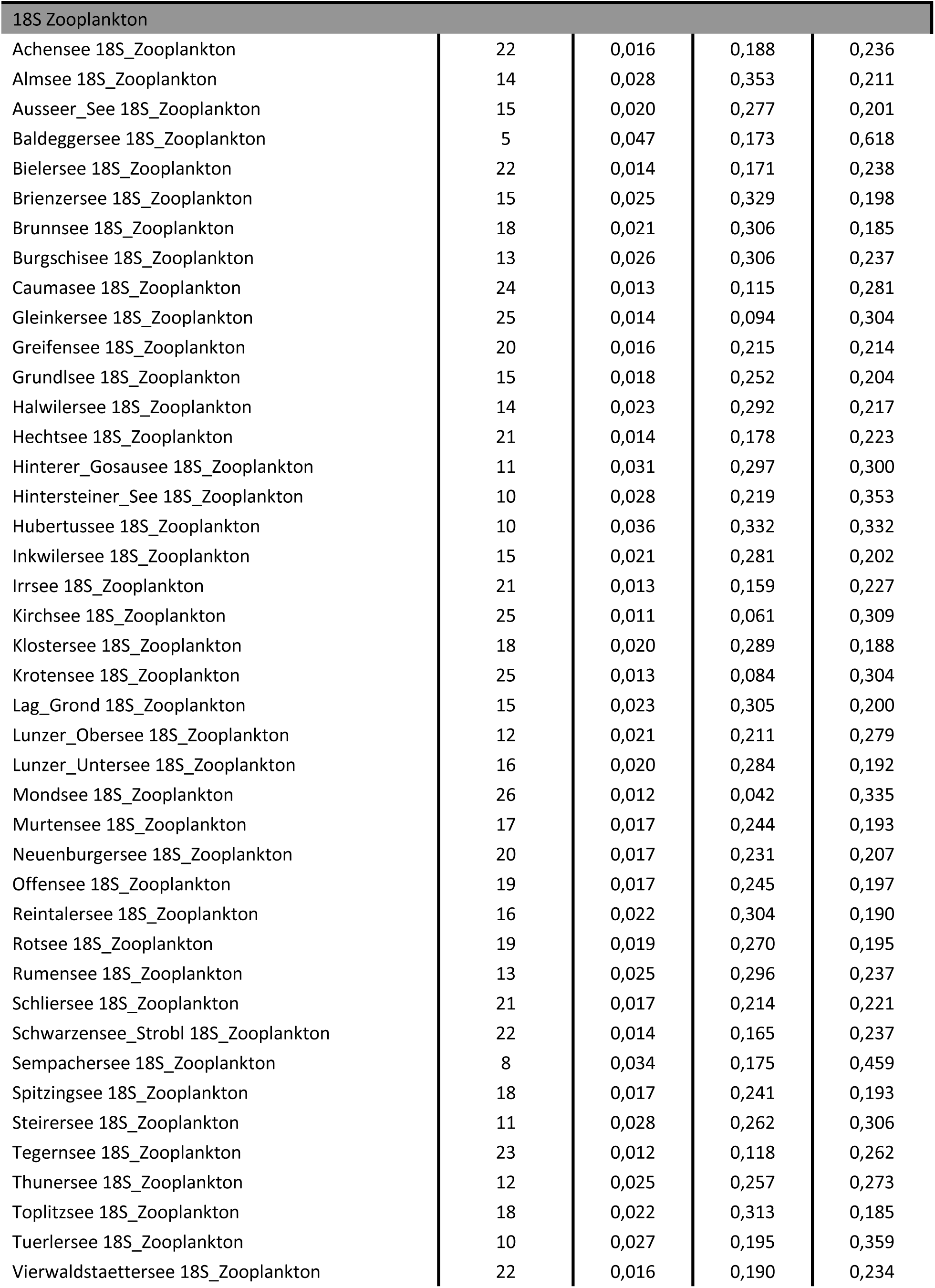
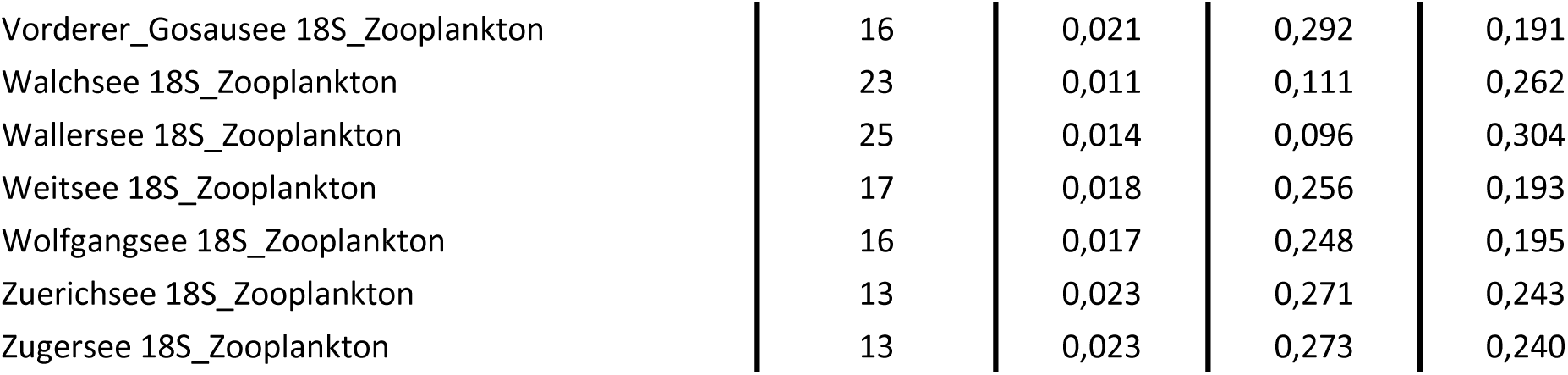
Community structure values for all the organisms’ groups (grey rows). *Rich* corresponds to the species richness, *LCBD* to the local contribution to beta diversity, *Repl* to the average site replacement and *RichDif* to average site richness difference. Both *Repl* and *RichDif* correspond to the beta diversity partitions calculated with Jaccard dissimilarity index.

**Supplementary S4:**
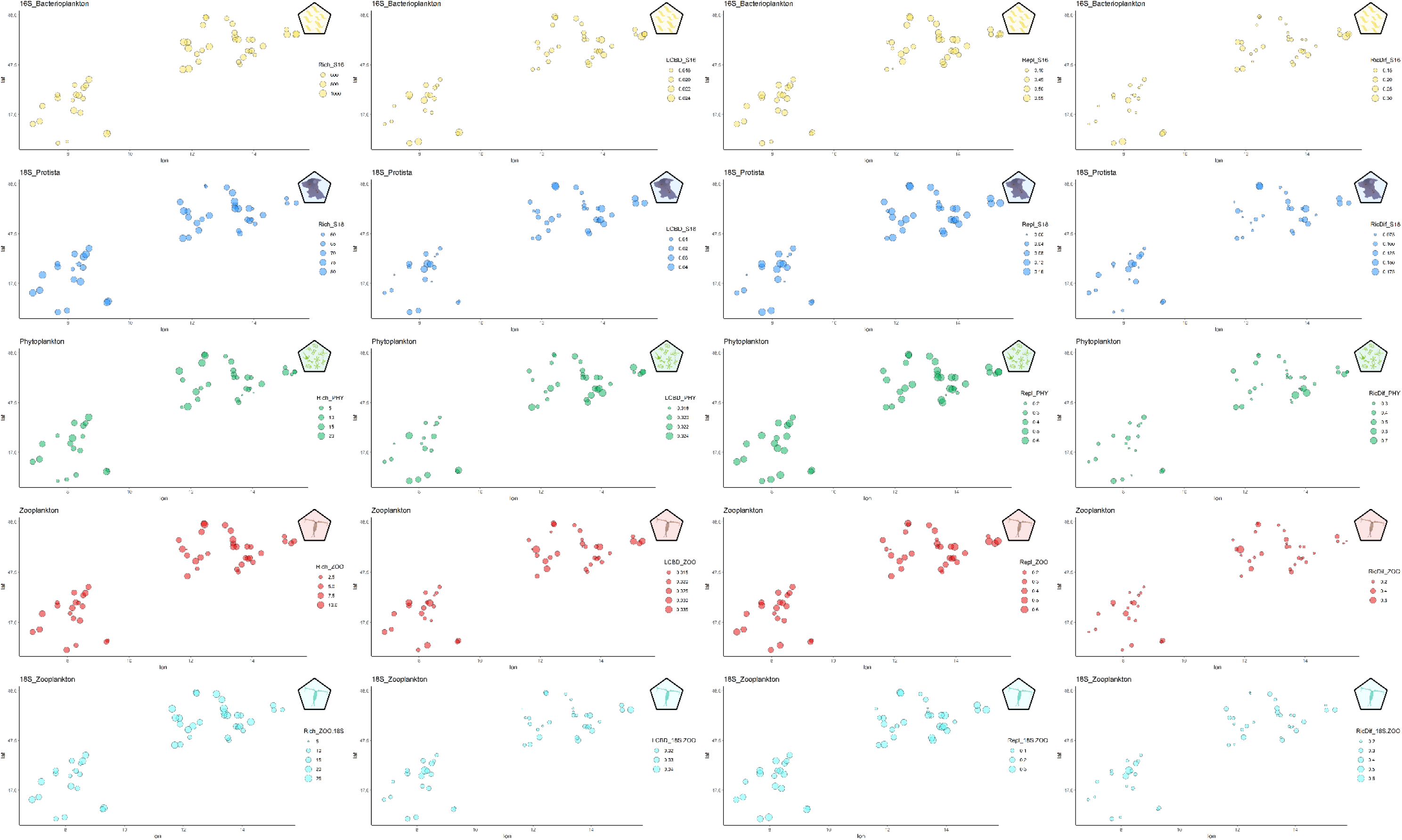
Plots showing diversity metrics distribution for each one of the studied groups. Size indicates metrics values. Colours correspond to the different groups: bacterioplankton, protist, phytoplankton, zooplankton and 18S zooplankton respectively.

**Supplementary S5:**
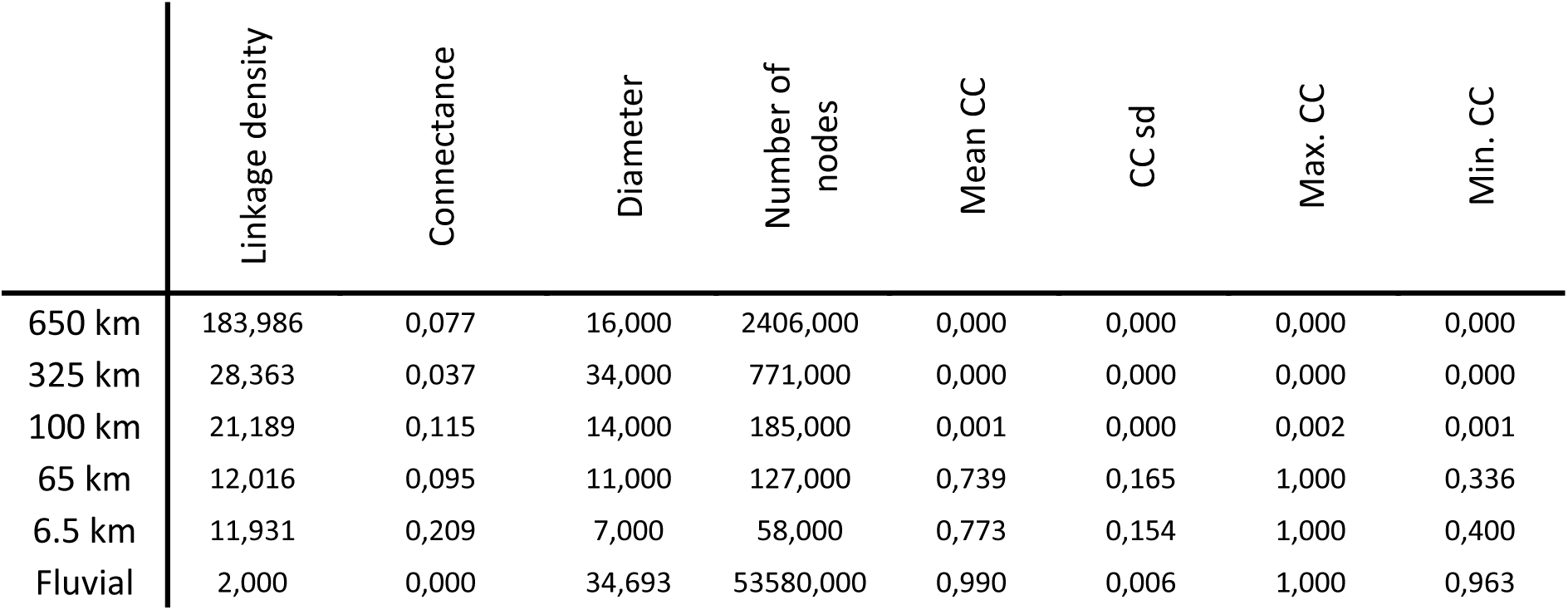
Network descriptive parameters calculated for each one of the constructed networks. Note that Fluvial network corresponds to a dendritic network and some of the values (e.g. degree) must be understood correspondingly.

**Supplementary S6:**
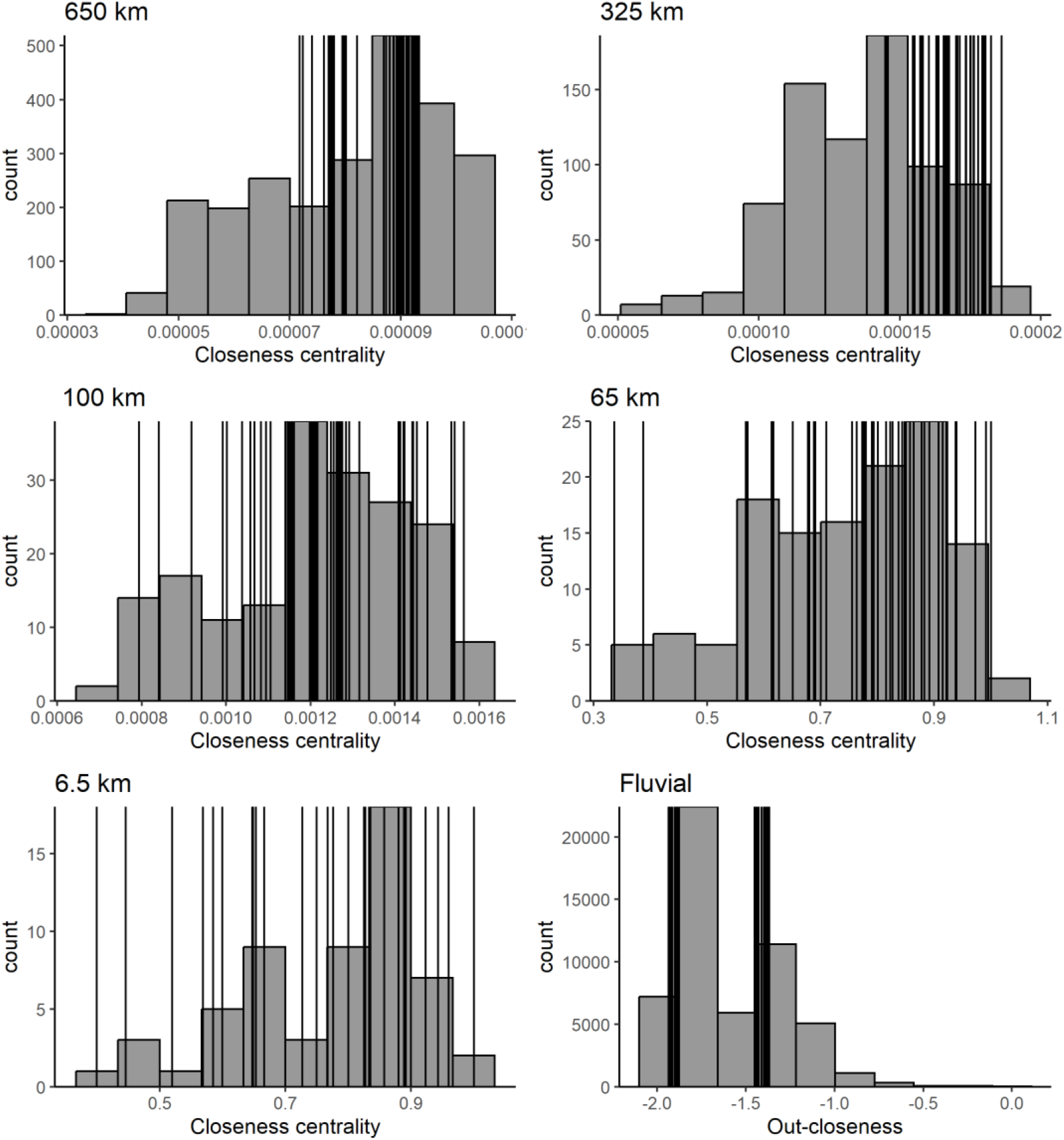
Network centrality values distribution within each one of the constructed networks. Black lines correspond to the sampled lakes that appear well distributed along the range of centrality values capturing its main variability or a wide part of it.

**Supplementary S7:**
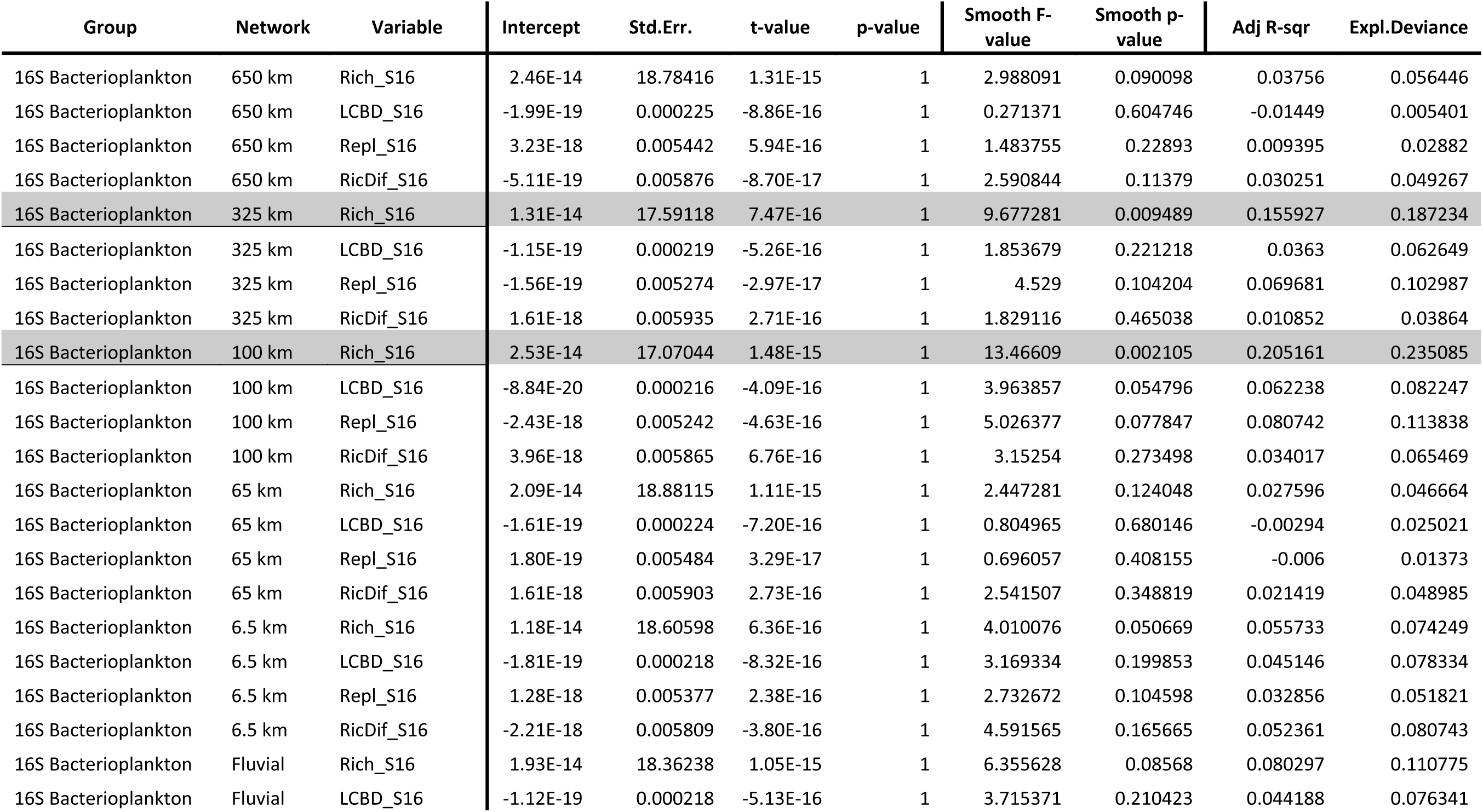
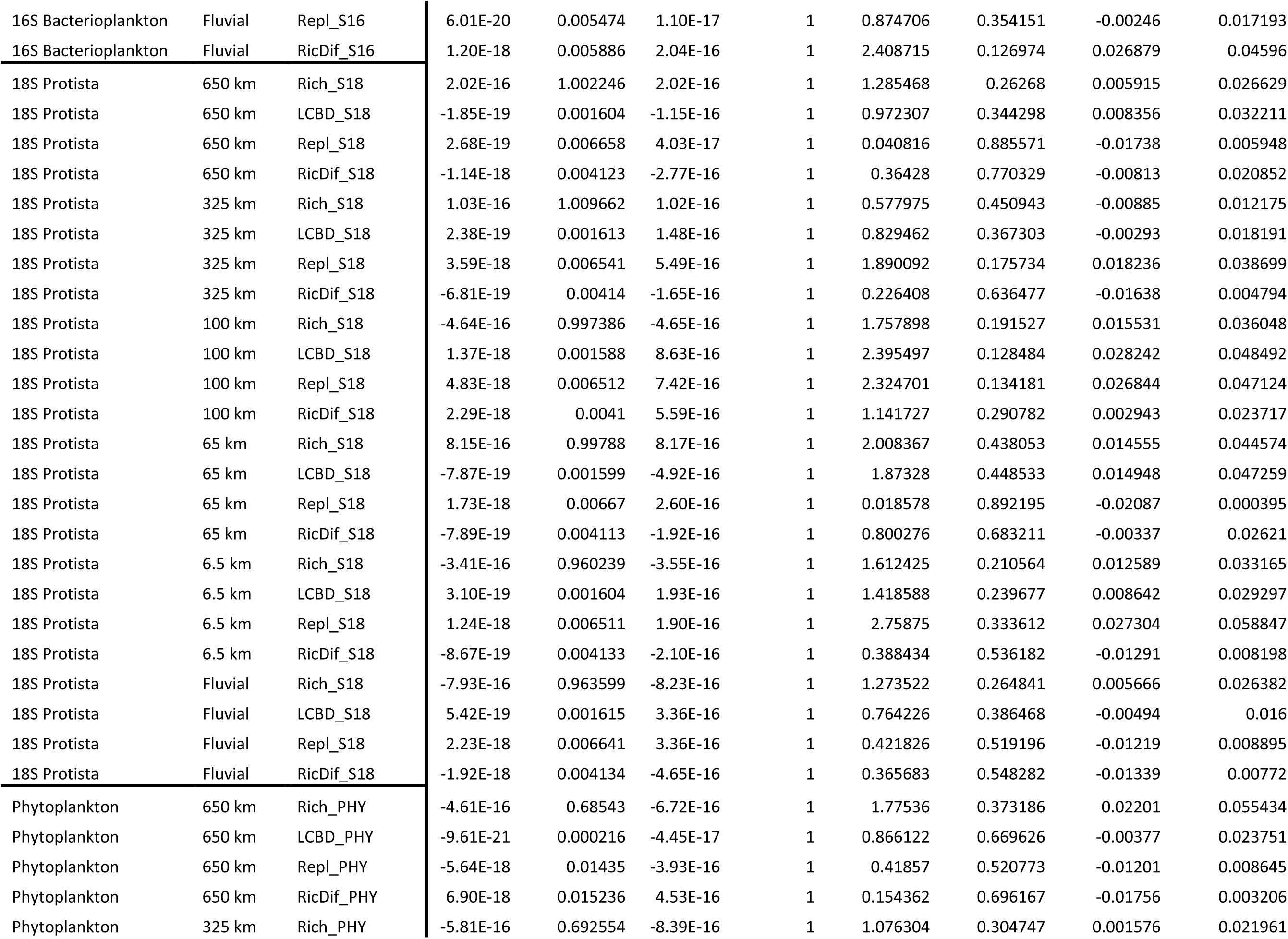
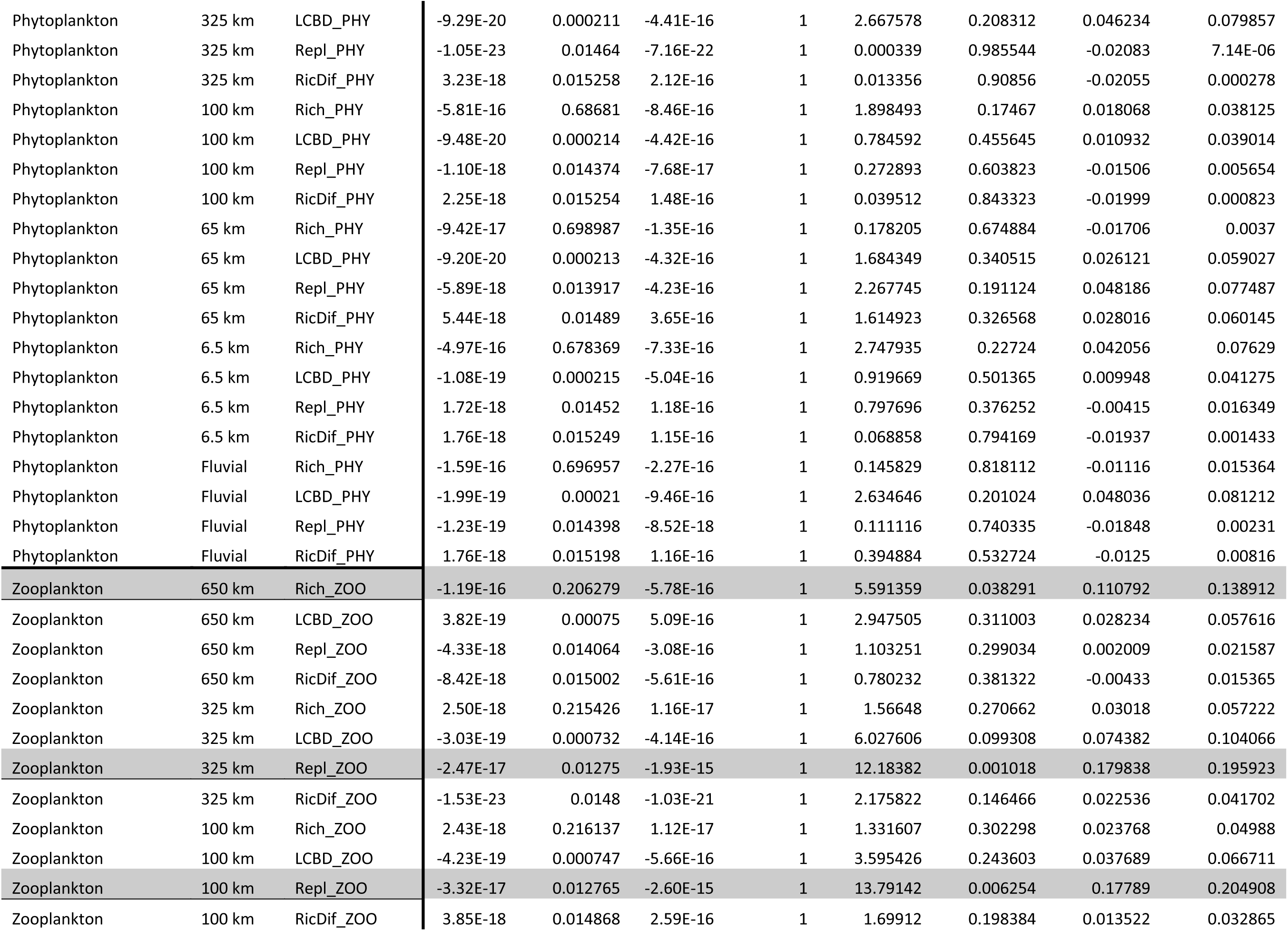
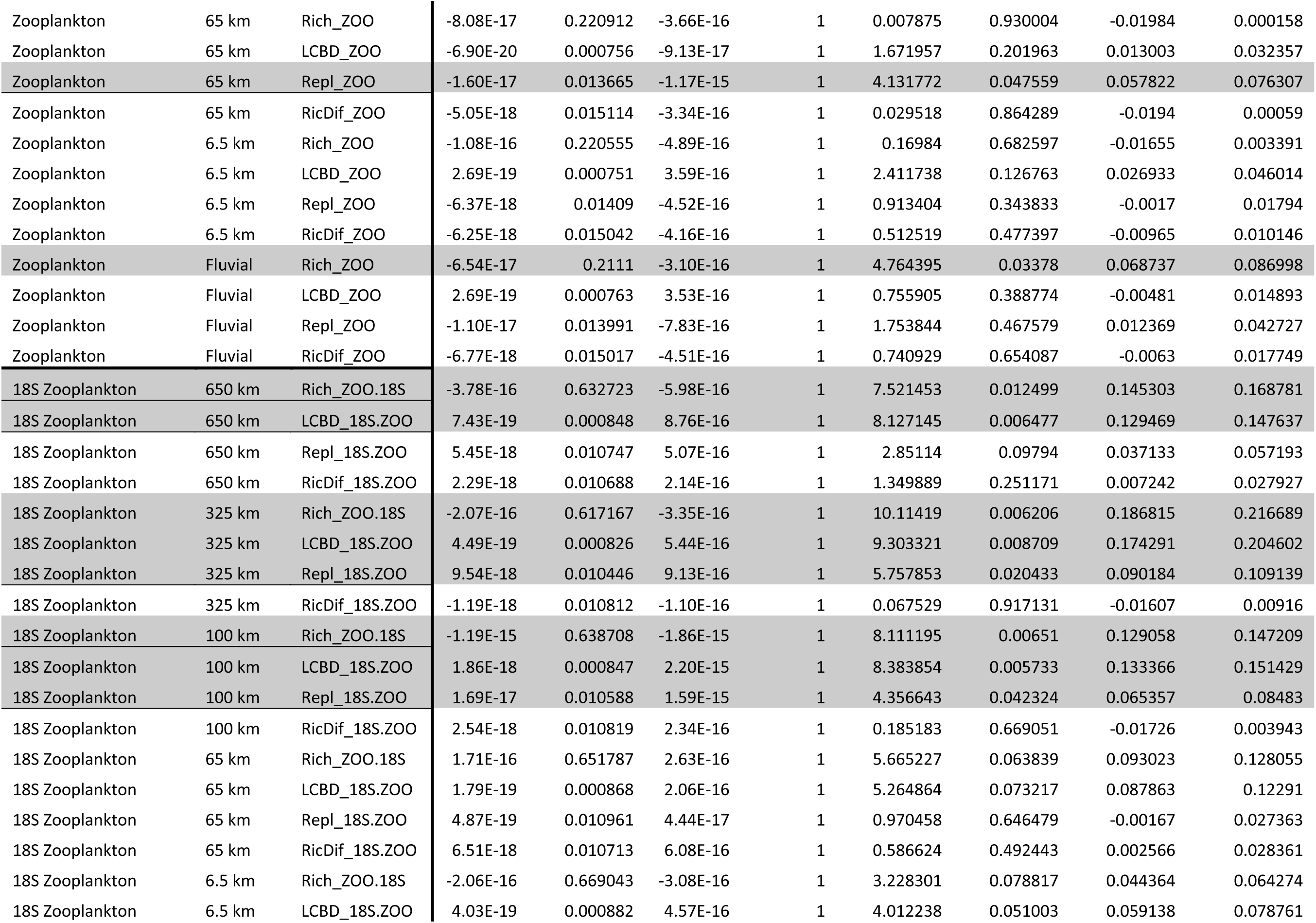
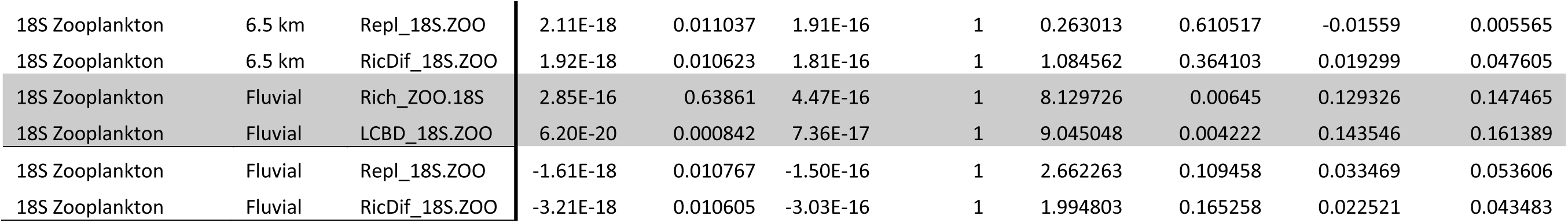
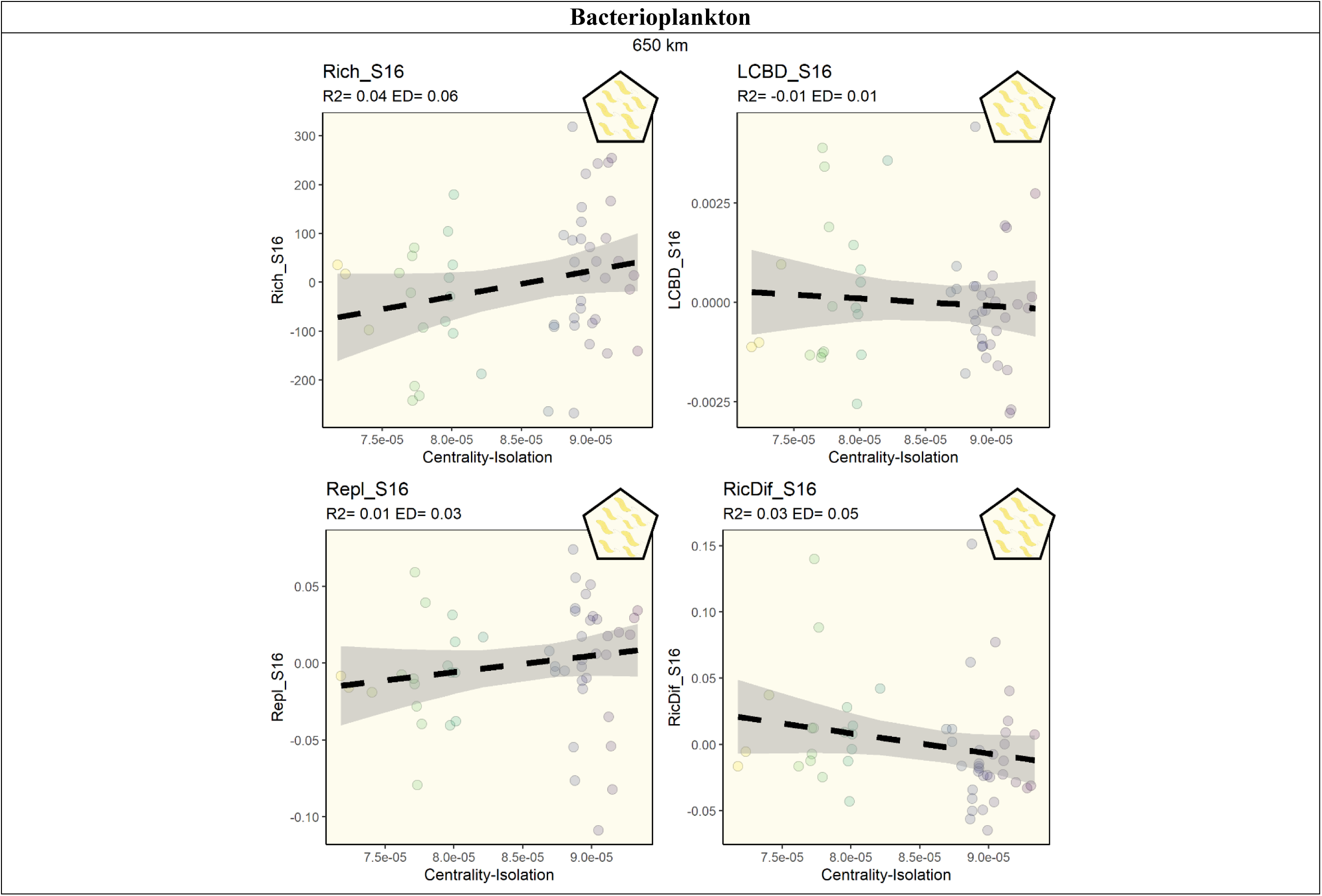
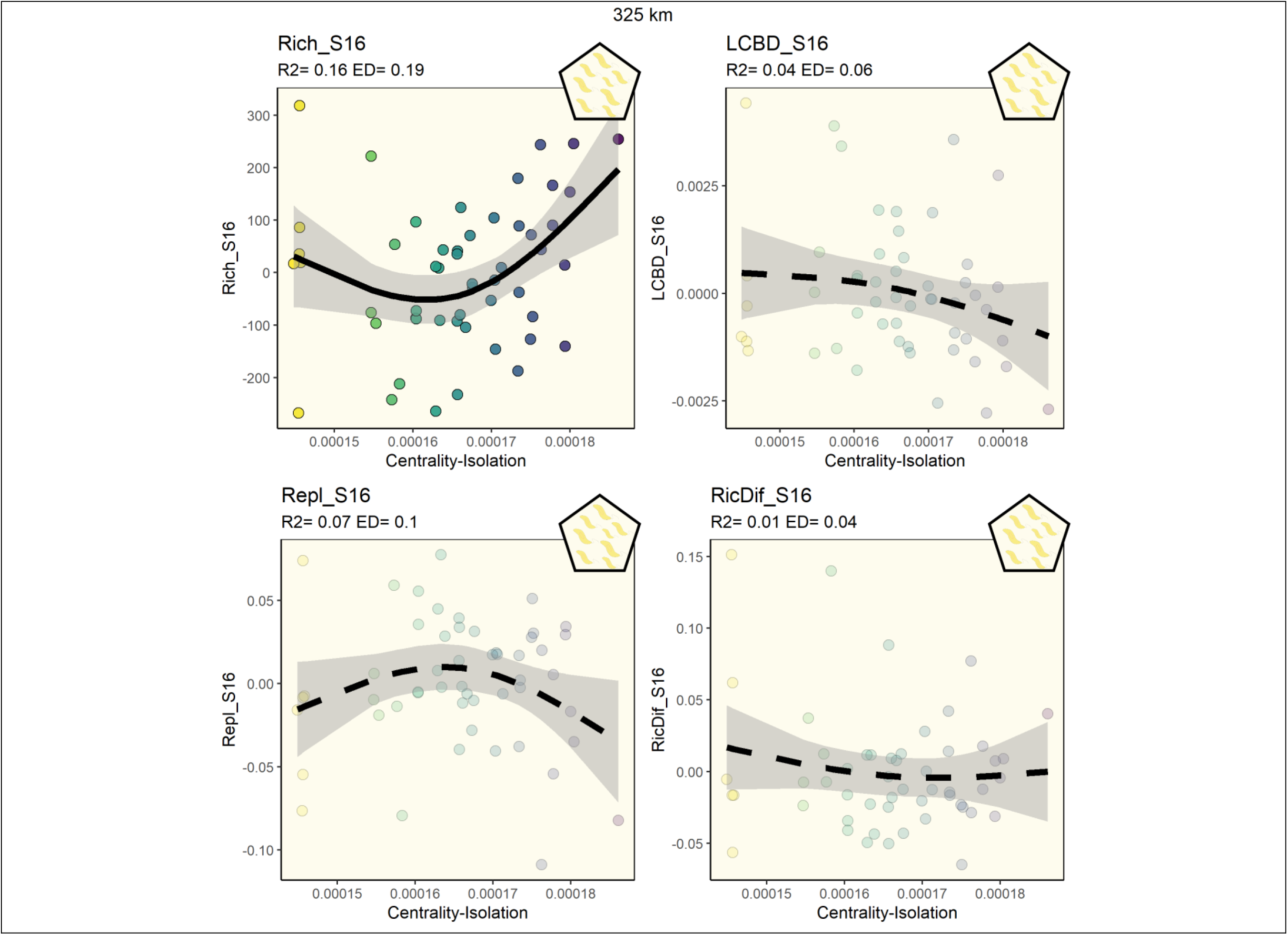
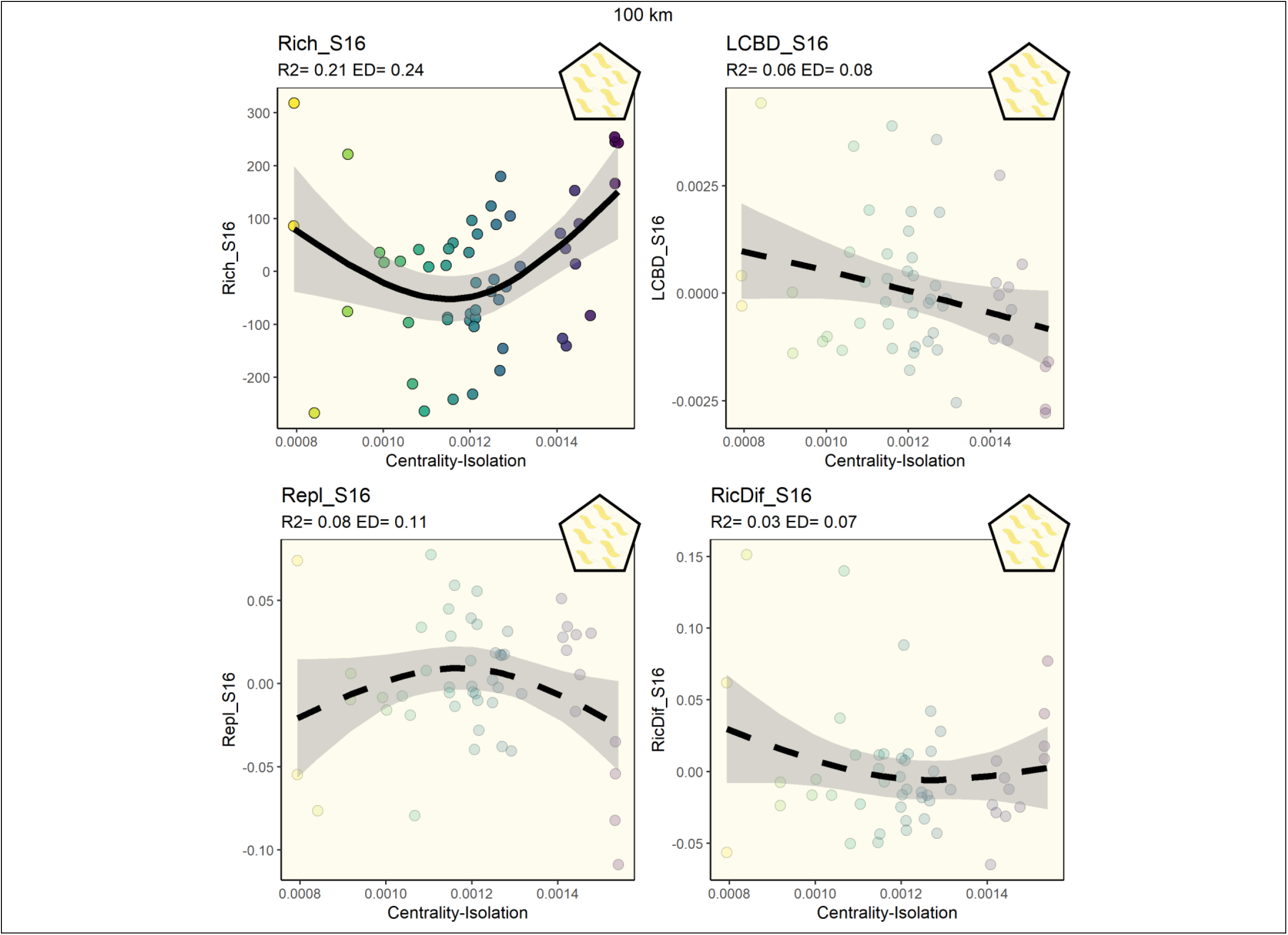
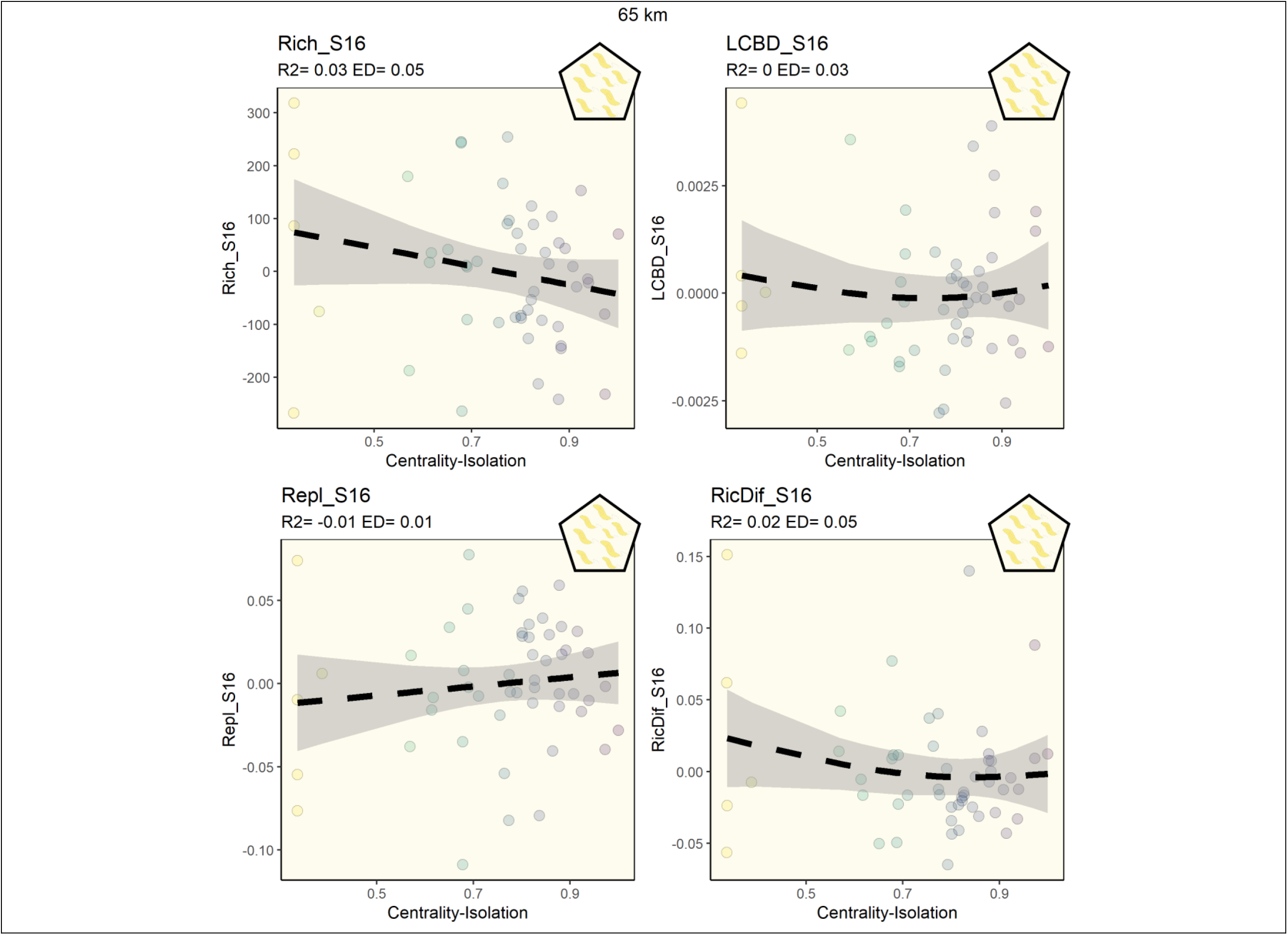
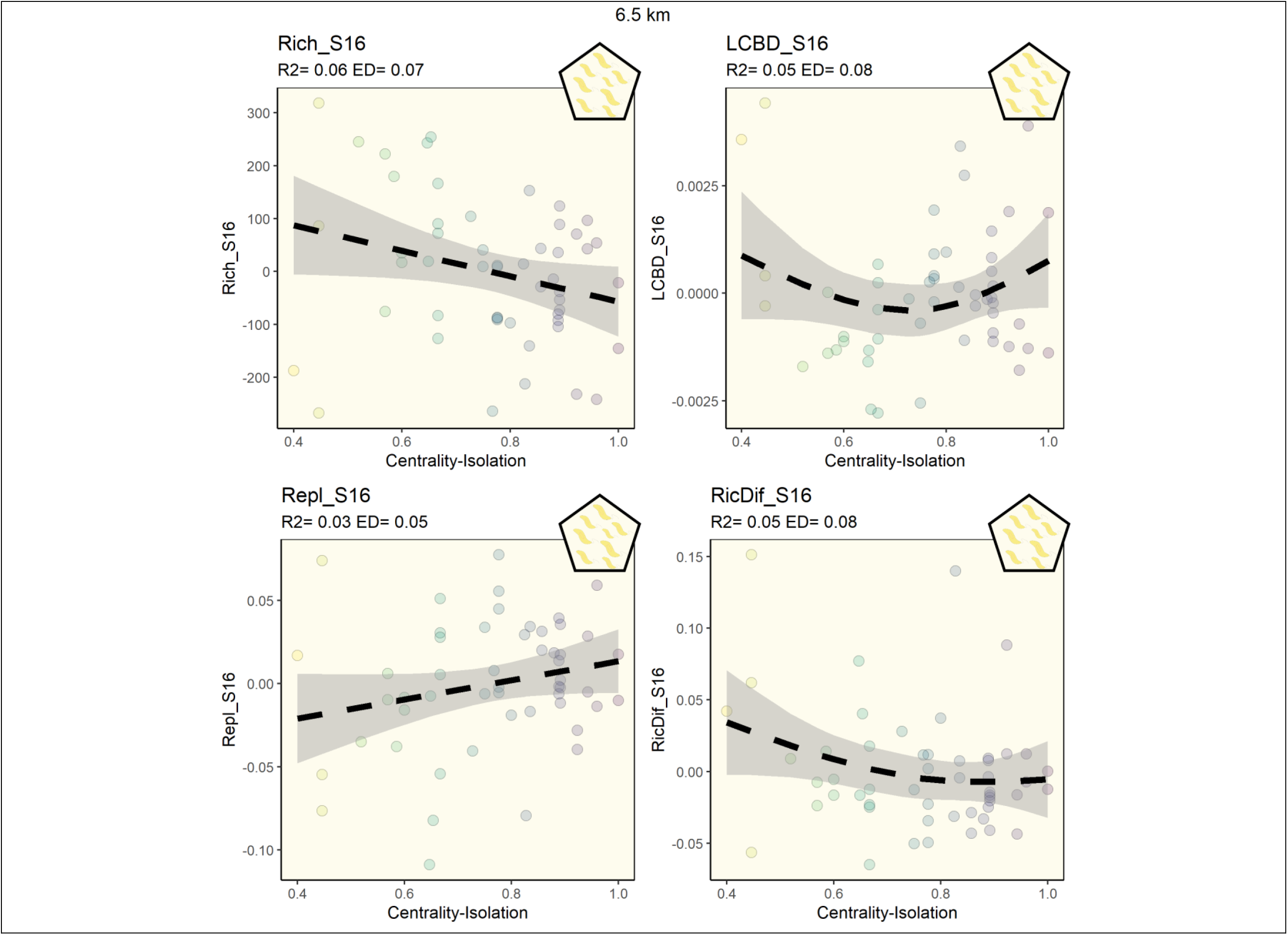
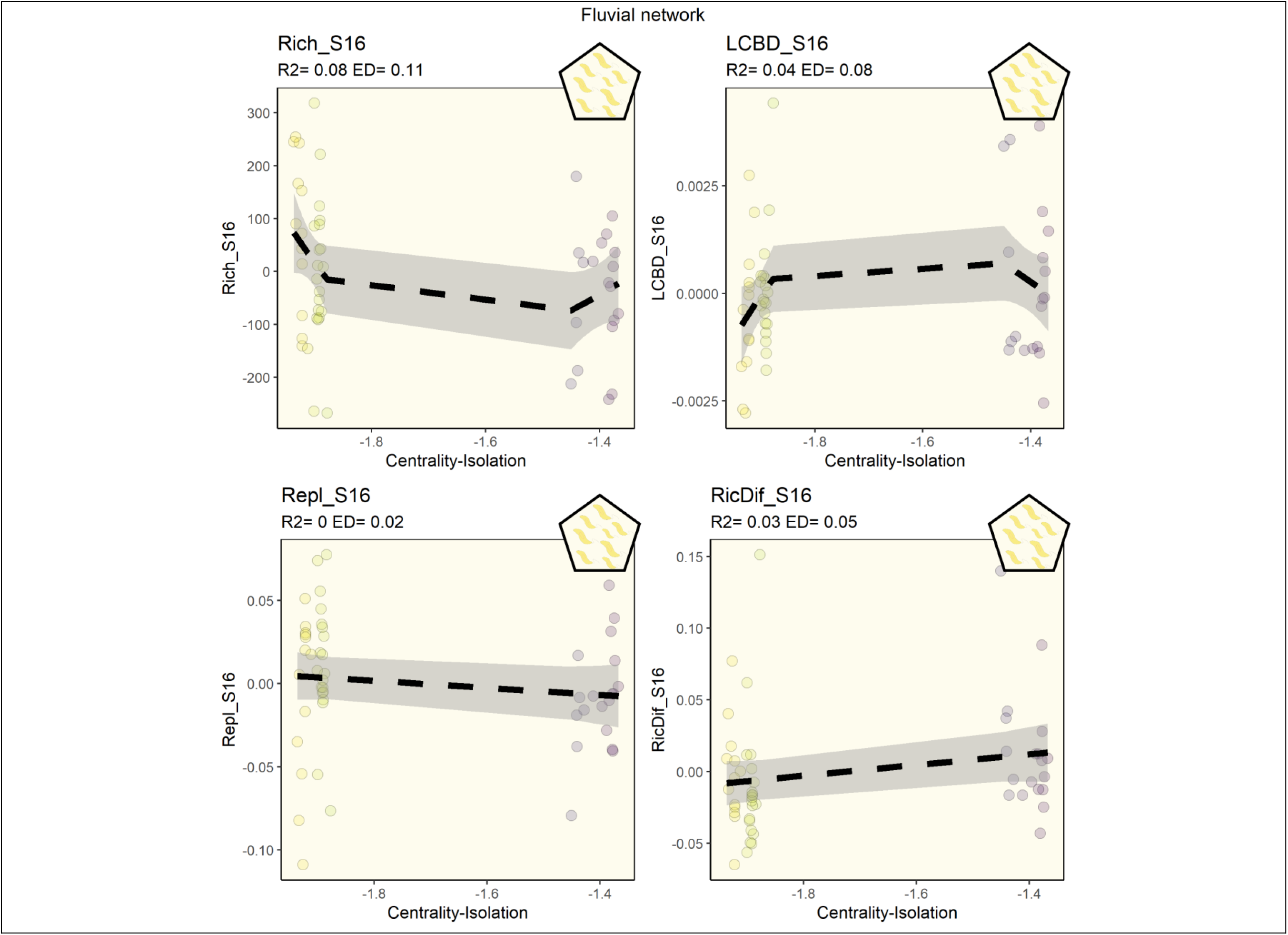
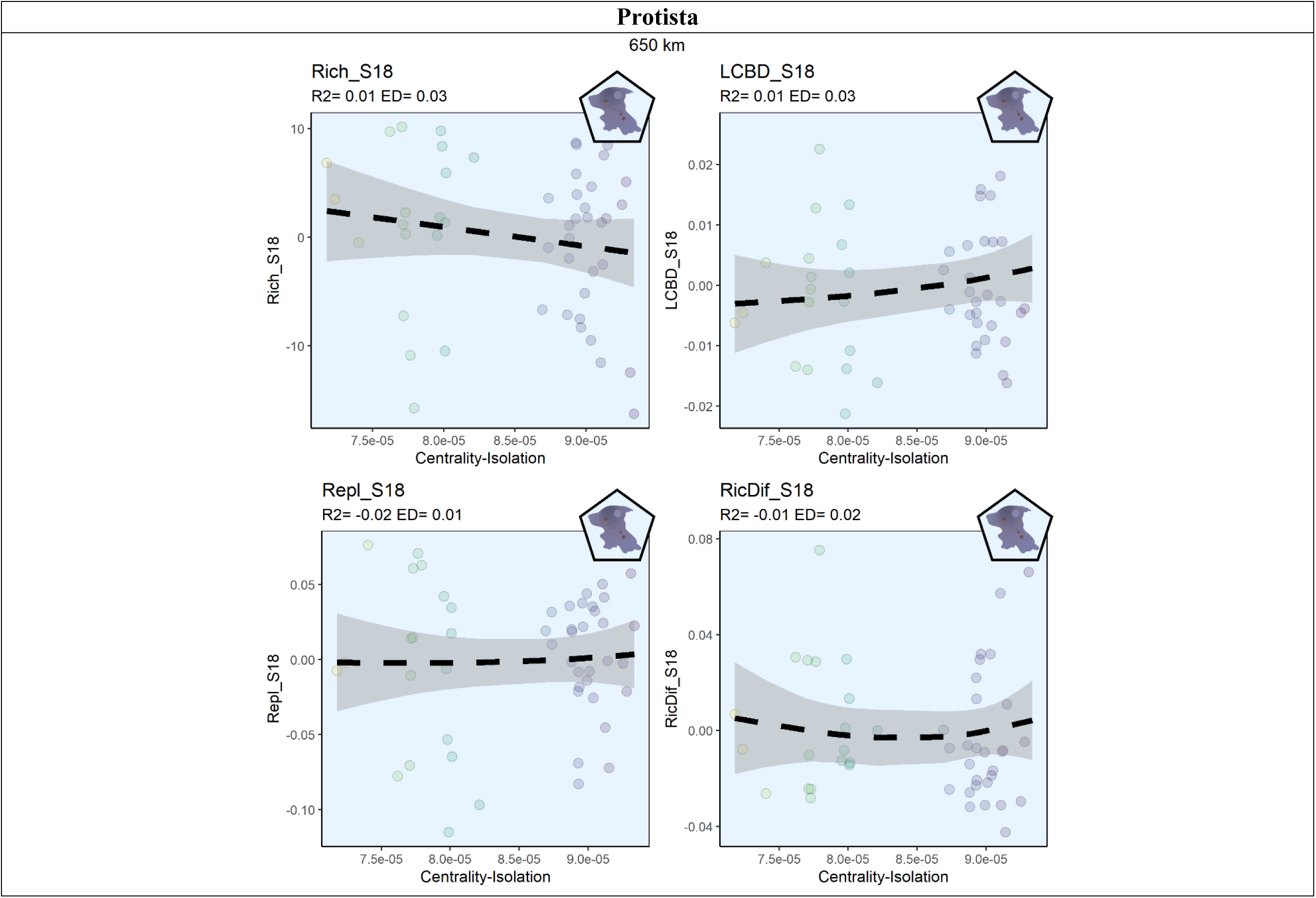
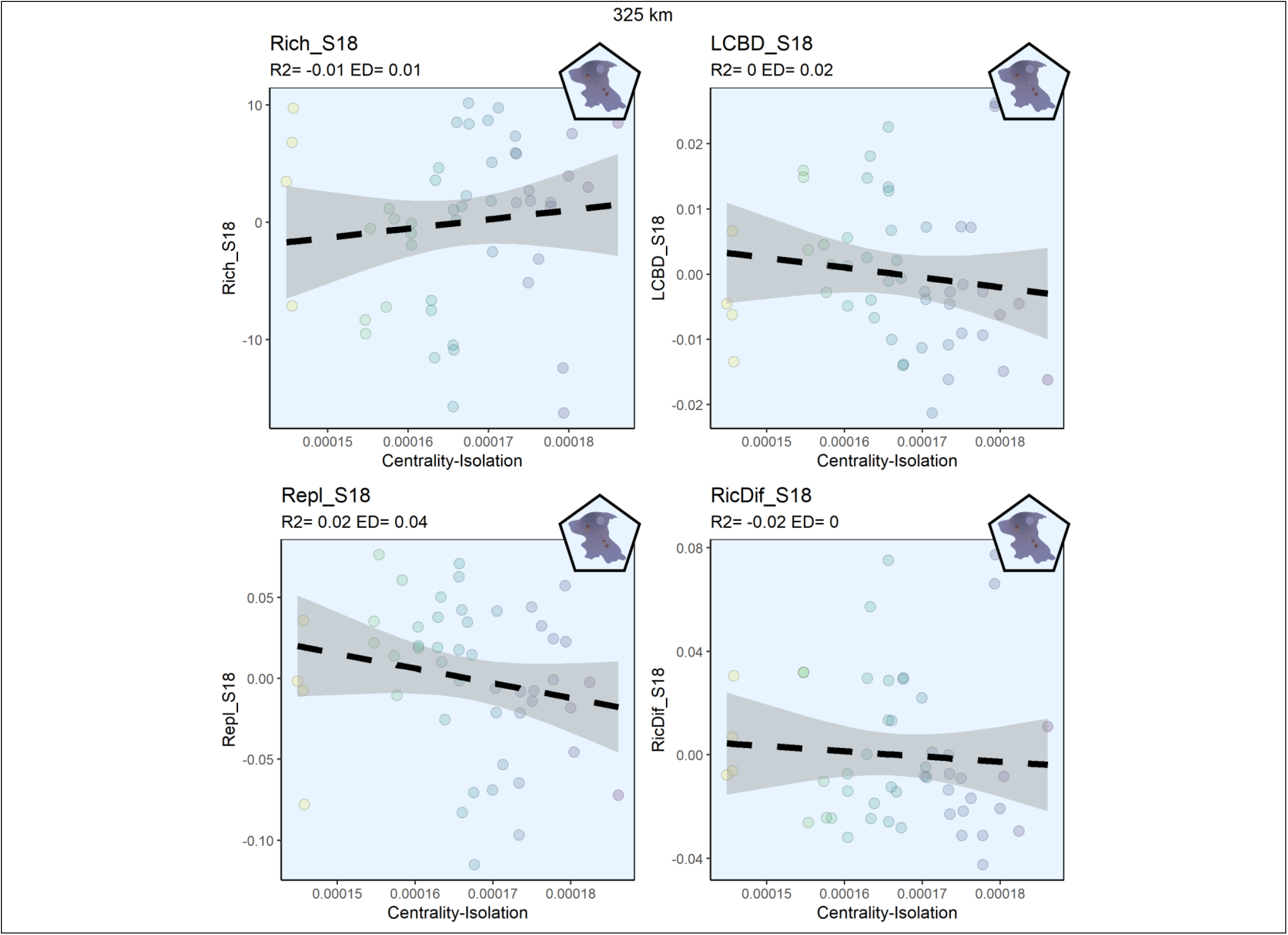
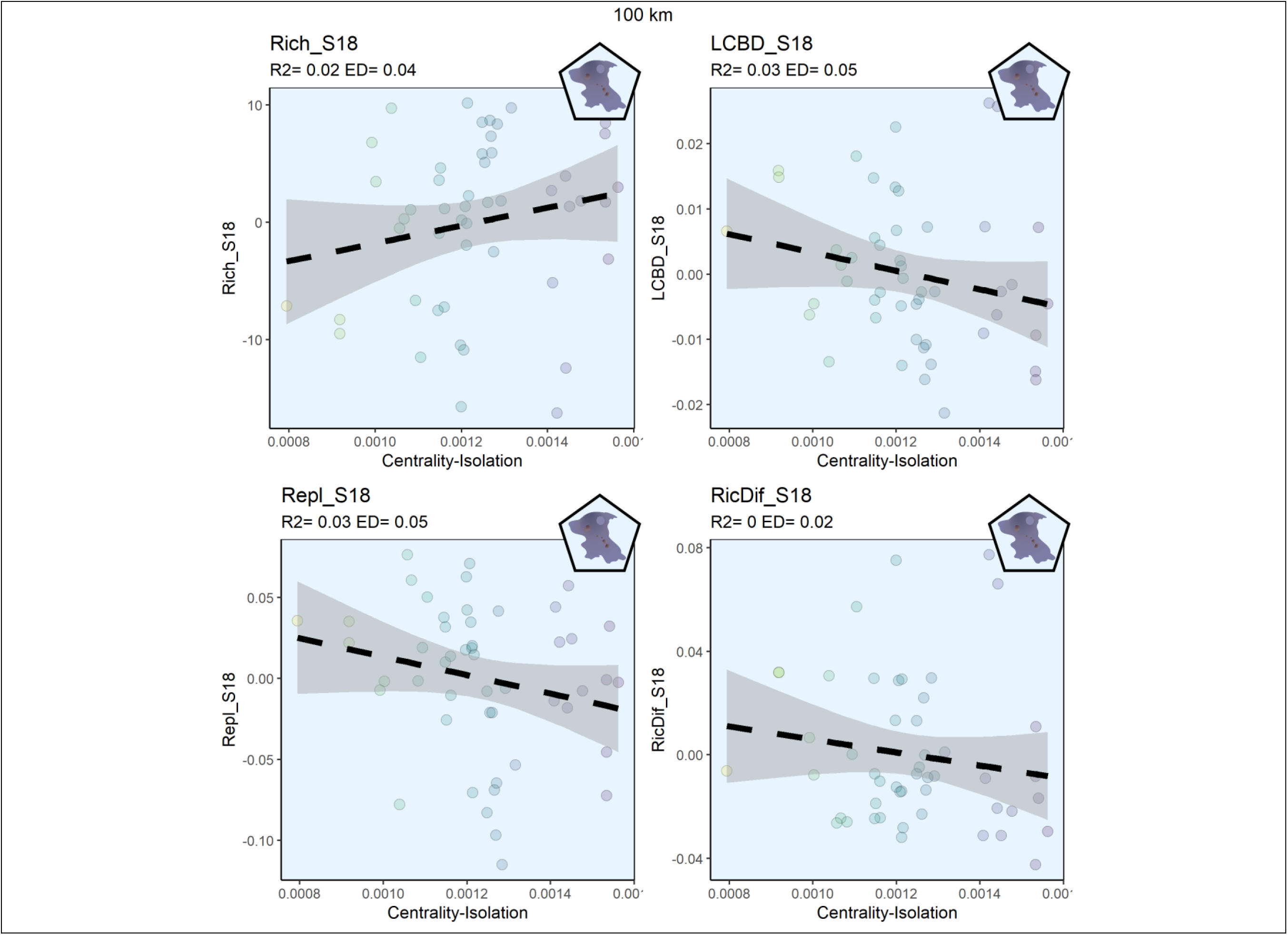
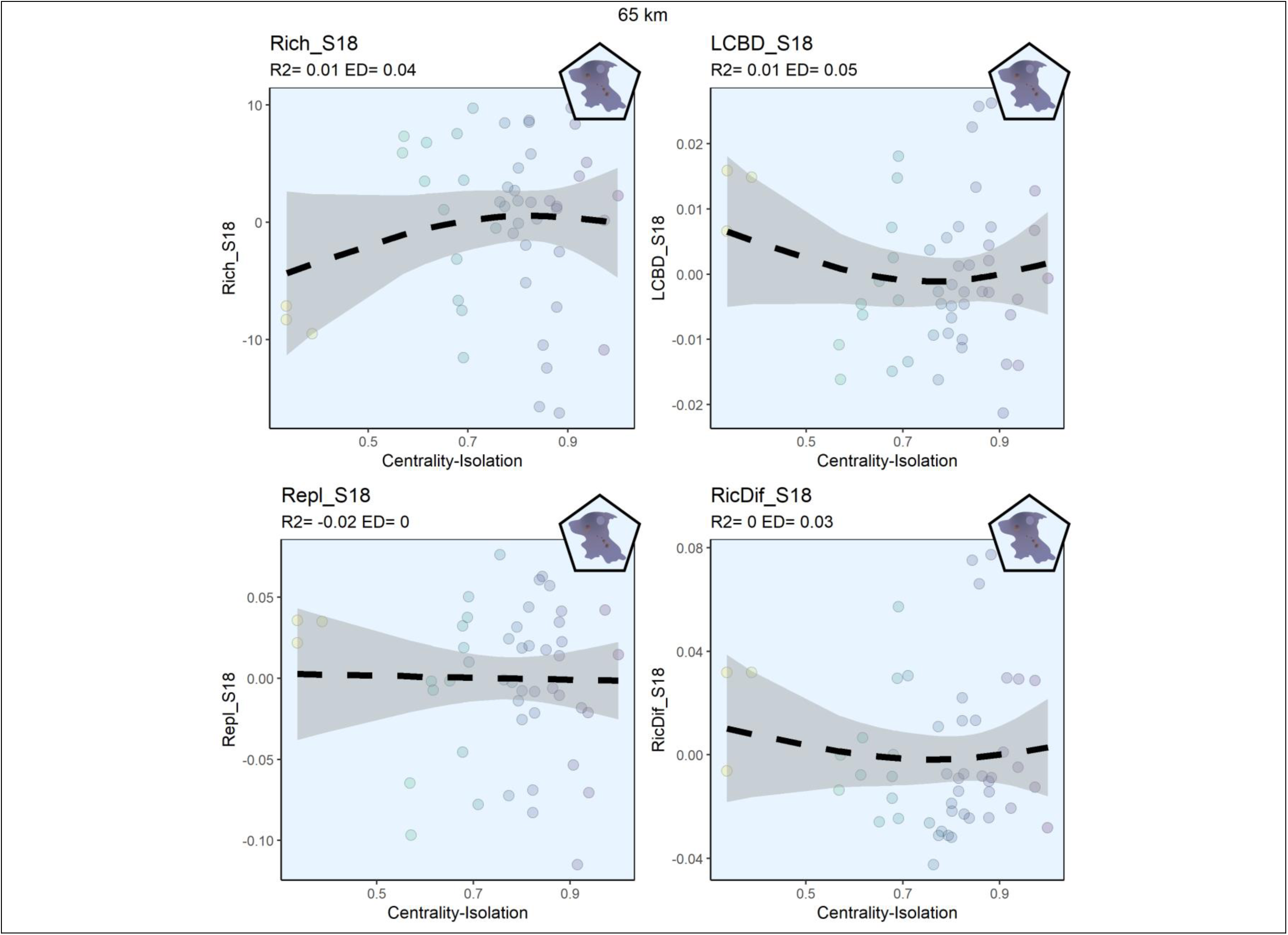
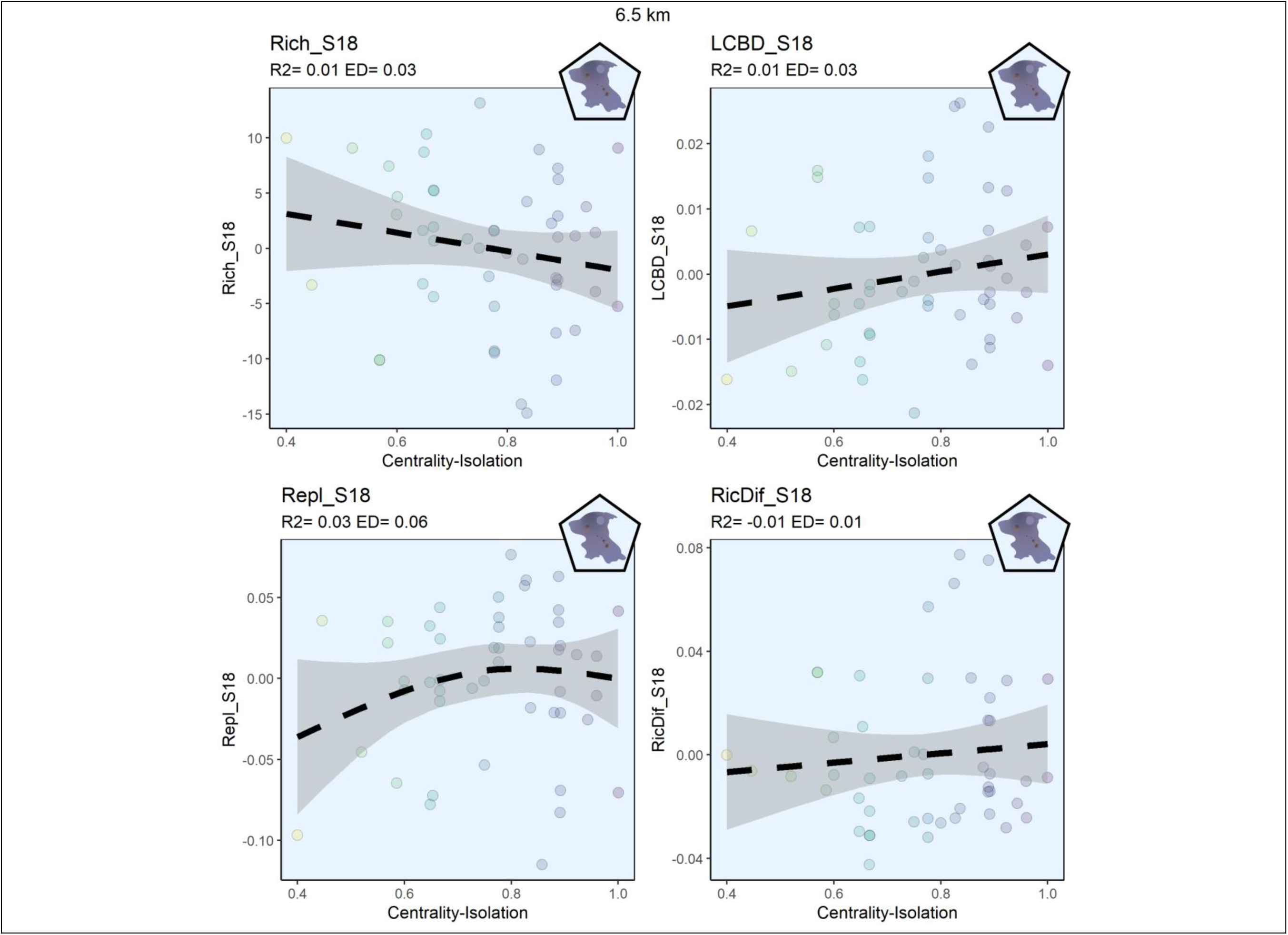
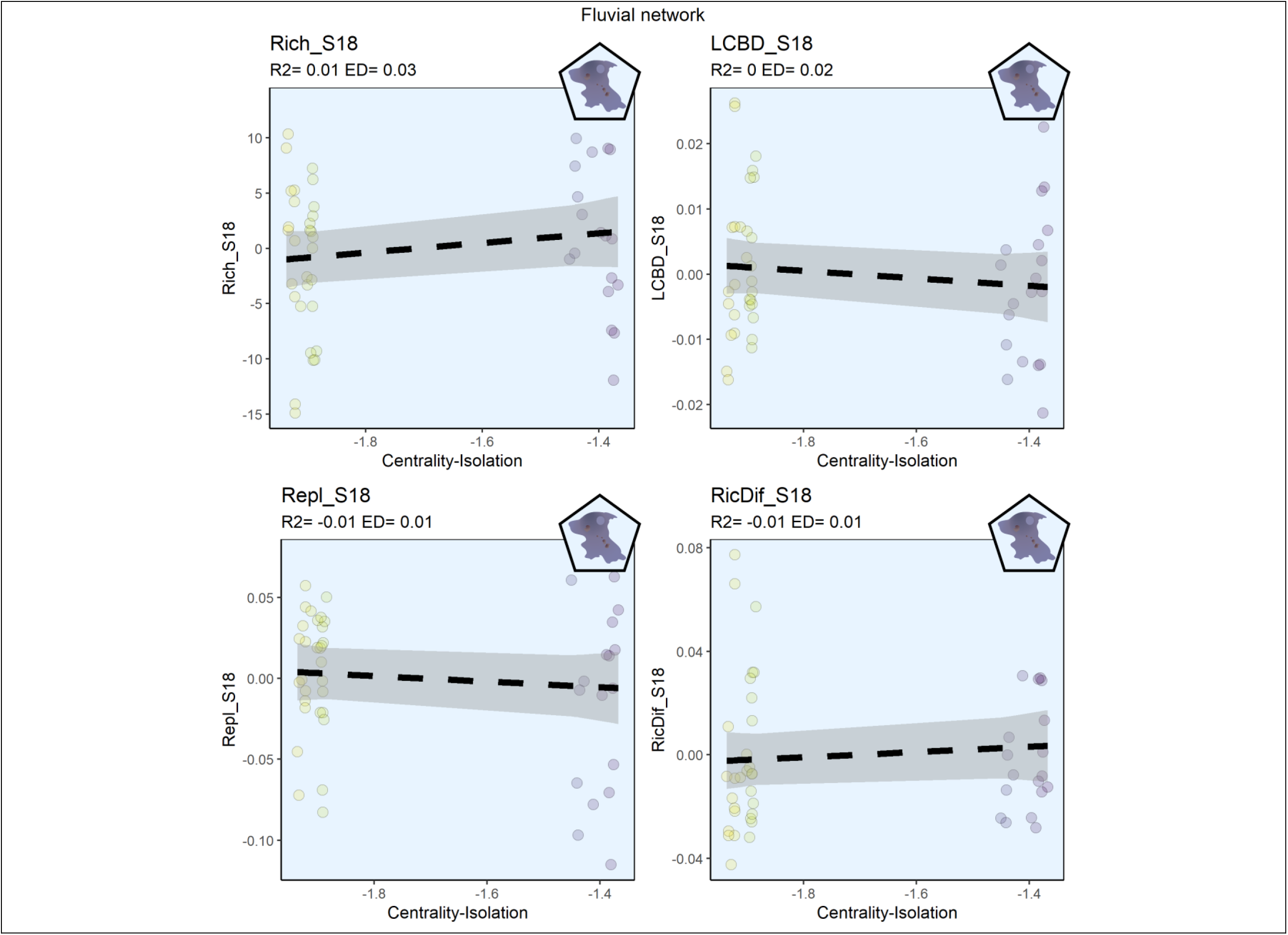
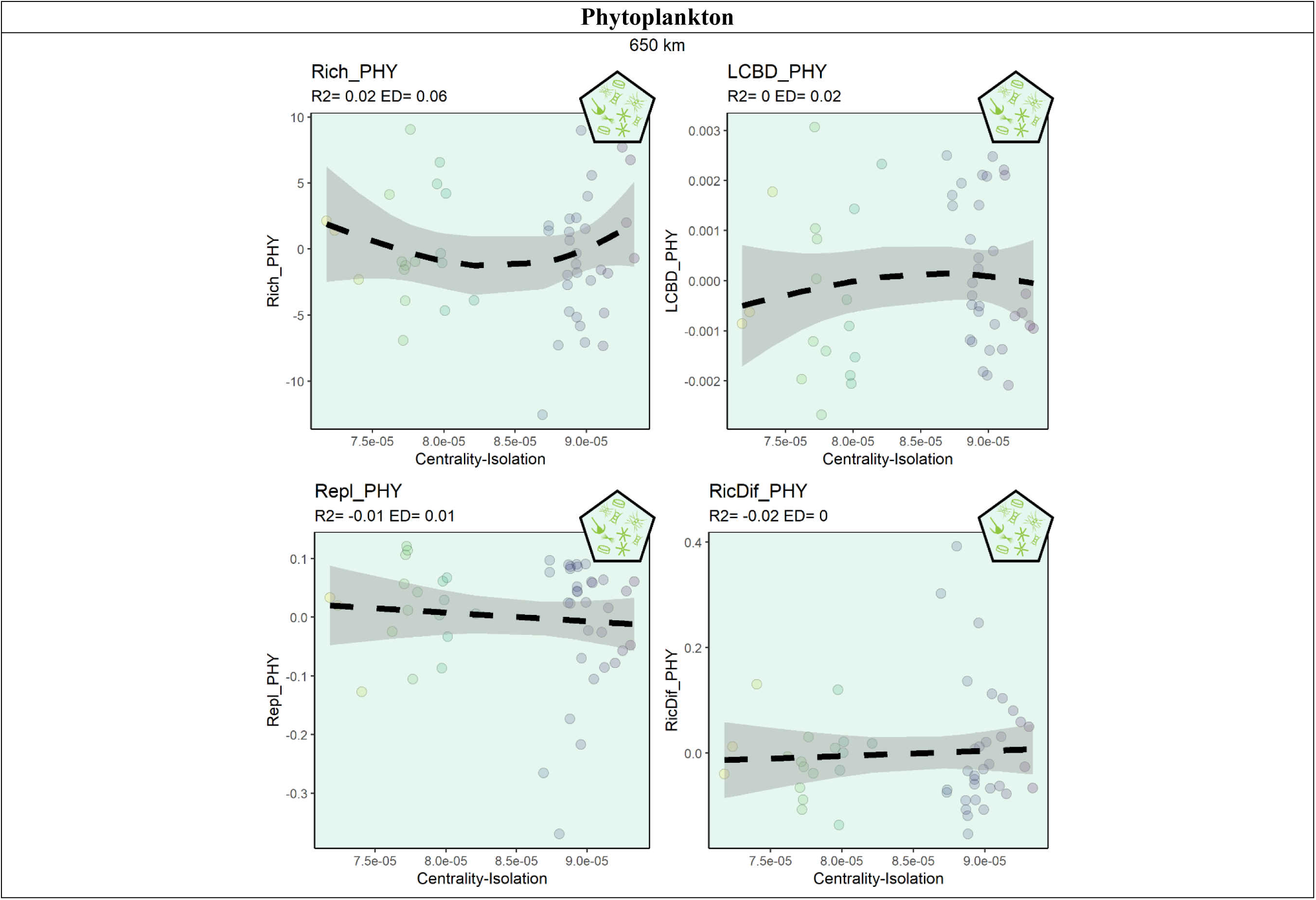
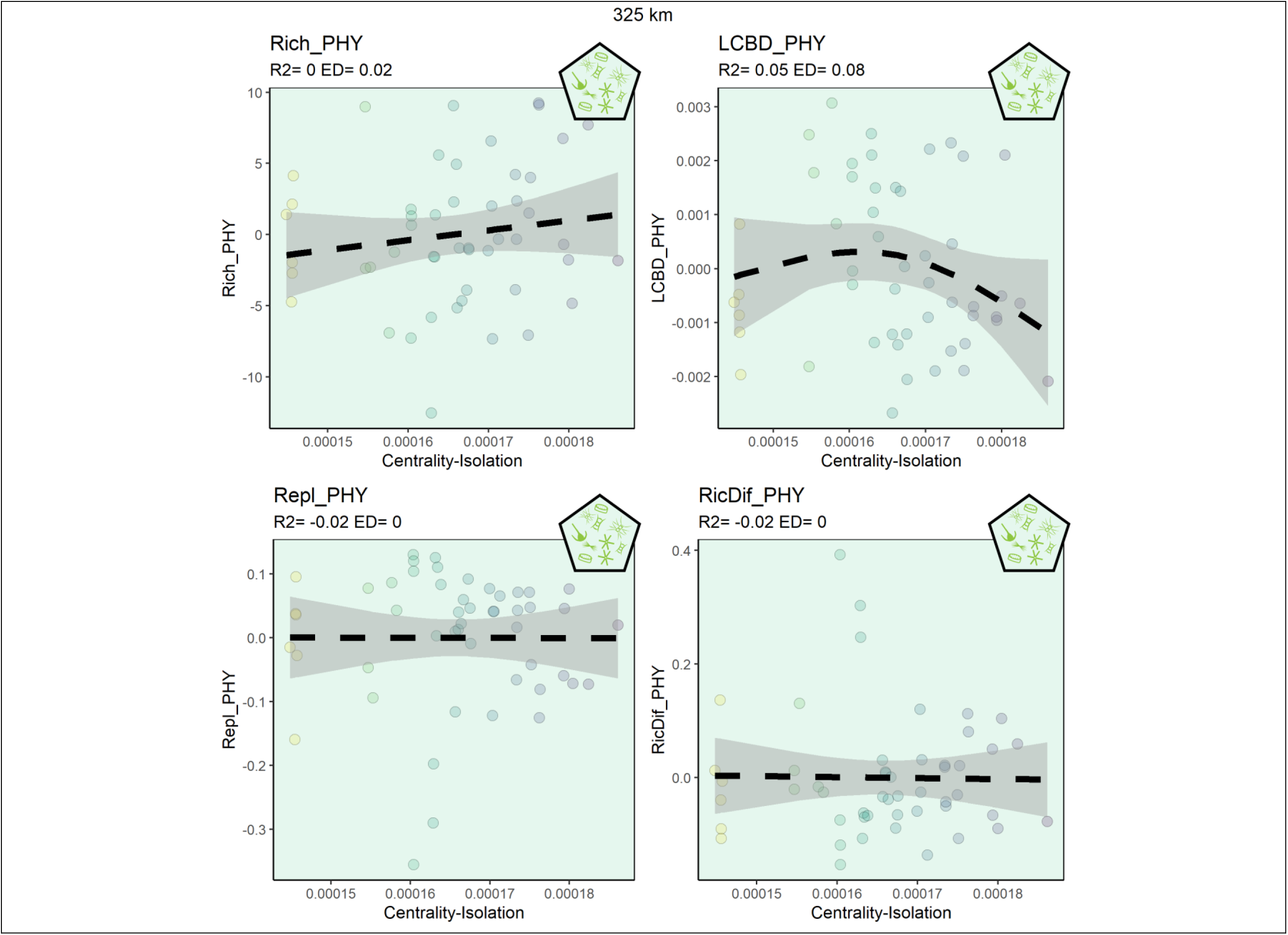
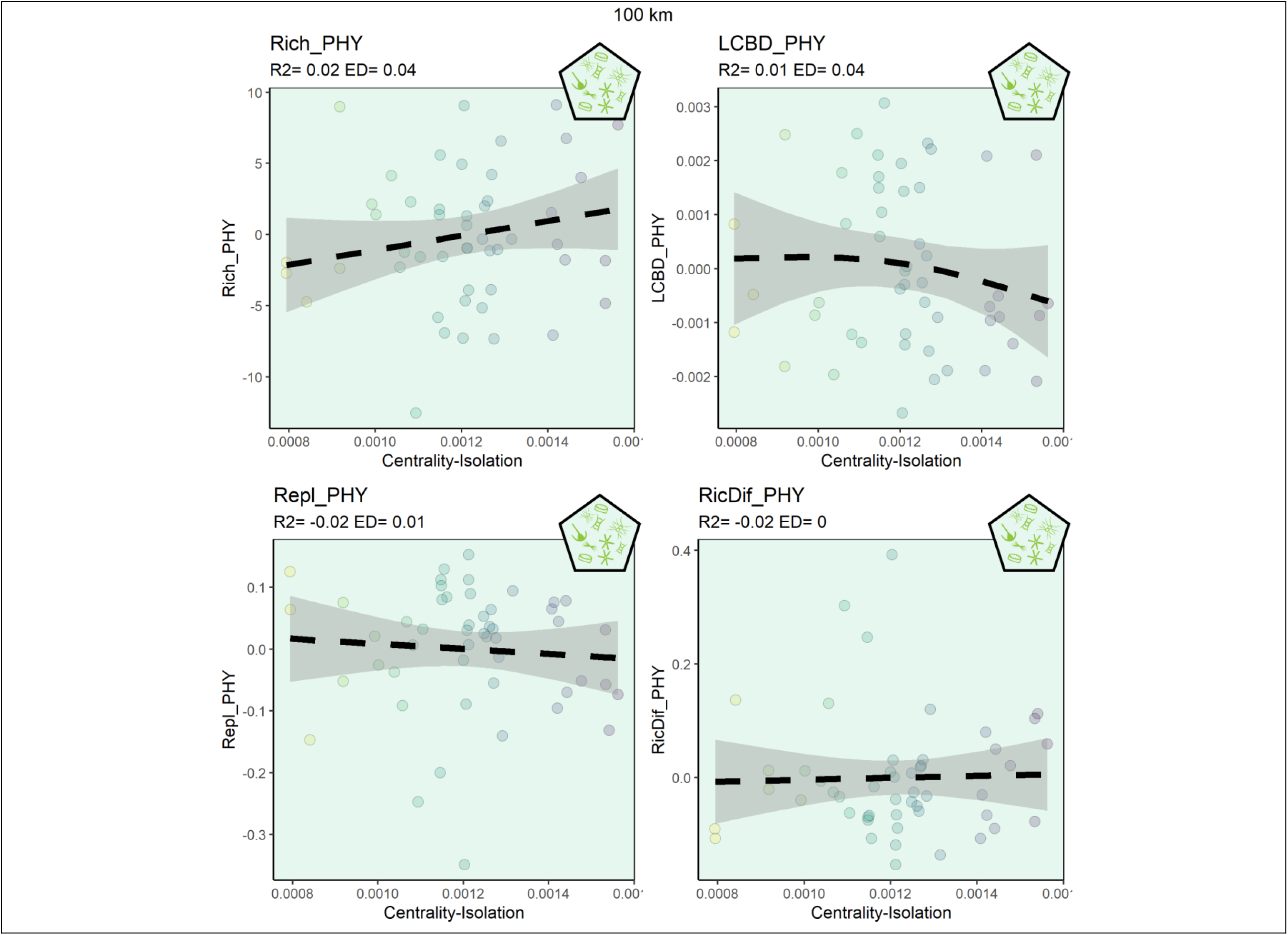
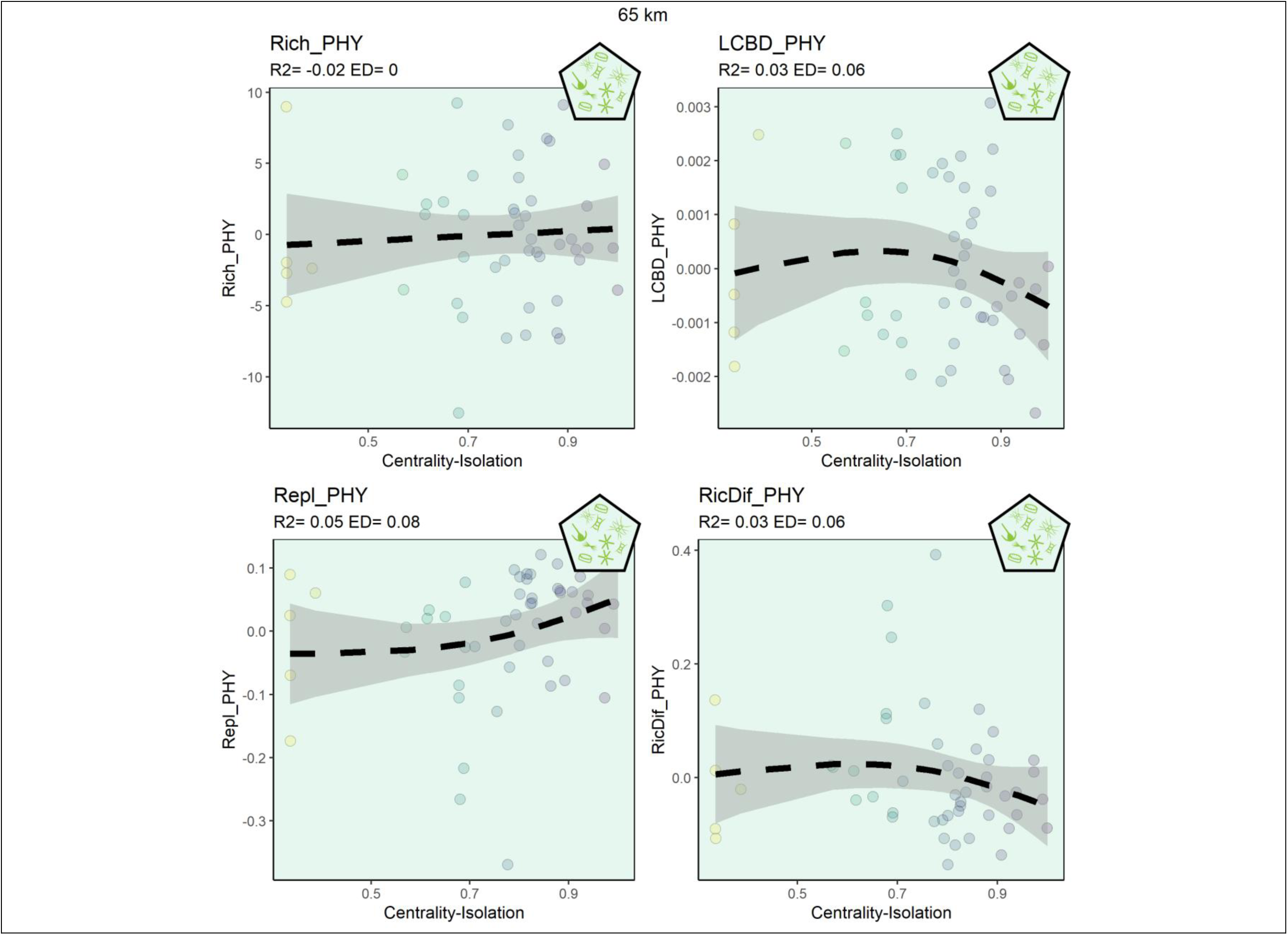
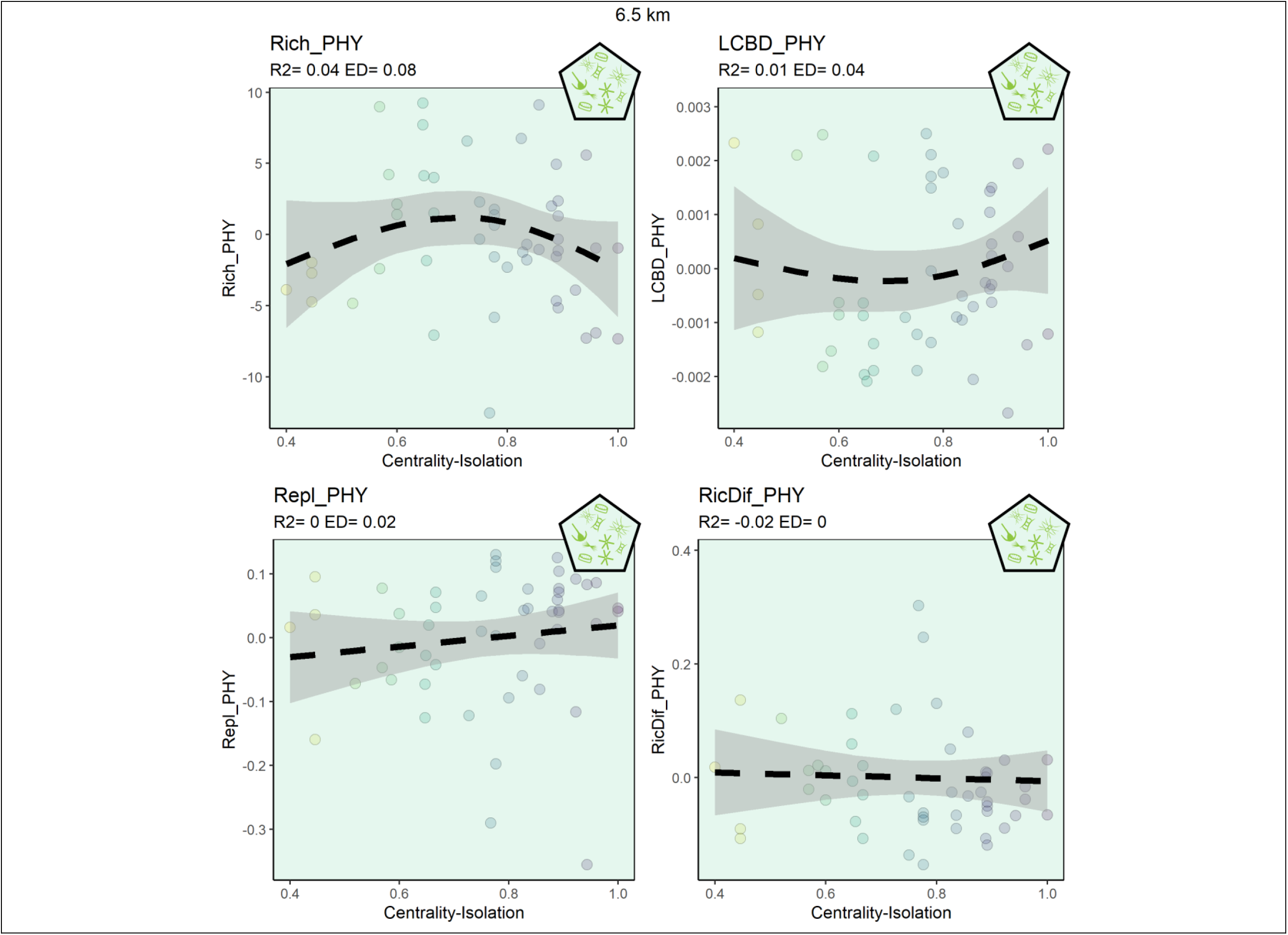
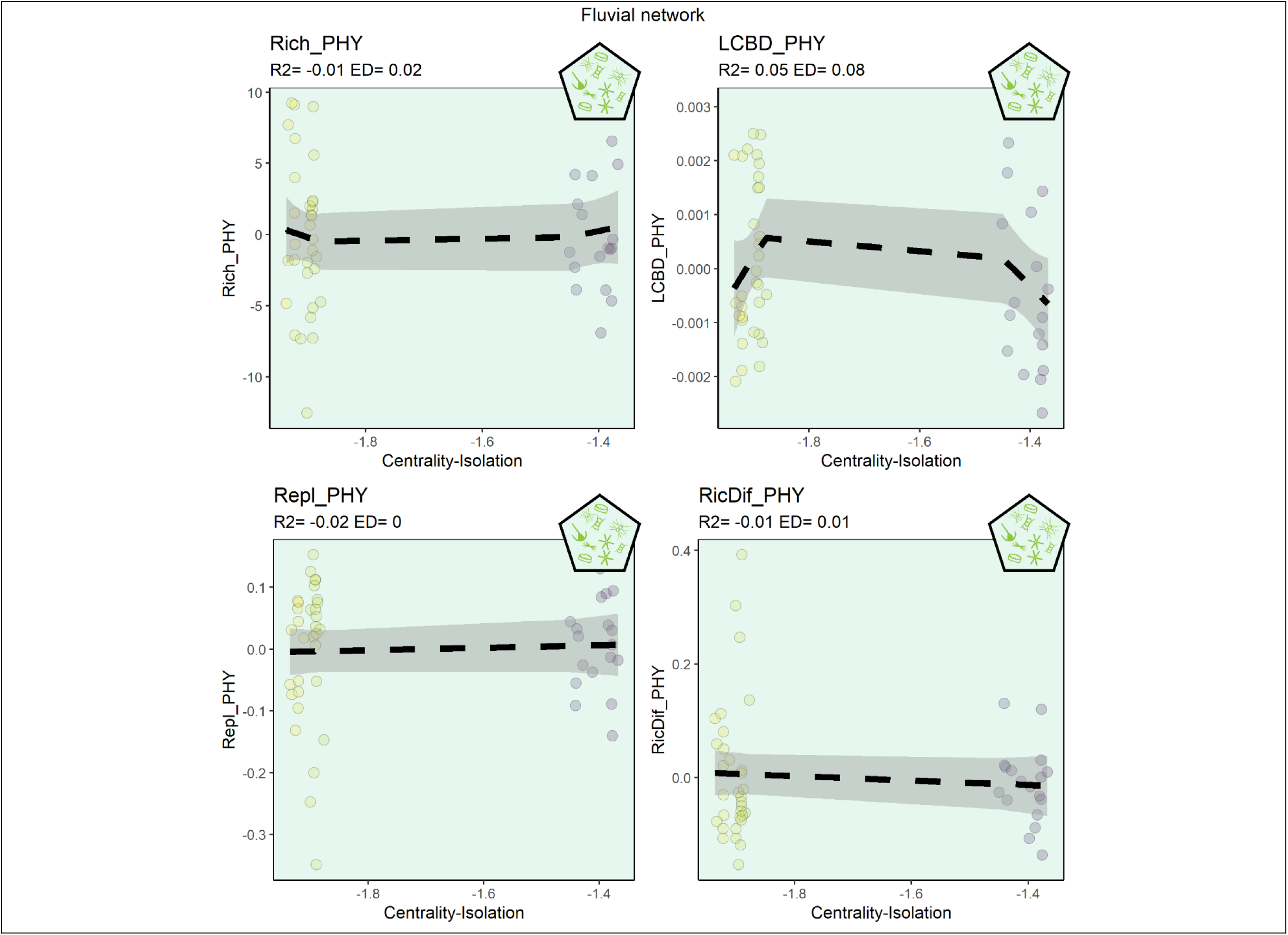
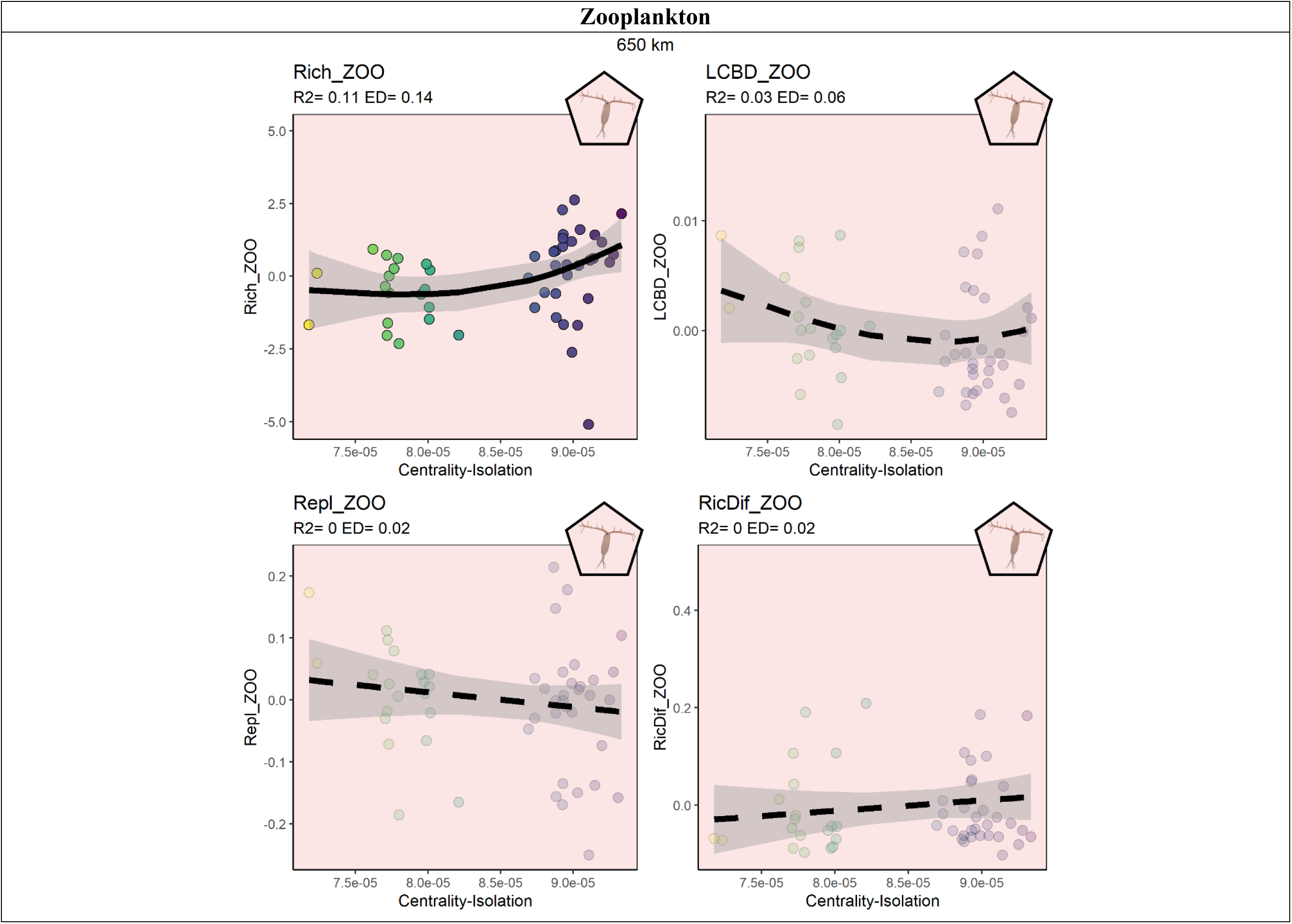
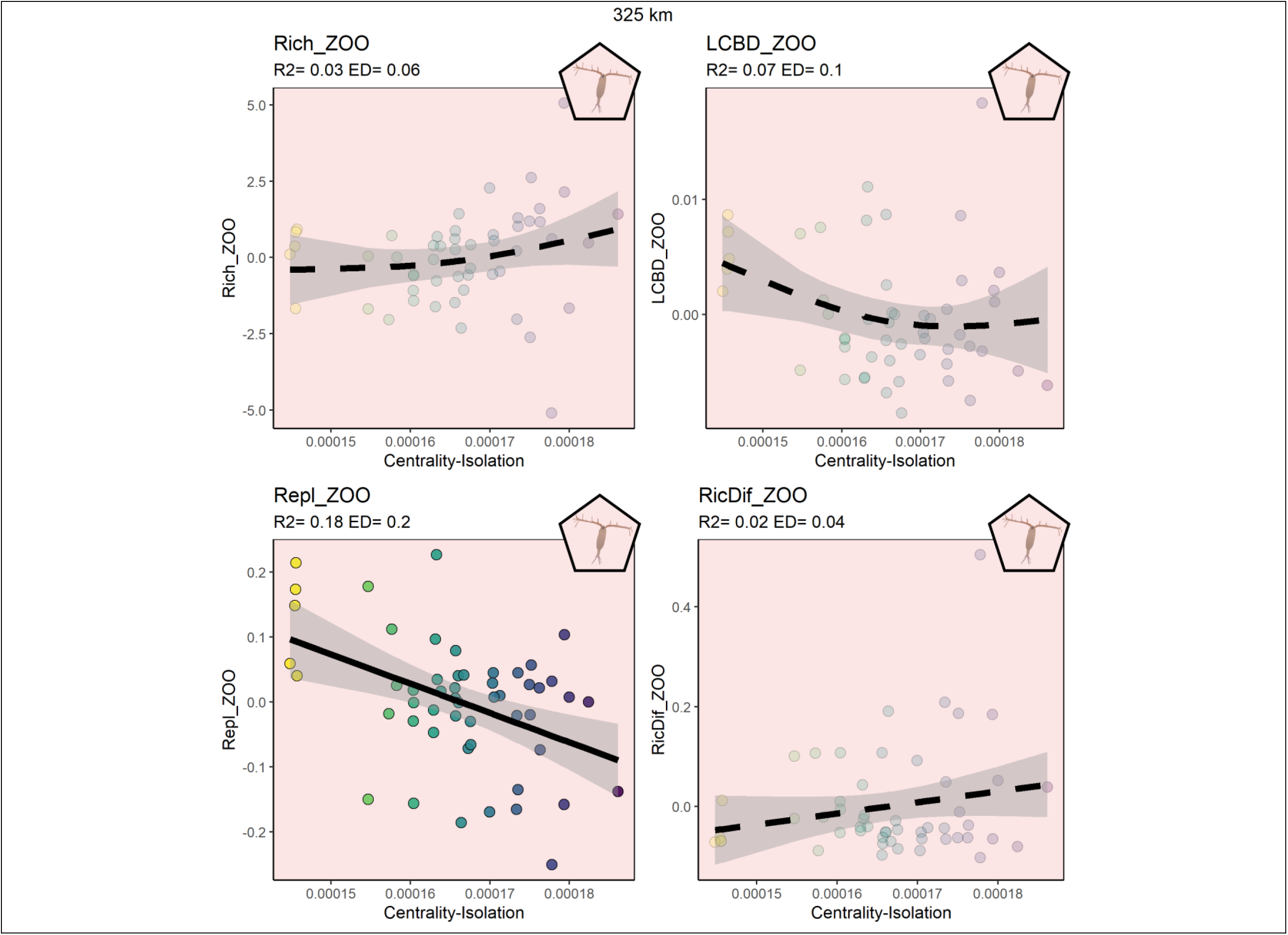
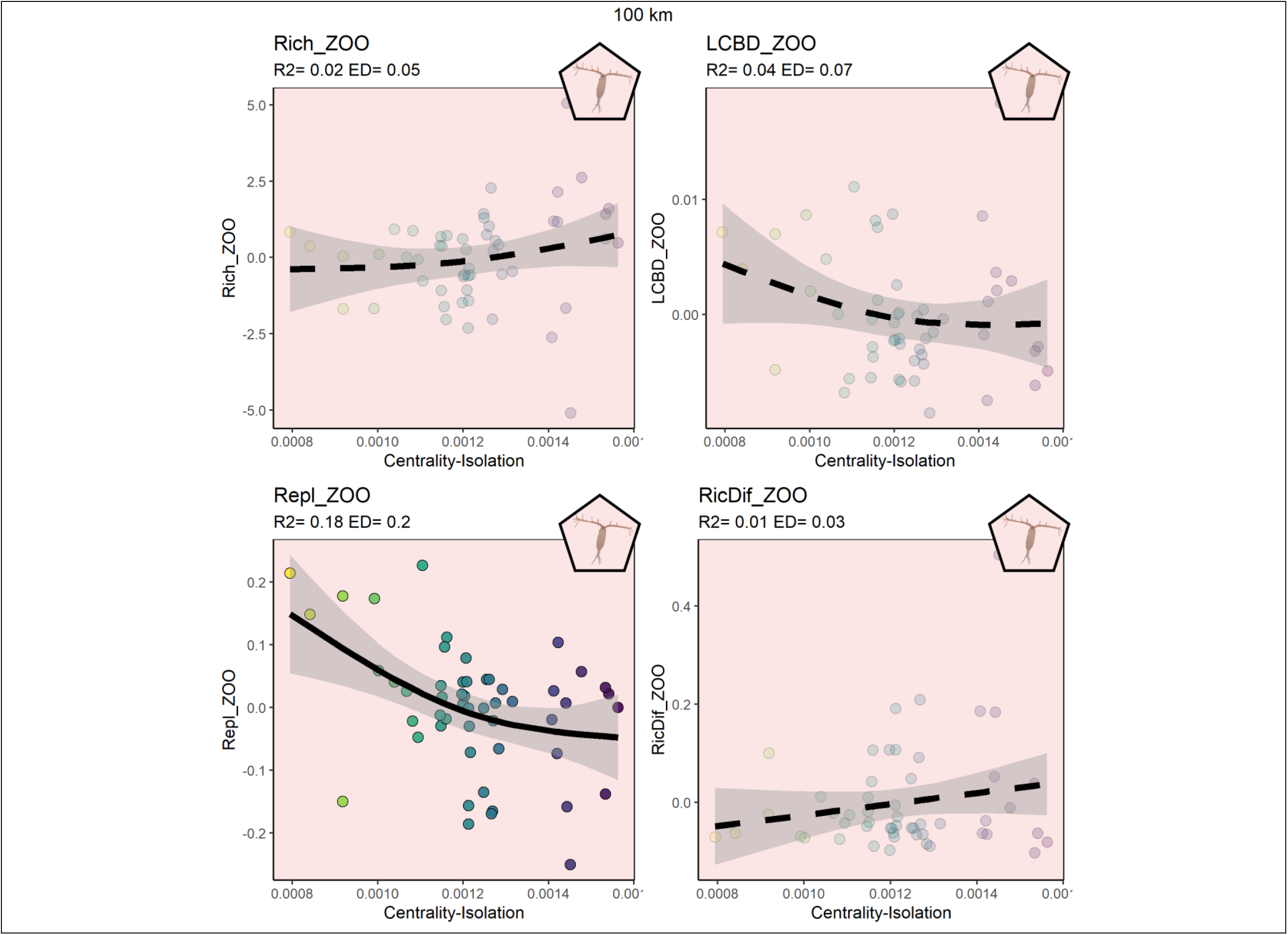
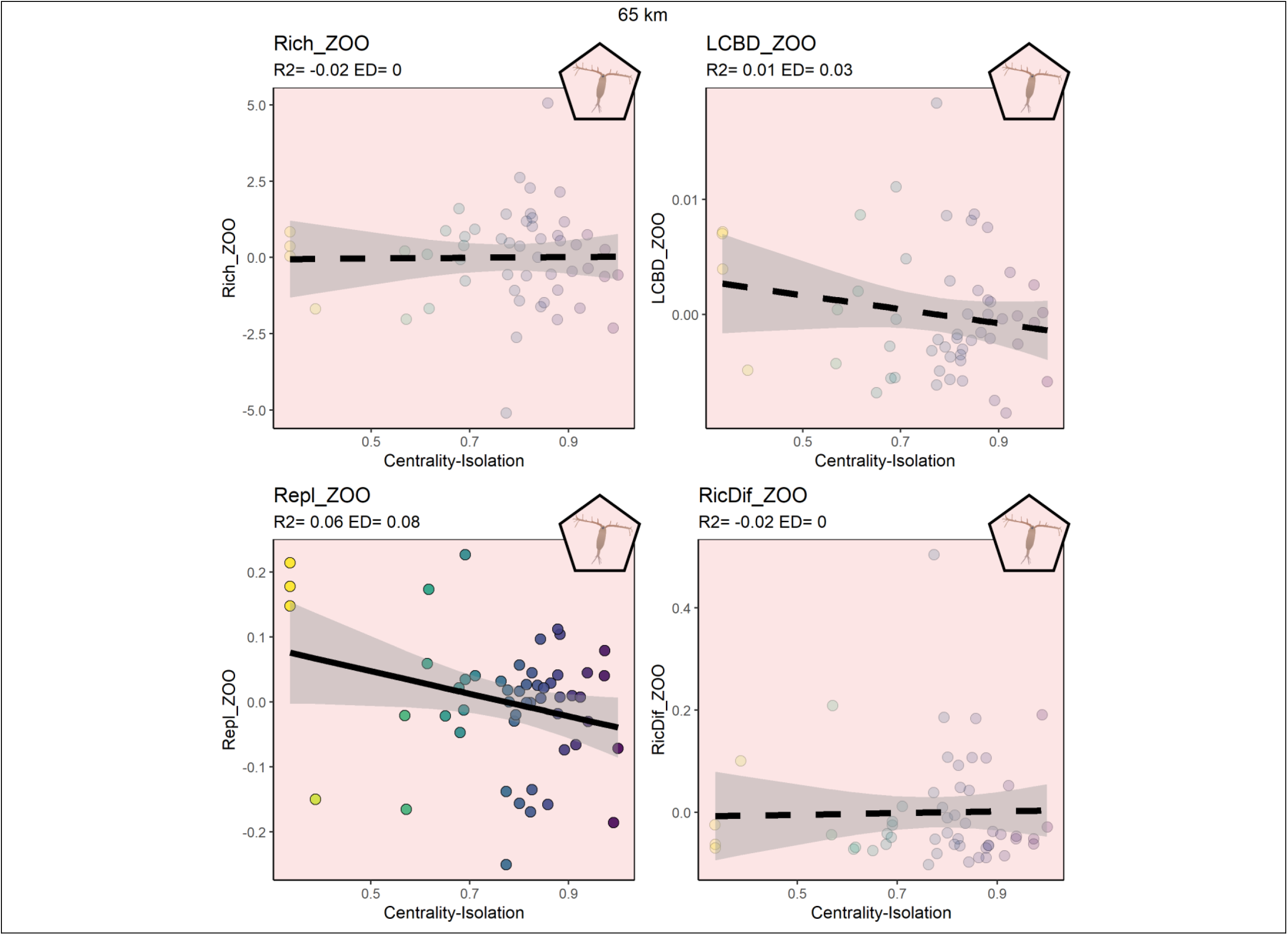
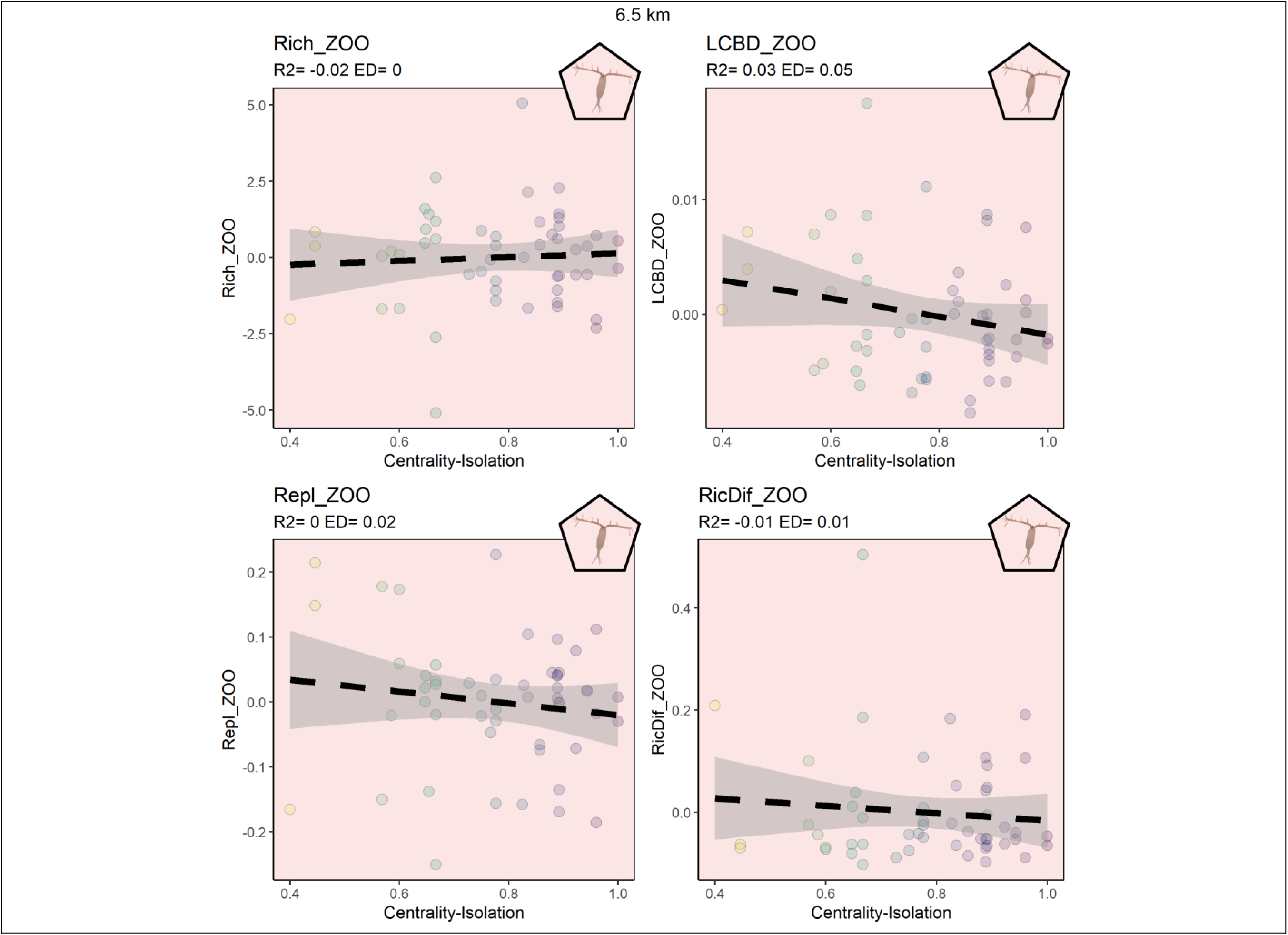
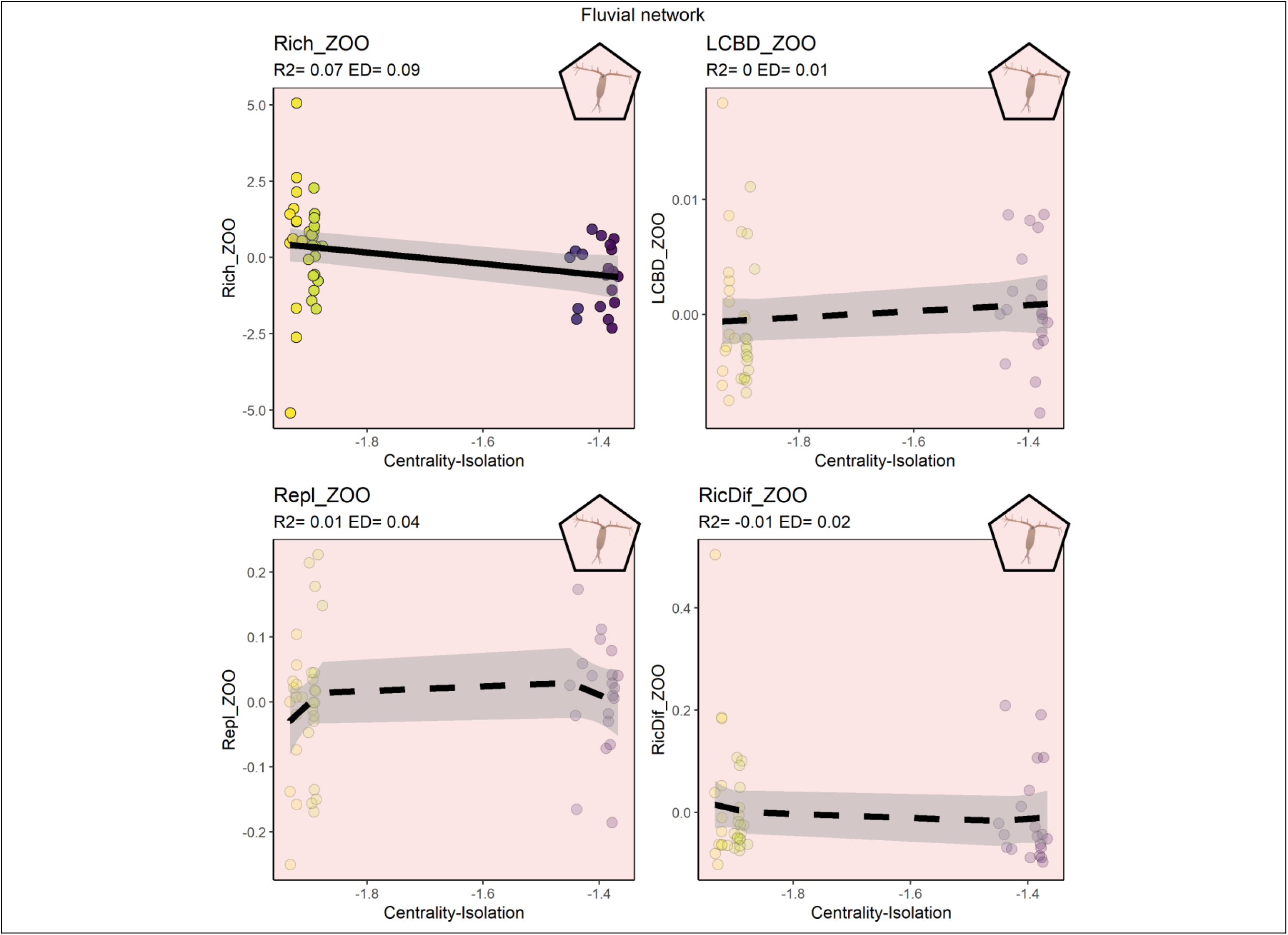
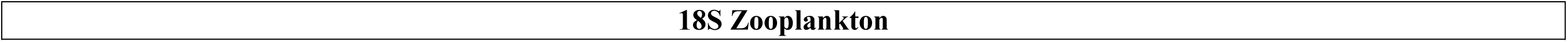
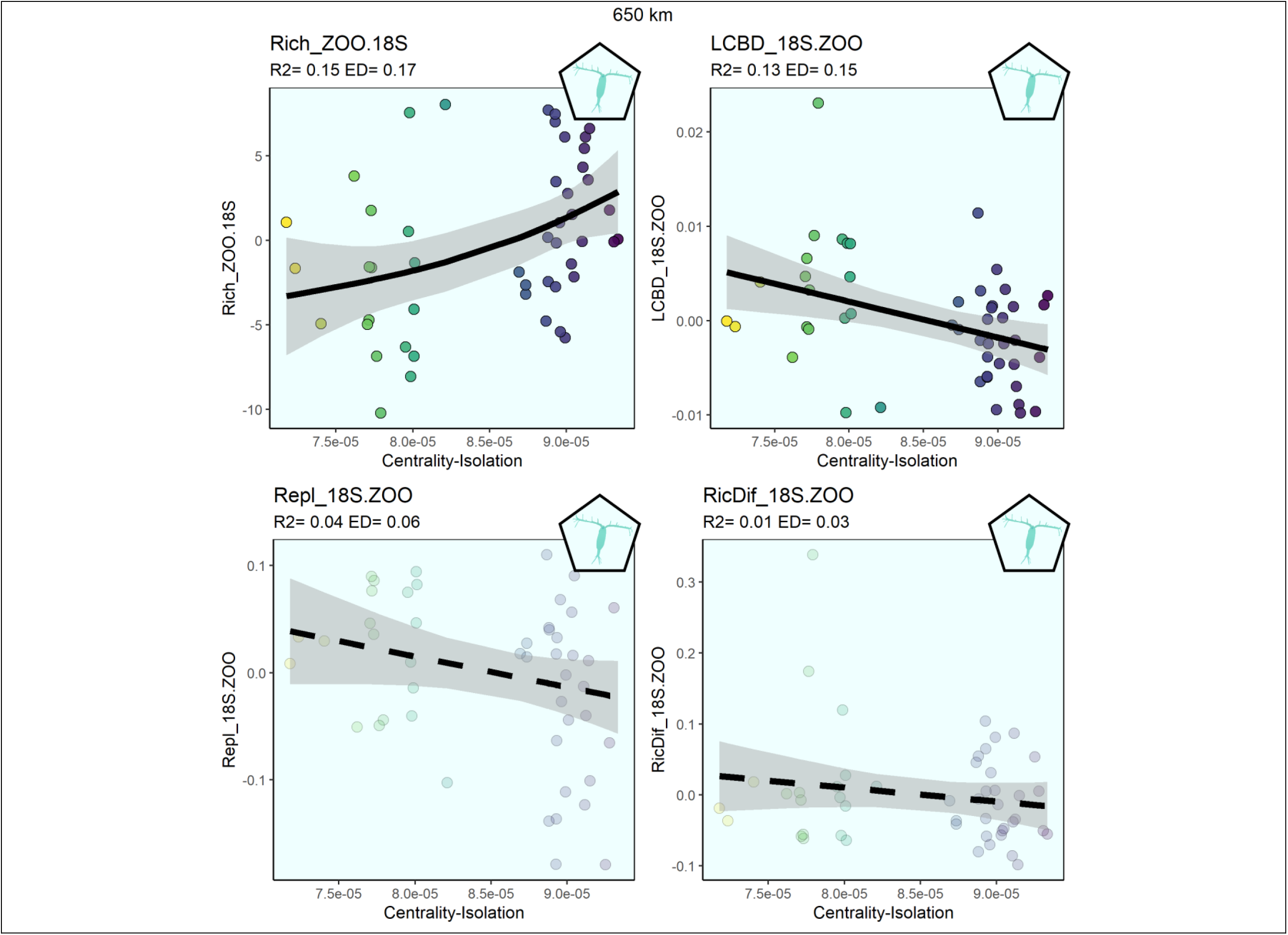
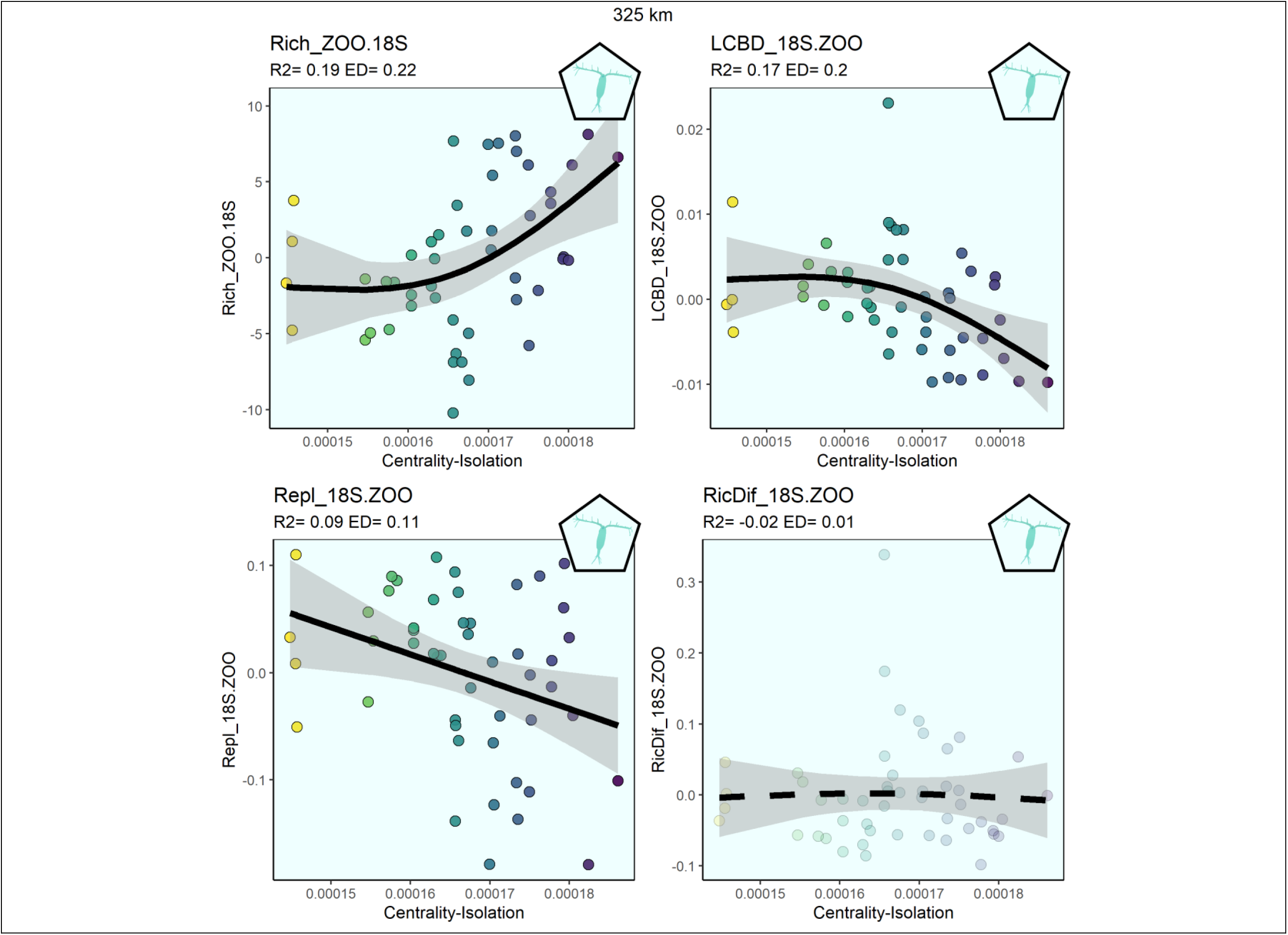
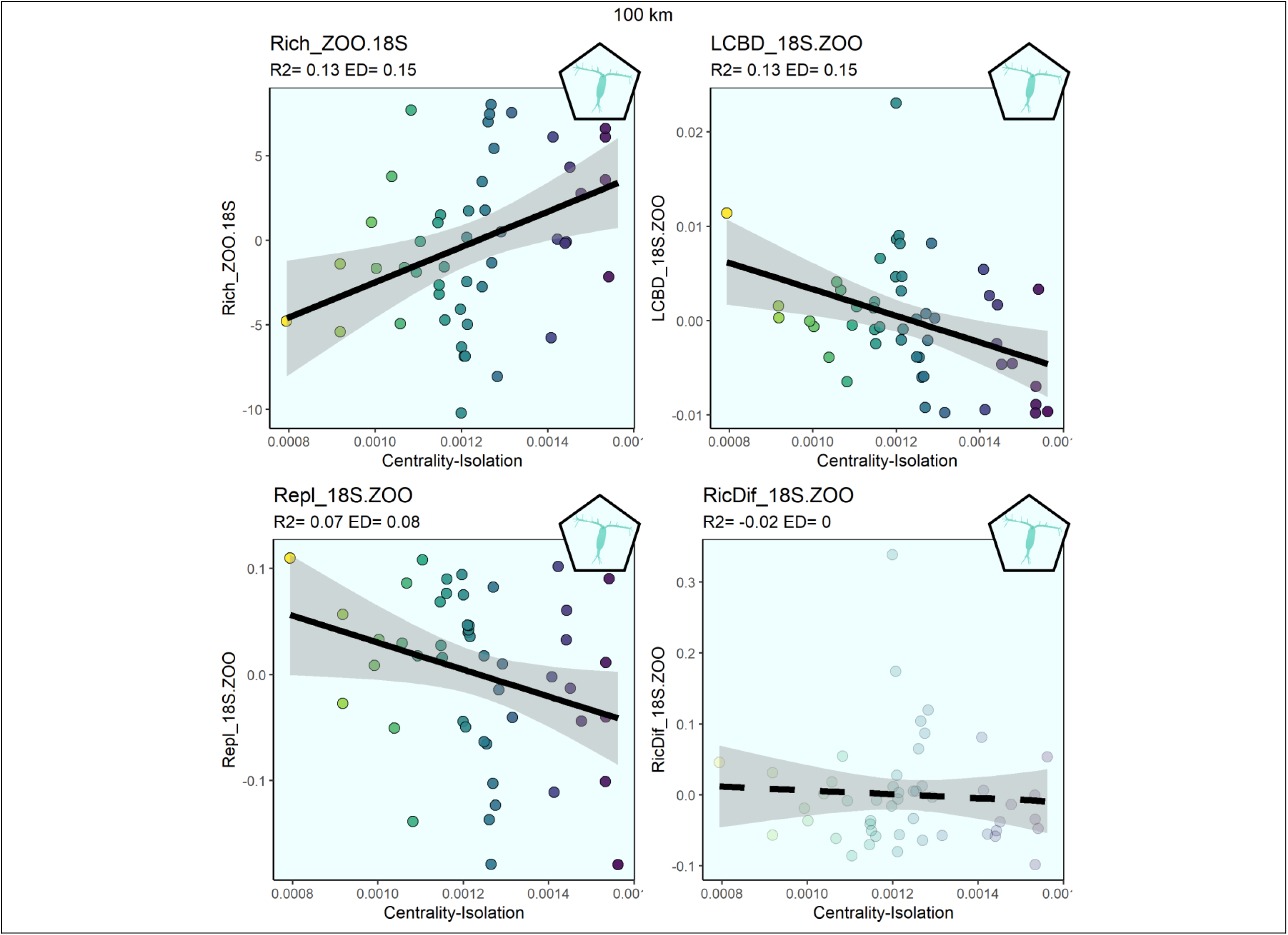
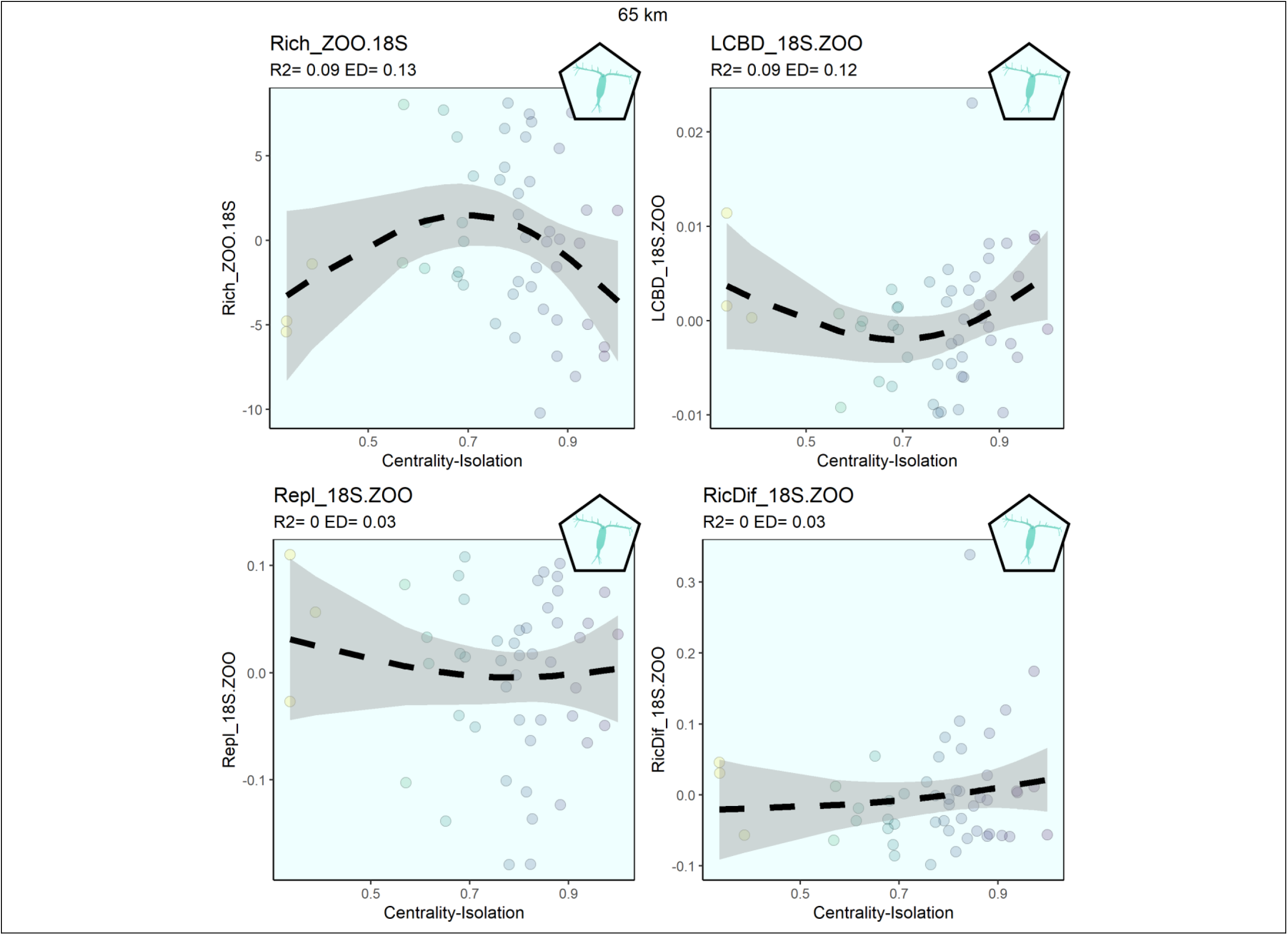
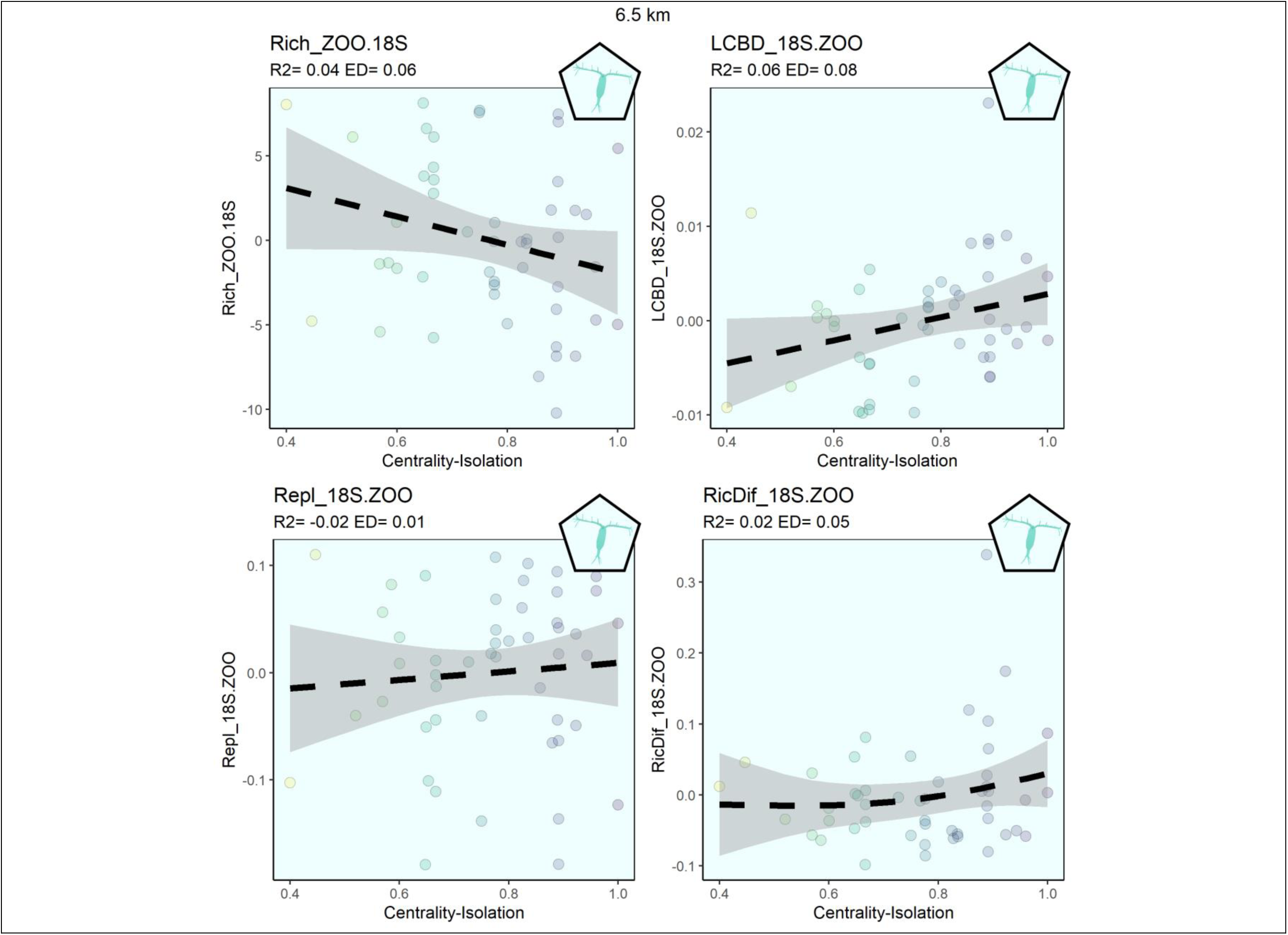
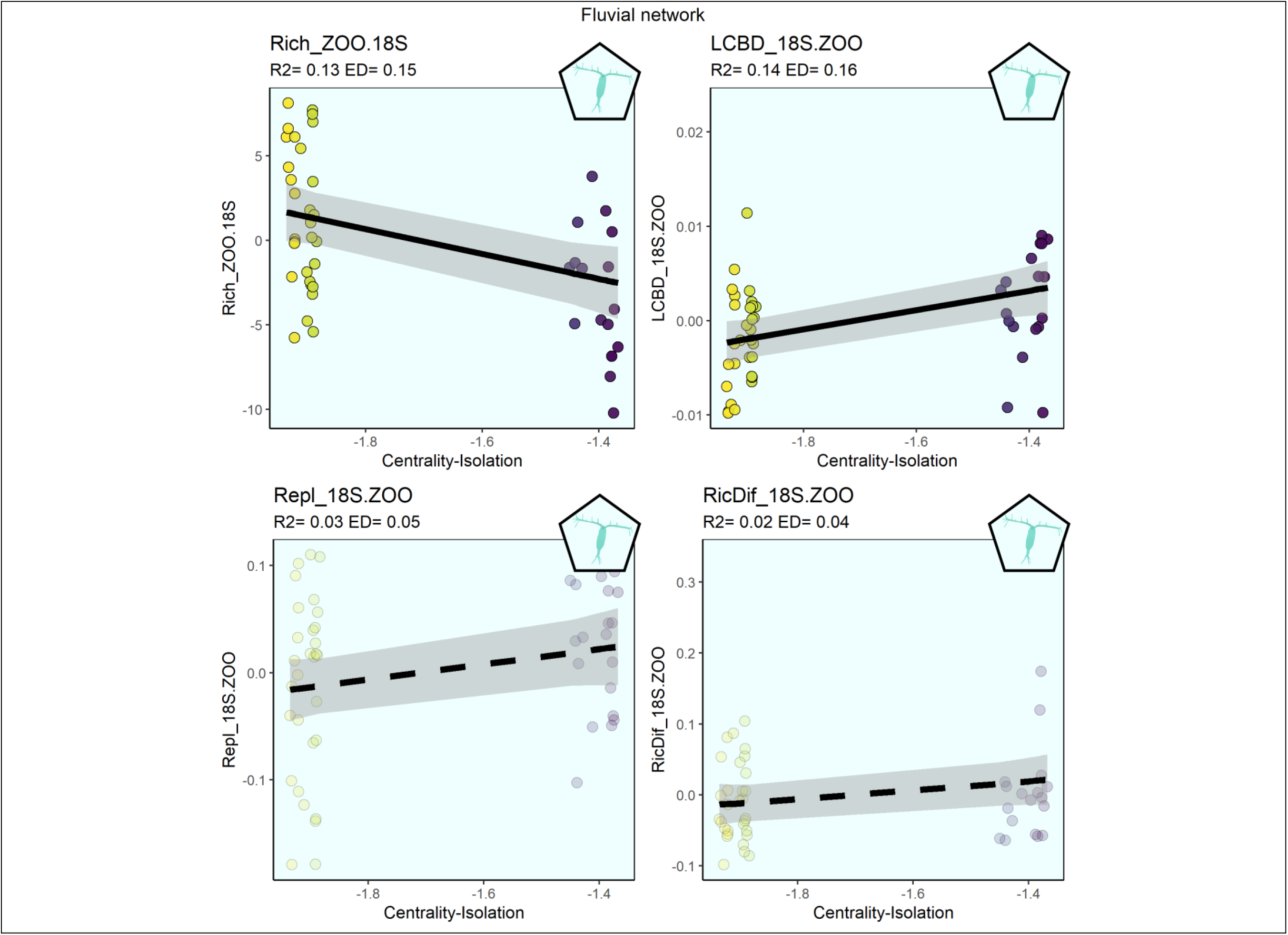
Generalized additive values model summary for each combination of group, network, and diversity metric residuals. Highlighted rows correspond to the significant relationships, also represented in Figure 4. Find below the model summary results the plots corresponding to each one of the detected relationships (both significant and non-significant ones) grouped in panels of 4 corresponding to the four diversity metrics for the same organism group (bacterioplankton in yellow, protists in blue, phytoplankton in green, zooplankton in red and 18S zooplankton in cyan) and network scales (650, 325,100,65,6.5, and Fluvial).

**Supplementary S8:**
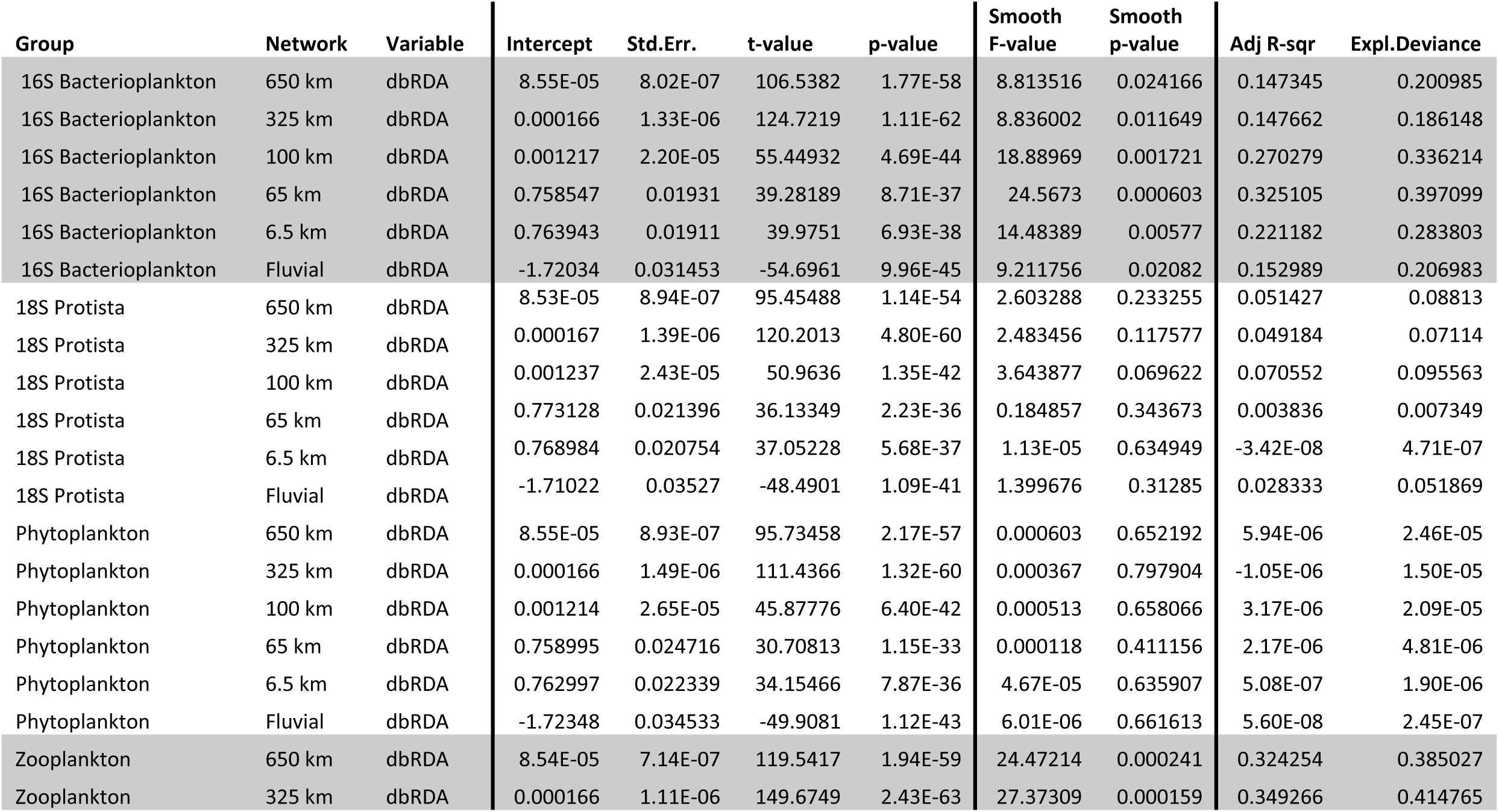
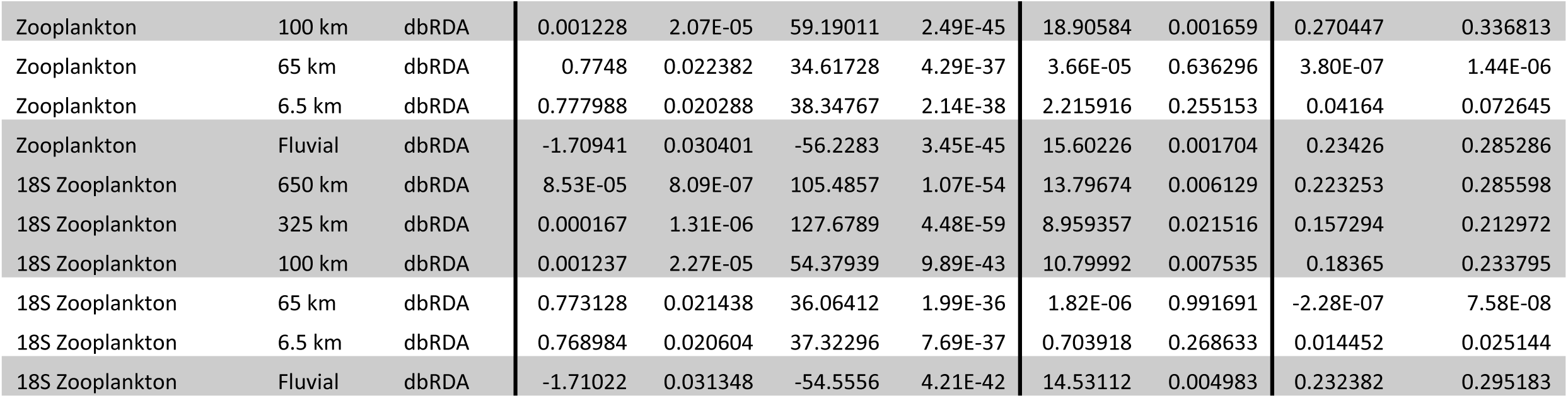
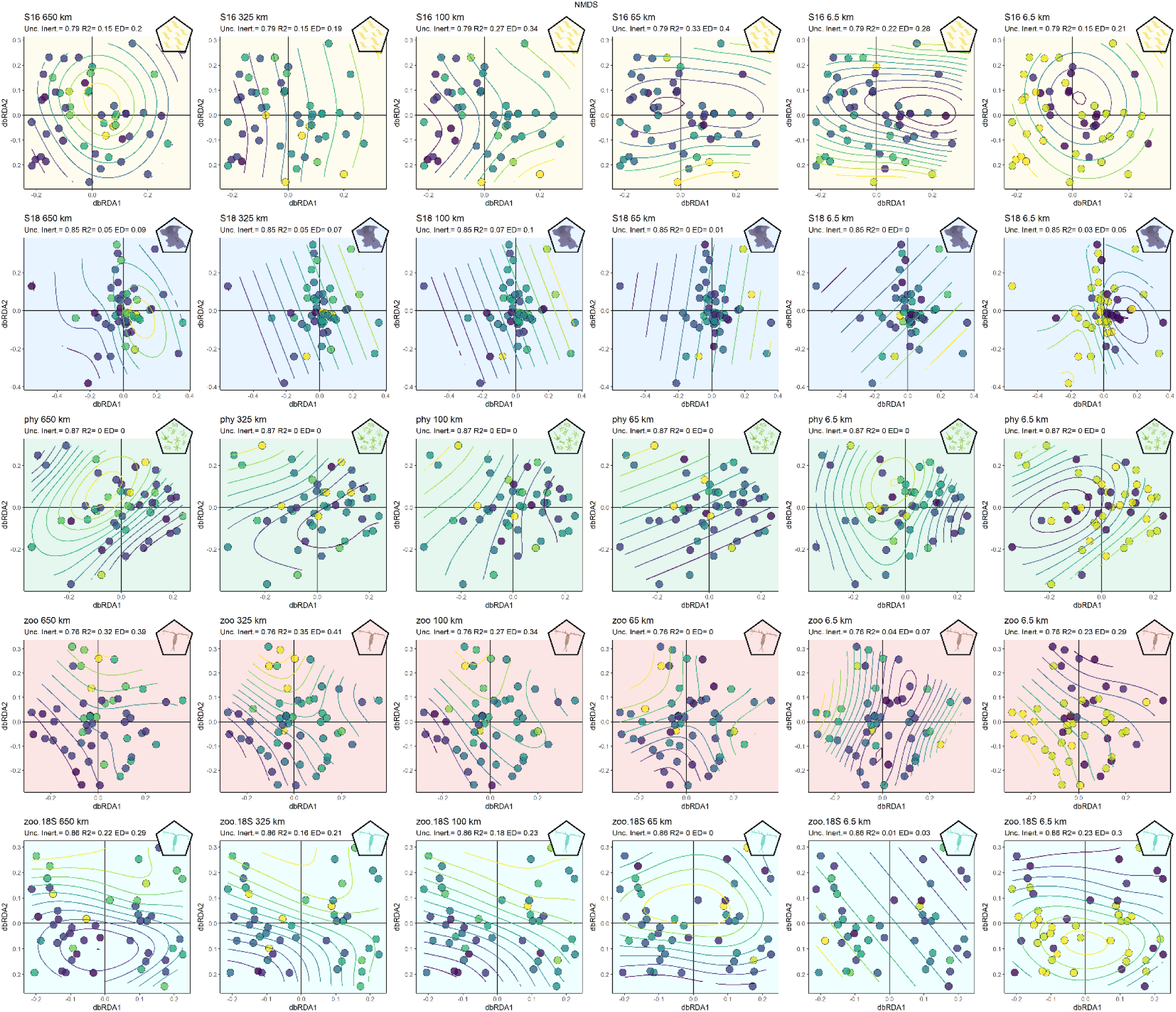
Generalized additive model summary corresponding to the smooth surfaces for each combination of group, network, and unexplained dbRDA. Highlighted rows correspond to the significant relationships, also represented in Figure 5. Find below the model summary results the plots corresponding to each one of the detected relationships (both significant and non-significant ones) for the same organism group (bacterioplankton in yellow, protists in blue, phytoplankton in green, zooplankton in red and 18S zooplankton in cyan) and network scales (650, 325,100,65,6.5, and Fluvial).

## Notes

### Competing Interest Statement

The authors have declared no competing interest.

## References

Abonyi, A., Z. Horváth, and R. Ptacnik. 2018. Functional richness outperforms taxonomic richness in predicting ecosystem functioning in natural phytoplankton communities. Freshwater Biology 63:178–186.

Ai, D., D. Gravel, C. Chu, and G. Wang. 2013. Spatial Structures of the Environment and of Dispersal Impact Species Distribution in Competitive Metacommunities. PLOS ONE 8:e68927.

Almeida-Gomes, M., F. Valente-Neto, E. O. Pacheco, C. C. Ganci, M. A. Leibold, A. S. Melo, and D. B. Provete. 2020. How Does the Landscape Affect Metacommunity Structure? A Quantitative Review for Lentic Environments. Current Landscape Ecology Reports 5:68–75.

Altermatt, F. 2013. Diversity in riverine metacommunities: A network perspective. Aquatic Ecology 47:365–377.

Altermatt, F., S. Schreiber, and M. Holyoak. 2011. Interactive effects of disturbance and dispersal directionality on species richness and composition in metacommunities. Ecology 92:859–870.

Baguette, M., S. Blanchet, D. Legrand, V. M. Stevens, and C. Turlure. 2013. Individual dispersal, landscape connectivity and ecological networks: Dispersal, connectivity and networks. Biological Reviews 88:310–326.

Baguette, M., and H. Van Dyck. 2007. Landscape connectivity and animal behavior: functional grain as a key determinant for dispersal. Landscape Ecology 22:1117–1129.

Barbour, K. M., A. Barrón-Sandoval, K. E. Walters, and J. B. H. Martiny. 2023. Towards quantifying microbial dispersal in the environment. Environmental Microbiology 25:137–142.

Başoğlu, D., M. Beklioğlu, and R. Ptacnik. 2017. AQUACOSM: Network of Leading European AQUAtic MesoCOSM Facilities, Deliverable No 4.1: Standardised protocols (SOPs) on data collection, data quality and assurances and processing. Version 1.4. Report, Forschungsverbund Berlin e.V. - Leibniz-Institut für Gewässerökologie und Binnenfischerei.

Bender, M. G., F. Leprieur, D. Mouillot, M. Kulbicki, V. Parravicini, M. R. Pie, D. R. Barneche, L. G. R. Oliveira-Santos, and S. R. Floeter. 2017. Isolation drives taxonomic and functional nestedness in tropical reef fish faunas. Ecography 40:425–435.

Bo, T., A. Doretto, M. Levrino, and S. Fenoglio. 2020. Contribution of beta diversity in shaping stream macroinvertebrate communities among hydro-ecoregions. Aquatic Ecology 6.

Boix, D., M. C. Caria, S. Gascón, M. A. Mariani, J. Sala, A. Ruhí, J. Compte, and S. Bagella. 2017. Contrasting intra-annual patterns of six biotic groups with different dispersal mode and ability in Mediterranean temporary ponds. Marine and Freshwater Research 64:1044–1060.

Borcard, D., F. Gillet, and P. Legendre. 2011. Numerical Ecology with R. Springer-Verlag New York.

Borics, G., A. Abonyi, N. Salmaso, and R. Ptacnik. 2021. Freshwater phytoplankton diversity: models, drivers and implications for ecosystem properties. Hydrobiologia 848:53–75.

Borthagaray, A. I., J. M. Barreneche, S. Abades, and M. Arim. 2014. Modularity along organism dispersal gradients challenges a prevailing view of abrupt transitions in animal landscape perception. Ecography 37:564–571.

Borthagaray, A. I., M. Berazategui, and M. Arim. 2015a. Disentangling the effects of local and regional processes on biodiversity patterns through taxon-contingent metacommunity network analysis. Oikos 124:1383–1390.

Borthagaray, A. I., D. Cunillera-Montcusí, J. Bou, J. Biggs, and M. Arim. 2023a. Pondscape or waterscape? The effect on the diversity of dispersal along different freshwater ecosystems. Hydrobiologia.

Borthagaray, A. I., D. Cunillera-Montcusí, J. Bou, I. Tornero, D. Boix, M. Anton-Pardo, E. Ortiz, T. Mehner, X. D. Quintana, S. Gascón, and M. Arim. 2023b. Heterogeneity in the isolation of patches may be essential for the action of metacommunity mechanisms. Frontiers in Ecology and Evolution 11.

Borthagaray, A. I., V. Pinelli, M. Berazategui, L. Rodríguez-Tricot, and M. Arim. 2015b. Effects of metacommunitity networks on local community structures: from theoretical predictions to empirical evaluations. Pages 75–111 in A. Belgrano, G. Woodward, and U. Jacob, editors. Aquatic Functional Biodiversity. Academic Press, Oxford.

Borthagaray, A. I., F. Teixeira-de Mello, G. Tesitore, E. Ortiz, M. Illarze, V. Pinelli, L. Urtado, P. Raftopulos, I. González-Bergonzoni, S. Abades, M. Loureiro, and M. Arim. 2020. Community isolation drives lower fish biomass and species richness, but higher functional evenness, in a river metacommunity. Freshwater Biology:1–15.

Braghin, L. S. M., B. R. S. Figueiredo, T. Meurer, T. S. Michelan, N. R. Simões, and C. C. Bonecker. 2015. Zooplankton diversity in a dammed river basin is maintained by preserved tributaries in a tropical floodplain. Aquatic Ecology 49:175–187.

Brauer, C. J., and L. B. Beheregaray. 2020. Recent and rapid anthropogenic habitat fragmentation increases extinction risk for freshwater biodiversity. Evolutionary Applications 13:2857–2869.

Burgazzi, G., A. Laini, O. Ovaskainen, M. Saccò, R. Stubbington, and P. Viaroli. 2020. Communities in high definition: spatial and environmental factors shape the microdistribution of aquatic invertebrates. Freshwater Biology.

Butts, C. T. 2023, January 24. sna: Tools for Social Network Analysis.

Cáceres, C. E., and D. a. Soluk. 2002. Blowing in the wind: A field test of overland dispersal and colonization by aquatic invertebrates. Oecologia 131:402–408.

Cañedo-Argüelles, M., C. Gutiérrez-Cánovas, R. Acosta, D. Castro-López, N. Cid, P. Fortuño, A. Munné, C. Múrria, A. R. Pimentão, R. Sarremejane, M. Soria, P. Tarrats, I. Verkaik, N. Prat, and N. Bonada. 2020. As time goes by: 20 years of changes in the aquatic macroinvertebrate metacommunity of Mediterranean river networks. Journal of Biogeography 47:1861–1874.

Caporaso, J. G., C. L. Lauber, W. A. Walters, D. Berg-Lyons, C. A. Lozupone, P. J. Turnbaugh, N. Fierer, and R. Knight. 2011. Global patterns of 16S rRNA diversity at a depth of millions of sequences per sample. Proceedings of the National Academy of Sciences 108:4516–4522.

Carrara, F., A. Rinaldo, A. Giometto, and F. Altermatt. 2014. Complex interaction of dendritic connectivity and hierarchical patch size on biodiversity in river-like landscapes. American Naturalist 183:13–25.

Chaparro, G., Z. Horváth, I. O’Farrell, R. Ptacnik, and T. Hein. 2018. Plankton metacommunities in floodplain wetlands under contrasting hydrological conditions. Freshwater Biology 63:380–391.

Chase, J. M., and T. M. Knight. 2013. Scale-dependent effect sizes of ecological drivers on biodiversity: why standardised sampling is not enough. Ecology Letters 16:17–26.

Chase, J. M., and R. S. Shulman. 2009. Wetland isolation facilitates larval mosquito density through the reduction of predators. Ecological Entomology 34:741–747.

Chaudhary, V. B., C. A. Aguilar-Trigueros, I. Mansour, and M. C. Rillig. 2022. Fungal Dispersal Across Spatial Scales. Annual Review of Ecology, Evolution, and Systematics 53:69–85.

Chisholm, C., Z. Lindo, and A. Gonzalez. 2011. Metacommunity diversity depends on connectivity and patch arrangement in heterogeneous habitat networks. Ecography 34:415–424.

Choudoir, M. J., and K. M. DeAngelis. 2022. A framework for integrating microbial dispersal modes into soil ecosystem ecology. iScience 25:103887.

Chubaty, A. M., P. Galpern, and S. C. Doctolero. 2020. The r toolbox grainscape for modelling and visualizing landscape connectivity using spatially explicit networks. Methods in Ecology and Evolution 2020:591–595.

Coughlan, N. E., T. C. Kelly, J. Davenport, and M. A. K. Jansen. 2017. Up, up and away: bird-mediated ectozoochorous dispersal between aquatic environments. Freshwater Biology 62:631–648.

Csabai, Z., and P. Boda. 2005. Effect of the wind speed on the migration activity of aquatic insects (Coleoptera, Heteroptera). Acta Biologica Debrecina Oecologica Hungarica 13:37–42.

Csárdi, G., and T. Nepusz. 2006. The igraph software package for complex network research. C.

Cunillera-Montcusí, D., D. Boix, J. Sala, J. Compte, I. Tornero, X. D. Quintana, and S. Gascón. 2020. Large- and small-regional-scale variables interact in the dispersal patterns of aquatic macroinvertebrates from temporary ponds. Aquatic Ecology 54:1041–1058.

Cunillera-Montcusí, D., A. I. Borthagaray, D. Boix, S. Gascón, J. Sala, I. Tornero, X. D. Quintana, and M. Arim. 2021. Metacommunity resilience against simulated gradients of wildfire: disturbance intensity and species dispersal ability determine landscape recover capacity. Ecography 44:1022–1034.

Custer, G. F., L. Bresciani, and F. Dini-Andreote. 2022. Ecological and Evolutionary Implications of Microbial Dispersal. Frontiers in Microbiology 13:855859.

Daffonchio, D., A. Lorz, A. Valenzuela-cuevas, A. Barozzi, and J. M. Booth. 2019. Dispersal homogenizes communities via immigration even at low rates in a simplified synthetic bacterial metacommunity. Nature Communications 10:1314.

Dale, M. 2017. Applying Graph Theory in Ecological Research. Cambridge University Press., Cambridge.

Declerck, S. a J., J. S. Coronel, P. Legendre, and L. Brendonck. 2011. Scale dependency of processes structuring metacommunities of cladocerans in temporary pools of High-Andes wetlands. Ecography 34:296–305.

Declerck, S. A. J., C. Winter, J. B. Shurin, C. A. Suttle, and B. Matthews. 2013. Effects of patch connectivity and heterogeneity on metacommunity structure of planktonic bacteria and viruses. The ISME journal 7:533–542.

Deiner, K., J.-C. Walser, E. Mächler, and F. Altermatt. 2015. Choice of capture and extraction methods affect detection of freshwater biodiversity from environmental DNA. Biological Conservation 183:53–63.

Dodson, S. 1992. Predicting crustacean zooplankton species richness. Limnology and Oceanography 37:848–856.

Doretto, A., F. Bona, E. Falasco, D. Morandini, E. Piano, and S. Fenoglio. 2020. Stay with the flow: How macroinvertebrate communities recover during the rewetting phase in Alpine streams affected by an exceptional drought. River Research and Applications 36:91–101.

Downing, J. A. 2010. Emerging global role of small lakes and ponds: Little things mean a lot. Limnetica 29:9–24.

Dray, S., P. Legendre, and P. R. Peres-Neto. 2006. Spatial modelling: a comprehensive framework for principal coordinate analysis of neighbour matrices (PCNM). Ecological Modelling 196:483–493.

Economo, E. P., and T. H. Keitt. 2008. Species diversity in neutral metacommunities: A network approach. Ecology Letters 11:52–62.

Economo, E. P., and T. H. Keitt. 2010. Network isolation and local diversity in neutral metacommunities. Oikos 119:1355–1363.

Edge, C. B., M.-J. Fortin, D. A. Jackson, D. Lawrie, L. Stanfield, and N. Shrestha. 2017. Habitat alteration and habitat fragmentation differentially affect beta diversity of stream fish communities. Landscape Ecology 32:647–662.

Epele, L. B., D. A. Dos Santos, R. Sarremejane, M. G. Grech, P. A. Macchi, L. M. Manzo, M. L. Miserendino, N. Bonada, and M. Cañedo-Argüelles. 2021. Blowin’ in the wind: Wind directionality affects wetland invertebrate metacommunities in Patagonia. Global Ecology and Biogeography:1–13.

Erős, T., and W. H. Lowe. 2019. The Landscape Ecology of Rivers: from Patch-Based to Spatial Network Analyses. Current Landscape Ecology Reports 4:103–112.

Estrada, E., and Ö. Bodin. 2008. Using Network Centrality Measures to Manage Landscape Connectivity. Ecological Applications 18:1810–1825.

Faggioni, G. P., F. L. Souza, A. C. Paranhos Filho, R. M. Gamarra, and C. P. A. Prado. 2021. Amount and spatial distribution of habitats influence occupancy and dispersal of frogs at multiple scales in agricultural landscape. Austral Ecology 46:126–138.

Florencio, M., R. Fernández-Zamudio, M. Lozano, and C. Díaz-Paniagua. 2020. Interannual variation in filling season affects zooplankton diversity in Mediterranean temporary ponds. Hydrobiologia 847:1195–1205.

Fontaneto, D., T. G. Barraclough, K. Chen, C. Ricci, and E. A. Herniou. 2008. Molecular evidence for broad-scale distributions in bdelloid rotifers: everything is not everywhere but most things are very widespread. Molecular Ecology 17:3136–3146.

Fukami, T. 2015. Historical Contingency in Community Assembly: Integrating Niches, Species Pools, and Priority Effects. Annual Review of Ecology, Evolution, and Systematics 46:1–23.

Fuller, M. R., M. W. Doyle, and D. L. Strayer. 2015. Causes and consequences of habitat fragmentation in river networks: River fragmentation. Annals of the New York Academy of Sciences 1355:31–51.

Füreder, L., R. Ettinger, A. Boggero, B. Thaler, and H. Thies. 2006. Macroinvertebrate Diversity in Alpine Lakes: Effects of Altitude and Catchment Properties. Hydrobiologia 562:123–144.

Gilarranz, L. J. 2020. Generic Emergence of Modularity in Spatial Networks. Scientific Reports 10:1–8.

Gilbert, B., and J. R. Bennett. 2010. Partitioning variation in ecological communities: Do the numbers add up? Journal of Applied Ecology 47:1071–1082.

Gounand, I., E. Harvey, C. J. Little, and F. Altermatt. 2018. Meta -Ecosystems 2.0: Rooting the Theory into the Field. Trends in Ecology & Evolution 33:36–46.

Grainger, T. N., and B. Gilbert. 2016. Dispersal and diversity in experimental metacommunities: linking theory and practice. Oikos 125:1213–1223.

Green, A. J., J. Figuerola, and M. I. Sánchez. 2002. Implications of waterbird ecology for the dispersal of aquatic organisms. Acta Oecologica 23:177–189.

Hanski, I. 1999. Habitat Connectivity, Habitat Continuity, and Metapopulations in Dynamic Landscapes. Oikos 87:209.

Heino, J., J. Alahuhta, L. M. Bini, Y. Cai, A. S. Heiskanen, S. Hellsten, P. Kortelainen, N. Kotamäki, K. T. Tolonen, P. Vihervaara, A. Vilmi, and D. G. Angeler. 2021. Lakes in the era of global change: moving beyond single-lake thinking in maintaining biodiversity and ecosystem services. Biological Reviews 96:89–106.

Heino, J., A. S. Melo, T. Siqueira, J. Soininen, S. Valanko, and L. M. Bini. 2015. Metacommunity organisation, spatial extent and dispersal in aquatic systems: patterns, processes and prospects. Freshwater Biology 60:845–869.

Henriques-Silva, R., M. Logez, N. Reynaud, P. A. Tedesco, S. Brosse, S. R. Januchowski-Hartley, T. Oberdorff, and C. Argillier. 2019. A comprehensive examination of the network position hypothesis across multiple river metacommunities. Ecography 42:284–294.

Herrera-Pérez, J., J. L. Parra, D. Restrepo-Santamaría, and L. F. Jiménez-Segura. 2019. The Influence of Abiotic Environment and Connectivity on the Distribution of Diversity in an Andean Fish Fluvial Network. Frontiers in Environmental Science 7.

Hill, M. J., H. M. Greaves, C. D. Sayer, C. Hassall, M. Milin, V. S. Milner, L. Marazzi, R. Hall, L. R. Harper, I. Thornhill, R. Walton, J. Biggs, N. Ewald, A. Law, N. Willby, J. C. White, R. A. Briers, K. L. Mathers, M. J. Jeffries, and P. J. Wood. 2021. Pond ecology and conservation: research priorities and knowledge gaps. Ecosphere 12:e03853.

Hill, M. J., J. Heino, I. Thornhill, D. B. Ryves, and P. J. Wood. 2017. Effects of dispersal mode on the environmental and spatial correlates of nestedness and species turnover in pond communities. Oikos 126:1575–1585.

Hillebrand, H., C.-D. Dürselen, D. Kirschtel, U. Pollingher, and T. Zohary. 1999. Biovolume calculation for pelagic and benthic microalgae. Journal of Phycology 35:403–424.

Horváth, Z., R. Ptacnik, C. F. Vad, and J. M. Chase. 2019. Habitat loss over six decades accelerates regional and local biodiversity loss via changing landscape connectance. Ecology Letters 22:1019–1027.

Horváth, Z., C. F. Vad, C. Preiler, J. Birtel, B. Matthews, R. Ptáčníková, and R. Ptáčník. 2017. Zooplankton communities and Bythotrephes longimanus in lakes of the montane region of the northern Alps. Inland Waters 7:3–13.

Horváth, Z., C. F. Vad, and R. Ptacnik. 2016. Wind dispersal results in a gradient of dispersal limitation and environmental match among discrete aquatic habitats. Ecography 39:726–732.

Hugerth, L. W., E. E. L. Muller, Y. O. O. Hu, L. A. M. Lebrun, H. Roume, D. Lundin, P. Wilmes, and A. F. Andersson. 2014. Systematic Design of 18S rRNA Gene Primers for Determining Eukaryotic Diversity in Microbial Consortia. PLOS ONE 9:e95567.

Incagnone, G., F. Marrone, R. Barone, L. Robba, and L. Naselli-Flores. 2015. How do freshwater organisms cross the ‘‘dry ocean’’? A review on passive dispersal and colonization processes with a special focus on temporary ponds. Hydrobiologia 750:103–123.

Ishiyama, N., T. Akasaka, and F. Nakamura. 2014. Mobility-dependent response of aquatic animal species richness to a wetland network in an agricultural landscape. Aquatic Sciences 76:437– 449.

Ishwaran, H., and U. B. Kogalur. 2023, May 23. randomForestSRC: Fast Unified Random Forests for Survival, Regression, and Classification (RF-SRC).

Jacquemin, C., C. Bertrand, E. Franquet, S. Mounier, B. Misson, B. Oursel, and L. Cavalli. 2019. Effects of catchment area and nutrient deposition regime on phytoplankton functionality in alpine lakes. Science of The Total Environment 674:114–127.

Jamoneau, A., S. I. Passy, J. Soininen, T. Leboucher, and J. Tison-Rosebery. 2018. Beta diversity of diatom species and ecological guilds: Response to environmental and spatial mechanisms along the stream watercourse. Freshwater Biology 63:62–73.

Kappes, H., and P. Haase. 2012. Slow, but steady: Dispersal of freshwater molluscs. Aquatic Sciences 74:1–14.

Keitt, T. H. 1997. Stability and complexity on a lattice: Coexistence of species in an individual-based food web model. Ecological Modelling 102:243–258.

Klaus, M., J. Karlsson, and D. Seekell. 2021. Tree line advance reduces mixing and oxygen concentrations in arctic–alpine lakes through wind sheltering and organic carbon supply. Global Change Biology 27:4238–4253.

Lanzén, A., S. L. Jørgensen, D. H. Huson, M. Gorfer, S. H. Grindhaug, I. Jonassen, L. Øvreås, and T. Urich. 2012. CREST – Classification Resources for Environmental Sequence Tags. PLOS ONE 7:e49334.

Larsen, L. G., J. Choi, M. K. Nungesser, and J. W. Harvey. 2012. Directional connectivity in hydrology and ecology. Ecological Applications 22:2204–2220.

Legendre, P. 2014. Interpreting the replacement and richness difference components of beta diversity. Global Ecology and Biogeography 23:1324–1334.

Legendre, P., and M. De Cáceres. 2013. Beta diversity as the variance of community data: dissimilarity coefficients and partitioning. Ecology Letters 16:951–963.

Lehner, B., and P. Do. 2004. Development and validation of a global database of lakes, reservoirs and wetlands. Journal of Hydrology 296:1–22.

Lehner, B., and G. Grill. 2013. Global river hydrography and network routing: Baseline data and new approaches to study the world’s large river systems. Hydrological Processes 27:2171–2186.

Lehner, B., K. Verdin, and A. Jarvis. 2008. New global hydrography derived from spaceborne elevation data. Eos, Transactions, American Geophysical Union 89:93–94.

Leibold, M. A., and J. M. Chase. 2018. Metacommunity Ecology. Princeton University Press.

Leibold, M. A., M. Holyoak, N. Mouquet, P. Amarasekare, J. M. M. Chase, M. F. F. Hoopes, R. D. D. Holt, J. B. B. Shurin, R. Law, D. Tilman, M. Loreau, and A. Gonzalez. 2004. The metacommunity concept: a framework for multi-scale community ecology. Ecology Letters 7:601–613.

Leibold, M. A., and J. Norberg. 2004. Biodiversity in metacommunities: Plankton as complex adaptive systems? Limnology and Oceanography 49:1278–1289.

Li, C., W. Feng, H. Chen, X. Li, F. Song, W. Guo, J. P. Giesy, and F. Sun. 2019. Temporal variation in zooplankton and phytoplankton community species composition and the affecting factors in Lake Taihu—a large freshwater lake in China. Environmental Pollution 245:1050–1057.

Li, Z., Y. Gao, S. Wang, Y. Lu, K. Sun, J. Jia, and Y. Wang. 2021. Phytoplankton community response to nutrients along lake salinity and altitude gradients on the Qinghai-Tibet Plateau. Ecological Indicators 128:107848.

Loewen, C. J. G., F. R. Wyatt, C. A. Mortimer, R. D. Vinebrooke, and R. W. Zurawell. 2020. Multiscale drivers of phytoplankton communities in north-temperate lakes. Ecological Applications 30:e02102.

Lovas-Kiss, Á., M. I. Sánchez, D. M. Wilkinson, N. E. Coughlan, J. A. Alves, and A. J. Green. 2019. Shorebirds as important vectors for plant dispersal in Europe. Ecography 42:956–967.

Lovas-Kiss, Á., O. Vincze, E. Kleyheeg, G. Sramkó, L. Laczkó, R. Fekete, A. Molnár V., and A. J. Green. 2020. Seed mass, hardness, and phylogeny explain the potential for endozoochory by granivorous waterbirds. Ecology and Evolution 10:1413–1424.

Lu, B., H. Sun, P. Harris, M. Xu, and M. Charlton. 2018. Shp2graph : Tools to Convert a Spatial Network into an Igraph Graph in R. International Journal of Geo-Information 7:1–19.

MacArthur, R. H., and E. O. Wilson. 1963. An Equilibrium Theory of Insular Zoogeography. Evolution 17:373–387.

Martins, K. P., M. G. da S. Bandeira, C. Palma-Silva, and E. F. Albertoni. 2019. Microcrustacean metacommunities in urban temporary ponds. Aquatic Sciences 81.

Michels, E., K. Cottenie, L. Neys, and L. De Meester. 2001. Zooplankton on the move: first results on the quantification of dispersal of zooplankton in a set of interconnected ponds. Hydrobiologia 442:117–126.

Monaghan, M. T., C. T. Robinson, P. Spaak, and J. V. Ward. 2005. Macroinvertebrate diversity in fragmented Alpine streams: Implications for freshwater conservation. Aquatic Sciences 67:454– 464.

Morán-Ordóñez, A., A. Pavlova, A. M. Pinder, L. Sim, P. Sunnucks, R. M. Thompson, and J. Davis. 2015. Aquatic communities in arid landscapes: Local conditions, dispersal traits and landscape configuration determine local biodiversity. Diversity and Distributions 21:1230–1241.

Moser, K. A., J. S. Baron, J. Brahney, I. A. Oleksy, J. E. Saros, E. J. Hundey, S. Sadro, J. Kopáček, R. Sommaruga, M. J. Kainz, A. L. Strecker, S. Chandra, D. M. Walters, D. L. Preston, N. Michelutti, F. Lepori, S. A. Spaulding, K. R. Christianson, J. M. Melack, and J. P. Smol. 2019. Mountain lakes: Eyes on global environmental change. Global and Planetary Change 178:77–95.

Mouquet, N., and M. Loreau. 2003. Community patterns in source-sink metacommunities. The American naturalist 162:544–557.

Naselli-Flores, L., and J. Padisák. 2016. Blowing in the wind: how many roads can a phytoplanktont walk down? A synthesis on phytoplankton biogeography and spatial processes. Hydrobiologia 764:303–313.

Newman, E. A. 2019. Disturbance Ecology in the Anthropocene. Frontiers in Ecology and Evolution 7.

Newman, M. E. J. 2003. The structure and function of complex networks:48.

O’Connor, M. I., D. R. Barneche, A. L. González, and J. Messier. 2020. Editorial: Unifying Ecology Across Scales: Progress, Challenges and Opportunities. Frontiers in Ecology and Evolution 8.

Oksanen, J., F. Guillaume, M. F. Blanchet, K. Roeland, P. Legendre, D. McGlinn, P. R. Minchin, R. B. O’Hara, G. L. Simpson, P. Solymos, M. H. H. Stevens, E. Szoecs, and H. Wagner. 2010. Vegan: community ecology package.

O’Malley, M. A. 2008. ‘Everything is everywhere: but the environment selects’: ubiquitous distribution and ecological determinism in microbial biogeography. Studies in History and Philosophy of Science Part C: Studies in History and Philosophy of Biological and Biomedical Sciences 39:314– 325.

Onandia, G., S. Maassen, C. L. Musseau, S. A. Berger, C. Olmo, J. M. Jeschke, and G. Lischeid. 2021. Key drivers structuring rotifer communities in ponds: Insights into an agricultural landscape. Journal of Plankton Research 43:396–412.

Ortiz, E., A. I. Borthagaray, R. Ramos-Jiliberto, and M. Arim. 2023. Scaling of biological rates with body size as a backbone in the assembly of metacommunity biodiversity. Biology Letters.

Padisák, J., G. Vasas, and G. Borics. 2016. Phycogeography of freshwater phytoplankton: traditional knowledge and new molecular tools. Hydrobiologia 764:3–27.

Pardini, R., E. Nichols, and T. Püttker. 2018. Biodiversity Response to Habitat Loss and Fragmentation. Pages 229–239 Encyclopedia of the Anthropocene. Elsevier.

Pekel, J. F., A. Cottam, N. Gorelick, and A. S. Belward. 2016. High-resolution mapping of global surface water and its long-term changes. Nature 540:418–422.

Peres-Neto, P. R., P. Legendre, S. Dray, and D. Borcard. 2006. Variation Partitioning of Species Data Matrices: Estimation and Comparison of Fractions. Ecology 87:2614–2625.

Perez Rocha, M., L. M. Bini, S. Domisch, K. T. Tolonen, J. Jyrkänkallio-Mikkola, J. Soininen, J. Hjort, and J. Heino. 2018. Local environment and space drive multiple facets of stream macroinvertebrate beta diversity. Journal of Biogeography 45:2744–2754.

Pinceel, T., L. Brendonck, and B. Vanschoenwinkel. 2016. Propagule size and shape may promote local wind dispersal in freshwater zooplankton-a wind tunnel experiment. Limnology and Oceanography 61:122–131.

Podani, J., and D. Schmera. 2011. A new conceptual and methodological framework for exploring and explaining pattern in presence-absence data. Oikos 120:1625–1638.

Radinger, J., and C. Wolter. 2014. Patterns and predictors of fish dispersal in rivers. Fish and Fisheries 15:456–473.

Reche, I., E. Pulido-Villena, R. Morales-Baquero, and E. O. Casamayor. 2005. Does Ecosystem Size Determine Aquatic Bacterial Richness? Ecology 86:1715–1722.

Robinson, C. T., and B. Kawecka. 2005. Benthic diatoms of an Alpine stream/lake network in Switzerland. Aquatic Sciences 67:492–506.

Rocha, B. da S., C. A. de Souza, K. B. Machado, L. C. G. Vieira, and J. C. Nabout. 2020. The relative influence of the environment, land use, and space on the functional and taxonomic structures of phytoplankton and zooplankton metacommunities in tropical reservoirs. Freshwater Science 39:321–333.

Rognes, T., T. Flouri, B. Nichols, C. Quince, and F. Mahé. 2016. VSEARCH: a versatile open source tool for metagenomics. PeerJ 4:e2584.

Sarremejane, R., H. Mykrä, N. Bonada, J. Aroviita, and T. Muotka. 2017. Habitat connectivity and dispersal ability drive the assembly mechanisms of macroinvertebrate communities in river networks. Freshwater Biology 62:1073–1082.

Savary, P., J.-P. Lessard, and P. R. Peres-Neto. 2023. Heterogeneous dispersal networks to improve biodiversity science. Trends in Ecology & Evolution.

Schindler, D. W. 2009. Lakes as sentinels and integrators for the effects of climate change on watersheds, airsheds, and landscapes. Limnology and Oceanography 54:2349–2358.

Schmera, D., D. Árva, P. Boda, E. Bódis, Á. Bolgovics, G. Borics, A. Csercsa, C. Deák, E. Krasznai, B. A. Lukács, P. Mauchart, A. Móra, P. Sály, A. Specziár, K. Süveges, I. Szivák, P. Takács, M. Tóth, G. Várbíró, A. E. Vojtkó, and T. Erős. 2018. Does isolation influence the relative role of environmental and dispersal-related processes in stream networks? An empirical test of the network position hypothesis using multiple taxa. Freshwater Biology 63:74–85.

Seymour, M., E. A. Fronhofer, and F. Altermatt. 2015. Dendritic network structure and dispersal affect temporal dynamics of diversity and species persistence. Oikos 124:908–916.

Solé, R. 2012. Phase transitions (Primers in complex systems). Princeton University Press.

Stomp, M., J. Huisman, G. G. Mittelbach, E. Litchman, and C. A. Klausmeier. 2011. Large-scale biodiversity patterns in freshwater phytoplankton. Ecology 92:2096–2107.

Suzuki, Y., and E. P. Economo. 2021. From species sorting to mass effects: spatial network structure mediates the shift between metacommunity archetypes. Ecography 44:715–726.

Thompson, P. L., L. M. Guzman, L. De Meester, Z. Horváth, R. Ptacnik, B. Vanschoenwinkel, D. S. Viana, and J. M. Chase. 2020. A process-based metacommunity framework linking local and regional scale community ecology. Ecology Letters 23:1314–1329.

Thompson, P. L., B. Rayfield, and A. Gonzalez. 2017. Loss of habitat and connectivity erodes species diversity, ecosystem functioning, and stability in metacommunity networks. Ecography 40:98– 108.

Tonkin, J. D., F. Altermatt, D. S. Finn, J. Heino, J. D. Olden, S. U. Pauls, and D. A. Lytle. 2018a. The role of dispersal in river network metacommunities: Patterns, processes, and pathways. Freshwater Biology 63:141–163.

Tonkin, J. D., J. Heino, and F. Altermatt. 2018b. Metacommunities in river networks: The importance of network structure and connectivity on patterns and processes. Freshwater Biology 63:1–5.

Tonkin, J. D., J. Heino, A. Sundermann, P. Haase, and S. C. Jähnig. 2016a. Context dependency in biodiversity patterns of central German stream metacommunities. Freshwater Biology 61:607– 620.

Tonkin, J. D., S. Stoll, S. C. Jähnig, and P. Haase. 2016b. Contrasting metacommunity structure and beta diversity in an aquatic-floodplain system. Oikos 125:686–697.

Turner, M. G. 2005. Landscape Ecology: What Is the State of the Science? Annual Review of Ecology, Evolution, and Systematics 36:319–344.

Turner, M. G., and R. H. Gardner. 2015. Landscape Ecology in Theory and Practice. Springer, New York.

Tytgat, B., E. Verleyen, M. Sweetlove, K. Van den Berge, E. Pinseel, D. A. Hodgson, S. L. Chown, K. Sabbe, A. Wilmotte, A. Willems, The Polar Lake Sampling Consortium, and W. Vyverman. 2023. Polar lake microbiomes have distinct evolutionary histories. Science Advances 9:eade7130.

Urban, D. L., and T. H. Keitt. 2001. Landscape connectivity: A graph-theoretic perspective. Ecology 85:1205–1218.

Utermöhl, H. 1958. Zur Vervollkommnung der quantitativen Phytoplankton-Methodik. Internationale Vereinigung für Theoretische und Angewandte Limnologie: Mitteilungen 9:1–38.

Vannette, R. L., and T. Fukami. 2017. Dispersal enhances beta diversity in nectar microbes. Ecology Letters 20:901–910.

Vanschoenwinkel, B., S. Gielen, M. Seaman, and L. Brendonck. 2008. Any way the wind blows - frequent wind dispersal drives species sorting in ephemeral aquatic communities. Oikos 117:125–134.

Vellend. 2016. The Theory of Ecological Communities. Princeton University Press.

Whitfield, J. 2005. Is Everything Everywhere? Science 310:960–961.

Winter, C., B. Matthews, and C. A. Suttle. 2013. Effects of environmental variation and spatial distance on Bacteria, Archaea and viruses in sub-polar and arctic waters. The ISME Journal 7:1507–1518.

Wood, S. N. 2003. Thin Plate Regression Splines. Journal of the Royal Statistical Society Series B: Statistical Methodology 65:95–114.

Zhang, G., T. Yao, S. Piao, T. Bolch, H. Xie, D. Chen, Y. Gao, C. M. O’Reilly, C. K. Shum, K. Yang, S. Yi, Y. Lei, W. Wang, Y. He, K. Shang, X. Yang, and H. Zhang. 2017. Extensive and drastically different alpine lake changes on Asia’s high plateaus during the past four decades. Geophysical Research Letters 44:252–260.

